# The Linked Selection Signature of Rapid Adaptation in Temporal Genomic Data

**DOI:** 10.1101/559419

**Authors:** Vince Buffalo, Graham Coop

## Abstract

The majority of empirical population genetic studies have tried to understand the evolutionary processes that have shaped genetic variation in a single sample taken from a present-day population. However, genomic data collected over tens of generations in both natural and lab populations are increasingly used to find selected loci underpinning adaptation over these short timescales. Although these studies have been quite successful in detecting selection on large-effect loci, the fitness differences between individuals are often polygenic, such that selection leads to allele frequency changes that are difficult to distinguish from genetic drift. However, one promising signal comes from polygenic selection’s effect on neutral sites that become stochastically associated with the genetic backgrounds that lead to fitness differences between individuals. Previous theoretic work has established that the random associations between a neutral allele and heritable fitness backgrounds act to reduce effective population experienced by this neutral allele (Santiago and Caballero 1995, 1998). These associations perturb neutral allele frequency trajectories, creating autocovariance in the allele frequency changes across generations. Here, we show how temporal genomic data allows us to measure the temporal autocovariance in allele frequency changes and characterize the genome-wide impact of polygenic selection. We develop expressions for these temporal autocovariances, showing their magnitude is determined by the level of additive genetic variation, recombination, and linkage disequilibria in a region. Furthermore, by using analytic expressions for the temporal variances and autocovariances in allele frequency, we demonstrate that one can estimate the additive genetic variation for fitness and the drift-effective population size from temporal genomic data. We also show how the proportion of total variation in allele frequency change due to linked selection can be estimated from temporal data. Overall, we demonstrate temporal genomic data offers opportunities to identify the role of linked selection on genome-wide diversity over short timescales, and can help bridge population genetic and quantitative genetic studies of adaptation.

---

Adaptation can occur over remarkably short, ecological timescales, with dramatic changes in phenotype occurring over just a few generations in natural populations. This rapid pace of adaptive change has been known to be mirrored at the genetic level since the early work of Fisher and Ford (1947) testing whether the rapid decline in a coloration polymorphism was consistent with natural selection or genetic drift. Since then, researchers have continued to use temporal data to detect selection on polymorphisms over short timescales in natural populations (Dobzhansky 1943, 1971; Fisher and Ford 1947; Kettlewell 1958, 1961; Mueller et al. 1985b), as well as quantify the rate of genetic drift (Mueller et al. 1985a; Nei and Tajima 1981; Pollak 1983; Prout 1954; Wallace 1956; Wang and Whitlock 2003; Waples 1989). However, this line of work in sexual populations has been partially eclipsed by a vast body of work examining large-scale population-genetic and -genomic data sets from a single contemporary timepoint. More recently, studies have applied similar temporal approaches to whole-genome data to discover selected loci in contemporaneous natural populations (Bergland et al. 2014; Rajpurohit et al. 2018), evolve and re-sequence studies (Barghi et al. 2019; Burke et al. 2010; Franssen et al. 2017; Johansson et al. 2010; Orozco-terWengel et al. 2012; Teotónio et al. 2009; Turner and Miller 2012; Turner et al. 2011), and ancient DNA (Fu et al. 2016; Mathieson et al. 2015). Furthermore, numerous methods have been developed to estimate effective population size (Nei and Tajima 1981; Pollak 1983; Waples 1989) and detect selected loci (Feder et al. 2014; Malaspinas et al. 2012; Mathieson and McVean 2013; Terhorst et al. 2015) from time-series data.

Overall, these approaches have identified compelling examples where selection has driven extreme allele frequency change at particular loci that is inconsistent with drift alone. However, most adaptation on ecological timescales likely involves selection on phenotypes with polygenic architecture and abundant standing variation (Endler 1986; Hendry and Kinnison 1999; Kinnison and Hendry 2001; Kopp and Hermisson 2009b). We know from theory that adaptation on such traits can result from very subtle allele frequency changes across the many loci that underlie the trait (Bulmer 1980), at least for the short-term evolutionary response (Chevin and Hospital 2008; Hermisson and Pennings 2005; Höllinger et al. 2019; Jain and Stephan 2015, 2017; Thornton 2018). These changes may be individually indistinguishable from genetic drift in temporal data. This poses a challenge for population genetic approaches to quantify selection: rapid phenotypic adaptations occurring on ecological timescales may leave a signal on genome-wide patterns of diversity that is undetectable by methods focused on individual loci.

Here, we explore an alternative: rather than aiming to find selected loci, we can use temporal data to quantify the genome-wide effects of linked selection during polygenic adaptation. Linked selection introduces a new source of stochasticity into evolution, as a neutral allele’s frequency change depends on the fitness of the set of random genetic backgrounds it finds itself on (Gillespie 2000). The impact linked selection has on neutral loci is mediated by associations (linkage disequilibria, or LD) with selected loci and hence their recombination environment; neutral loci tightly linked to selected sites experience greater average reductions in diversity than more loosely coupled sites. Studies using a single timepoint have long exploited this idea, with some of the first evidence of pervasive natural selection being the correlation between diversity and recombination in *Drosophila* (Aguade et al. 1989; Begun and Aquadro 1992). Various forms of linked selection give rise to such patterns with much attention focusing on the hitchhiking (positive selection; Maynard Smith and Haigh 1974), or background selection models (negative, or purifying selection; Charlesworth et al. 1993; Hudson and Kaplan 1995). Recent genomic studies have modeled patterns of genome-wide diversity considering substitutions, functional constraint, and recombination environments to estimate parameters of hitchhiking and background selection models, and have begun to differentiate between these models (Elyashiv et al. 2016; Hernandez et al. 2011; McVicker et al. 2009). Across-taxa comparisons have shown that signals of linked selection are present in many sexual organisms, and that in some species a proportion of the stochastic change in allele frequencies is due to the randomness of linked selection instead of genetic drift (Coop 2016; Corbett-Detig et al. 2015; Cutter and Payseur 2013). Likewise, in asexual and facultatively sexual organisms both theory and empirical work show that linked selection and interference are a primary determinant of the levels of genetic diversity (Good et al. 2017, 2014; Neher and Shraiman 2011; Neher 2013).

In this paper, we extend this well-established approach of quantifying genome-wide selection through its indirect impact on linked neutral sites to temporal genomic data. We show that during rapid polygenic selection, linked selection leaves a signal in temporal genomic data that can readily be differentiated from neutral processes. Specifically, selected alleles perturb the allele frequency trajectories of neighboring neutral loci, increasing the variance of neutral allele frequency change and creating covariance between the neutral allele frequency changes across generations. Earlier work has modeled this effect on neutral alleles as a long-term reduction in the effective population size (J A Woolliams, N R Wray, R Thompson 2008; Robertson 1961; Santiago and Caballero 1995, 1998; Wray and Thompson 1990), but the increasing availability of genome-wide frequency data across multiple timepoints allows us to directly quantify the extent of linked selection over short ecological timescales (tens of generations). We develop theory for the variances and covariances of neutral allele frequency change under selection, and show that analogous to diversity in a single timepoint study, their magnitude depends on the local fitness variation, recombination, and linkage disequilibria environment. Furthermore, we show our theory can (1) directly partition the variation in genome-wide frequency change into the components caused by drift and selection, (2) estimate the level of additive genetic variation for fitness and how it changes over time, and (3) detect patterns of fluctuating selection from temporal data. Overall, we believe our approach to modeling temporal genomic data will provide a more complete picture of how selection shapes allele frequency changes over ecological timescales in natural populations, potentially allowing us to understand short-term effects of linked selection that would otherwise not be perceptible from studies using a single timepoint.

## 1 Outline of Temporal Autocovariance Theory

Our goal is to understand how linked selection affects the frequency trajectories of neutral sites by modeling the variances and covariances of neutral allele frequency changes (Δ*p*_*t*_ = *p*_*t*+1_ − *p*_*t*_, where *p*_*t*_ is the population frequency at time *t*). We assume a closed population with discrete, non-overlapping generations. When there are no heritable fitness differences between individuals, genetic drift is the only source of stochasticity of allele frequency change due to two sources of variation: random non-heritable, or environmental differences in offspring number, and Mendelian segregation of heterozygotes. Both are directionless such that when averaged over evolutionary replicates, the expected change in allele frequency due to drift alone is 𝔼(Δ*p*_*t*_) = 0 and quantified by the variance in allele frequency change Var(Δ*p*_*t*_) (as quantified by the variance effective population size Charlesworth 2009; Crow and Kimura 1970; Wright 1938).

When there are heritable fitness differences between individuals in the population, a third source of stochasticity affects a neutral allele’s frequency change: neutral alleles can become randomly associated with the genetic backgrounds that determine the fitness differences between individuals (Robertson 1961; Santiago and Caballero 1995, 1998). Even though the neutral alleles do not impact fitness, their frequency trajectories are perturbed by their fitness background, as those on advantage backgrounds leave more descendants, while those on disadvantageous backgrounds leave fewer. We can partition a neutral allele frequency’s change into these three uncorrelated stochastic components (following Santiago and Caballero, 1995, see our Appendix Section A.1 for proof),

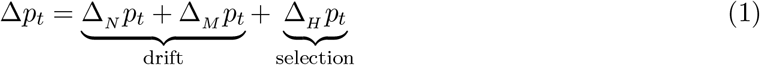

where, Δ_*N*_ *p*_*t*_, Δ_*M*_*p*_*t*_, and Δ_*H*_*p*_*t*_ are the neutral allele’s frequency changes due to non-heritable variation in fitness between diploid individuals, Mendelian segregation of heterozygotes into offspring, and heritable variation in fitness (we refer to this as the *heritable change* in neutral allele frequency), respectively. Note while throughout the paper we consider the allele frequency change between adjacent generations, the same approach can be extended to situations where the study system cannot be observed every generation. Like the stochastic components of drift, the allele frequency change due to heritable fitness differences is directionless (𝔼(Δ_*H*_*p*_*t*_) = 0). Additionally, since each component is uncorrelated with the others, the variance in allele frequency change is

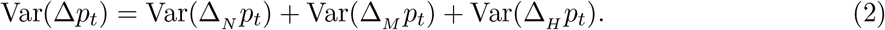

The terms Var(Δ_*N*_*p*_*t*_) and Var(Δ_*M*_*p*_*t*_) capture the variance due to the random reproduction process, and the former can accommodate extra non-heritable variance in offspring number (as long as individuals are exchangeable with respect to their genotype; Cannings 1974), while the term Var(Δ_*H*_*p*_*t*_) captures heritable fitness variation due to systematic differences in the fitness of individuals caused by their genotypes.

In addition to inflating the within-generation variance in allele frequency change, heritable fitness variation has another profound affect on neutral alleles: while the stochastic components of drift have independent effects on frequency change each generation, heritable variation in fitness creates *temporal autocovariance* in neutral allele frequency changes across generations. The contribution of temporal autocovariance is evident by writing the total cumulative allele frequency change as the sum of allele frequency changes each generation,

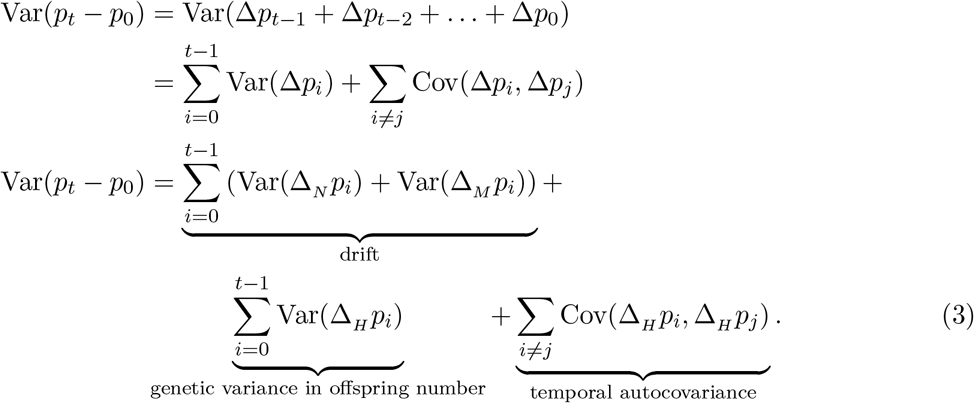

These covariance terms are expected to be non-zero only when there is heritable variation in fitness (assuming there is neither non-Mendelian segregation nor covariance between parental and offspring environment, e.g. as in Heyer et al. 2005).

Temporal autocovariance is caused by the persistence over generations of the statistical associations (linkage disequilibria) between a neutral allele and the fitnesses of the random genetic backgrounds it finds itself on; as long as some fraction of associations persist, the heritable variation for fitness in one generation is predictive of the change in later generations, as illustrated by the fact that Cov(Δ*p*_2_, Δ*p*_0_) > 0 (see Figure 1A). Ultimately segregation and recombination break down haplotypes and shuffle alleles among chromosomes, leading to the decay of autocovariance with time.

**Figure 1:**
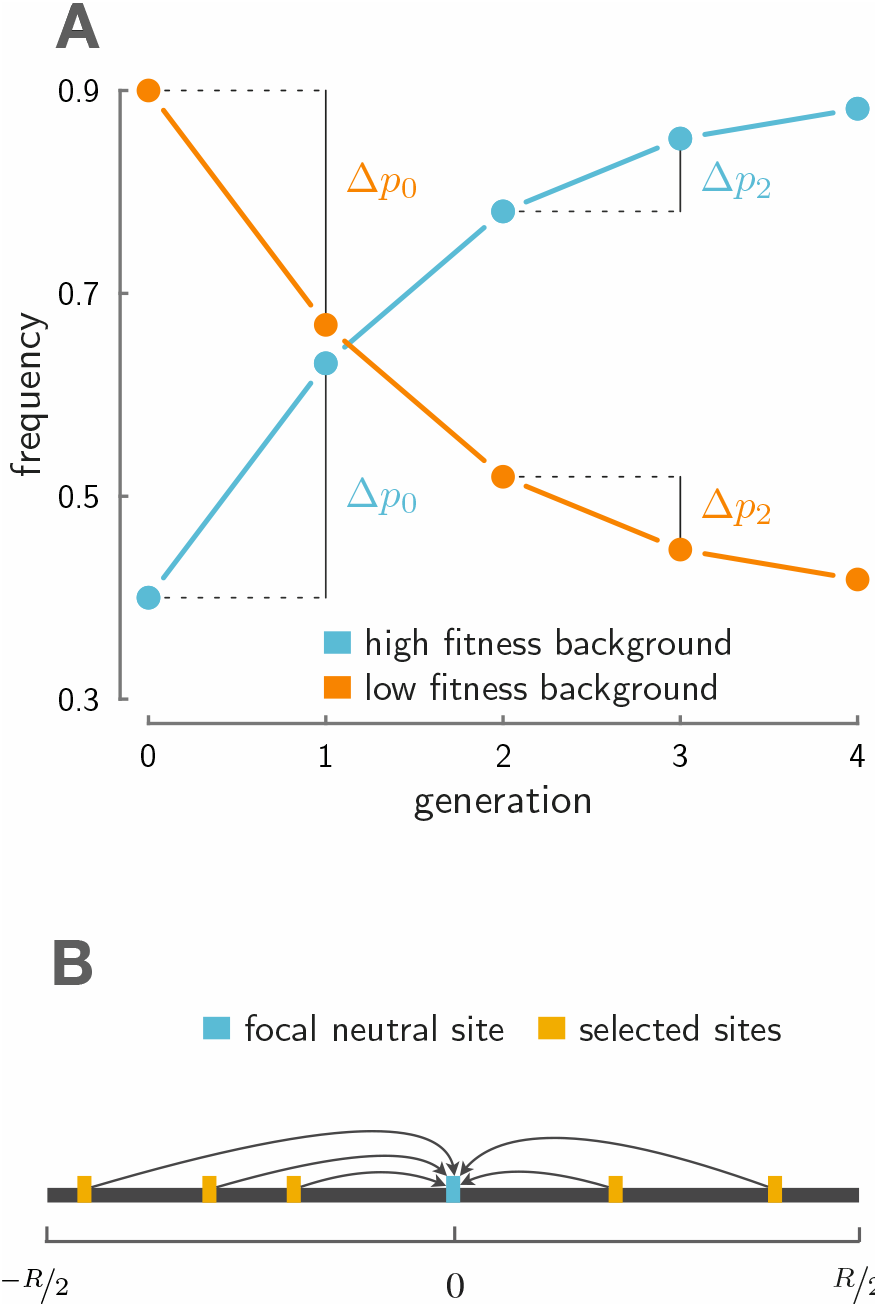
A: On an advantageous background (light blue), a neutral allele increases in frequency leading to a positive change in allele frequency early on, Δ*p*_0_ = *p*_1_ − *p*_0_. As long as some fraction of neutral alleles remain associated with this advantageous background, the neutral allele is expected to increase in frequency in later generations, here, Δ*p*_2_ = *p*_3_ − *p*_2_. This creates temporal autocovariance, Cov(Δ*p*_2_, Δ*p*_0_) > 0. Similarly, had the neutral allele found itself on a low-fitness background (orange), this would also create temporal autocovariance. B: This depicts the setup for our multilocus model. Multiple alleles (yellow) determine the fitness in a region *R* Morgans in length, and these perturb the allele frequency trajectory of a focal neutral site (light blue).

The effect heritable variation has on neutral alleles has traditionally been modeled in a quantitative genetics framework where a large number of loosely linked polymorphisms contribute to heritable fitness differences between individuals, and the impact of heritable fitness variation on a neutral allele is quantified as a reduction in its long-run effective population size (Robertson 1961; Santiago and Caballero 1995, 1998). This form of linked selection can be contrasted with classic population genetic hitchhiking theory (Maynard Smith and Haigh 1974), which considers how neutral alleles closely linked to a new beneficial mutation are effected as it sweeps to fixation. While classic population genetic linked selection models consider how neutral variation is affected by strong associations caused by tight linkage to an advantageous site, quantitative genetic models of linked selection considered the weakest forms of associations: those between unlinked loci within an individual (Morley 1954; Robertson 1961; Santiago and Caballero 1995; see Barton 2000 for more on the two models of linked selection). These distant associations are quickly established but are rapidly broken down by segregation and independent assortment, yet still can have a marked effect on diversity (Robertson 1961; Santiago and Caballero 1995). However, since the impact of heritable fitness variation has traditionally been modeled as causing a reduction in the effective population size, there has been no direct way to separately estimate its effects from those of drift. We show that with temporal genomic data, one can directly measure the levels of temporal variances and autocovariances of allele frequency change in the population. Additionally, we show temporal autocovariance is created under both tight and loose linked selection, and below, develop expressions for its magnitude that are applicable to both, bridging the two regimes of linked selection.

## 2 A model for multilocus temporal autocovariance

Here, we develop theory for the temporal autocovariance in a neutral allele’s frequency changes through time, generated by the presence of heritable fitness in the population. We measure the temporal autocovariance Cov(Δ*p*_*t*_, Δ*p*_*s*_) at a single diallelic neutral locus. Since only allele frequency changes due to heritable variation in *fitness* contribute to temporal autocovariance, we can focus exclusively on the behavior of Δ_*H*_*p*_*t*_ in deriving our expressions for the autocovariance across timepoints. We imagine that an individual *i* has fitness *f*_*i*_, i.e. that their expected number of children is *f*_*i*_. We assume a constant population size, and so the population average fitness 𝔼_*i*_(*f*_*i*_) = 1. Additionally, we assume that all fitness variation has an additive, polygenic architecture. Then, with *L* loci contributing to fitness, we can write individual *i*’s fitness as 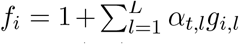, where *α*_*t,l*_ is the effect size in generation *t* and *g*_*i,l*_ ∈ {0, 1, 2} is individual *i*’s gene content at locus *l*. Here, each *α*_*t,l*_ is analogous to a selection coefficient acting at locus *l*, since the fitnesses for genotypes *A*_1_*A*_1_, *A*_1_*A*_2_, and *A*_2_*A*_2_ are 1, 1 + *α*_*t,l*_, and 1 + 2*α*_*t,l*_ respectively. This formulation is approximately equivalent to exponential directional selection on some additively determined trait, implying that selection does not create linkage disequilibria between unlinked loci; see Appendix A.2 for more detail.

When fitness variation exists in the population (that is, Var_*i*_(*f*_*i*_) > 0), the frequency of the neutral allele changes stochastically, as more fit individuals leave more descendents that inherit the neutral allele they carry and less fit individuals leave fewer. Across the population, stochastic associations can form between the genetic components of an individual’s fitness and the neutral allele they carry, leading the neutral allele frequency to change due to fitness differences across individuals. The total heritable change in neutral allele frequency Δ_*H*_*p*_*t*_ can then be partitioned into each individual’s contribution to this change based on their fitness *f*_*i*_ and the number of the tracked neutral alleles they carry, *x*_*i*_ ∈ {0, 1, 2}, giving us

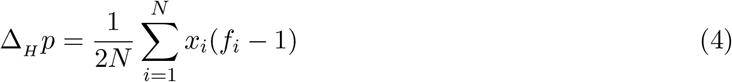

(Santiago and Caballero 1995). Substituting each fitness *f*_*i*_ with its genetic basis and simplifying (see Appendix Section A.2 for details) gives,

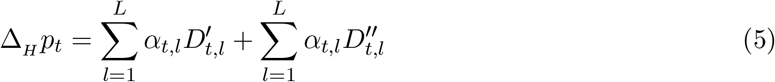

where 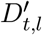 is the *gametic* linkage disequilibrium between the neutral allele and the allele at the selected site *l* on the same gamete, whereas 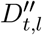 is the *non-gametic* linkage disequilibria, or the covariance *across* the neutral and selected allele on the two different gametes forming an individual (see Weir 1996, p. 121 for details and Appendix Figure 8A for an illustration). Intuitively, this expression tells us that the heritable change in neutral allele frequency is determined by the gametic and non-gametic linkage disequilibrium between the neutral site and all sites that affect an individual’s fitness, scaled by the magnitude of each selected locus’s effect. Alternatively, we can see this as the multivariate breeder’s equation, where the neutral allele is a correlated trait responding to selection on other traits/loci (Lande 1979). This expression is the multilocus analog of the change in a neutral site’s frequency due to hitchhiking at a single linked site (see e.g. equations (2) and (3) in Stephan et al. 2006).

Because the effects of the non-gametic LD are relatively weak compared to the gametic LD for tightly linked loci (see Appendix Section A.5 for an expression of their strength), we ignore these, and hereafter omit the primes in our notation so that *D*_*t,l*_ refers to 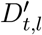. Since 𝔼(Δ_*H*_*p*_*t*_) = 0, we can write the covariance Cov(Δ_*H*_p_*t*_, Δ_*H*_*p*_*s*_) as 𝔼(Δ_*H*_*p*_*t*_Δ_*H*_*p*_*s*_). Hereafter, we also omit the subscript *H* since Cov(Δ*p*_*t*_, Δ*p*_*s*_) = Cov(Δ_*H*_*p*_*t*_, Δ_*H*_*p*_*s*_). Expanding these terms, the covariance between the allele frequency changes at generations t and s can be written as

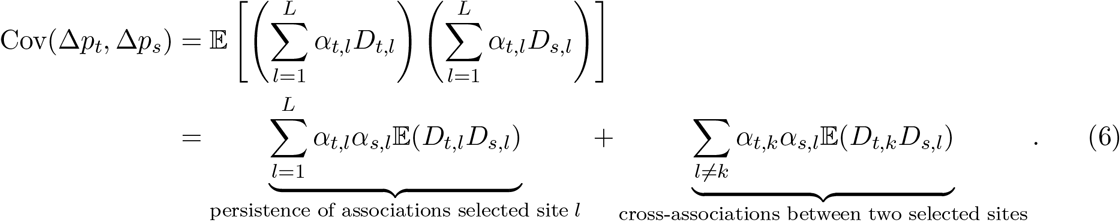

This statement for temporal autocovariance is fairly general, as it can handle fluctuating selection (e.g. when *α*_*t,l*_ varies with time *t*) and any additive multilocus evolution (as long the linkage disequilibria dynamics can be specified). Looking at the first term in this sum, we see the temporal autocovariance is determined in part by the terms 𝔼(*D*_*t,l*_*D*_*s,l*_). These expected LD products reflect the degree to which the association between the neutral locus and a selected site persists from generation *t* to *s* (here, *t* < *s*). Intuitively, the higher the initial association between the neutral and selected loci, and the slower rate of decay of LD between sites, the greater temporal autocovariance will be.

### 2.1 The multilocus temporal autocovariance model with directional selection

Thus far, in reaching Equation (6) we assume only that fitness is additive across loci. In this section, we develop a model of how temporal autocovariance behaves specifically under directional selection beginning at a specific time. We make three assumptions to simplify our expressions. First, we assume that the effect size remains constant through time, such that *α*_*l*_ ≔ *α*_*t,l*_ for all *t* (we relax this assumption in Section 3.2). Second, we ignore the contribution of the second term of Equation (6), 𝔼(*D*_*t,k*_*D*_*s,l*_) (for *k* ≠ *l*), to temporal autocovariance. Under the case where the population is initially at mutation-drift-recombination equilibrium, we expect this product to be zero as there is no directional association between the two selected sites and the neutral site. However, we note that interaction between selected sites (Hill-Robertson interference) will cause this term to become negative (Barton and Otto 2005), a point we return to later. Third, we assume that the selected sites increase in frequency independently, such that the dynamics of the linkage disequilibria between the neutral and selected sites pairs can be modeled using two-locus dynamics. Using a deterministic continuous-time model for the dynamics of the linkage disequilibrium between the selected and neutral site (Barton 2000; Maynard Smith and Haigh 1974), we rewrite the 𝔼(*D*_*t,l*_*D*_*s,l*_) terms in the expression for temporal autocovariance as

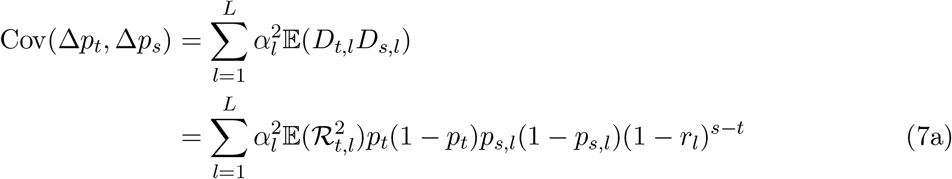

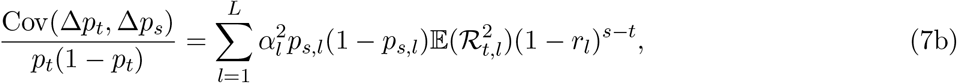

where 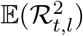 is the square of the correlation between the neutral site and selected site *l* at time *t* (a common measure of LD; Hill and Robertson 1968), and *r*_*l*_ is the recombination fraction between the neutral site and selected site *l*.

We can further simplify this expression by assuming that there is no covariation between the additive genic variation at a selected site and the linkage disequilibrium between that selected site and the neutral site (see Appendix Section A.2 for more detail). This allows us to factor out the average additive genic variation for fitness at time s and write the covariance as

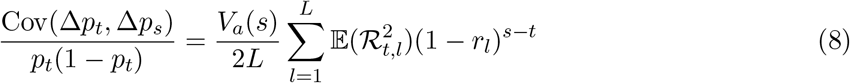

where *V*_*a*_(*s*) is the additive genic variance for fitness, which is the additive genetic variance for fitness (*V*_*A*_) without the contribution of linkage disequilibria between selected sites, 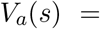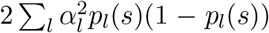. In part, our expression not relying on the linkage disequilibria between selected sites is a result of ignoring the second term in Equation (6); we revisit the consequences of this assumption further on in Section 2.5.

This expression allows us to calculate the temporal autocovariance in cases where we know the vector of recombination fractions between the neutral and each of the selected sites, *r*_1_, *r*_2_, …, *r*_*L*_. Often we do not know the exact positions of these sites, but we can treat these positions as randomly placed on a chromosome and further simplify our model to understand the factors that determine temporal autocovariance.

### 2.2 Temporal autocovariance for an average neutral polymorphism

In the second part of our derivation, we develop a simple intuitive model of how temporal autocovariance is determined by a few key parameters when we make two additional assumptions. First, we assume selected sites are randomly and uniformly distributed along the chromosome, such that a site’s position on the genetic map is a random variable *g* ~ *U*(−*R*/2, *R*/2) (where *R* is the region’s length in Morgans), and the focal neutral site with which we calculate temporal autocovariance lies in the middle of this idealized chromosome at the origin (as depicted in Figure 1B). Then, the recombination fraction between the focal neutral site and a selected site at random position *g* is given by the mapping function *r*(*g*), which maps the position g to a recombination fraction. A simple choice for *r*(*g*) is Haldane’s mapping function, 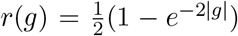, (Haldane 1919; note we take the absolute value of *g* to translate the position *g* to a distance to the focal neutral site) and we use that here. Second, we assume the linkage disequilibrium between each selected site and the focal neutral site depends only the recombination fraction *r*(*g*) between the two loci, and not their absolute positions or effect sizes; then, we rewrite 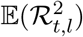 as the function 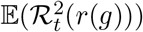. For example, if the population was initially at drift-recombination balance, this would be 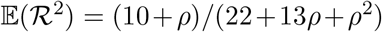 where *ρ* = 4*Nr*(*g*) (Hill and Robertson 1968; Ohta and Kimura 1969). These assumptions allow us to conceptually understand the factors that determine temporal autocovariance; in practice, in temporal studies with linkage disequilibria data and recombination maps, one can directly calculate the sum in Equation (8) (see Appendix Section A.4). We then write the temporal autocovariance experienced by a neutral allele in a region *R* Morgans long containing *V*_*a*_(*s*) fitness variation at time *s* as

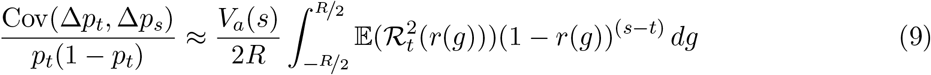

(see Appendix Section A.2 for details).

This integral is the sum of the initial LD between a typical neutral locus and selected site, weighted by the decay of LD due to recombination over *s* − *t* generations. Selection enters here through the total additive genic variance for fitness for the region divided by the genetic map length of the region (*V*_*a*_(*s*)/*R*). Thus a key compound parameter in describing the temporal covariance is the additive genic variance per Morgan, a quantity somewhat similar to the ratio of new adaptive mutations per basepair to recombination per basepair, *ν*_BP_/*r*_BP_, that occurs in models of recurrent sweeps (Stephan et al. 1992) and models of the limits of selection with linked loci (Robertson 1970; Robertson 1976). Note that we could further include the effects of genome-wide fitness variation by adding an additional term encompassing the additive variation residing on other chromosomes, but because the magnitude of these effects is roughly similar to those of the non-gametic LD, we could no longer ignore the 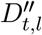 terms as we did after Equation (5).

To validate our theory, we simulate a fixed region of *R* Morgans and calculate the covariance in allele frequency changes by averaging over many uniformly distributed neutral sites within this region. Then, the random distance between a neutral site’s position *n* and a selected site’s position *g* is *c* = |*n* − *g*|, where *n*, *g* ~ *U*(0, *R*); this random variable *c* has a triangle distribution, *f*(*c*) = 2(*R* − *c*)/*R*^2^. Averaging over the positions of both randomly placed neutral and selected sites, the temporal autocovariance is

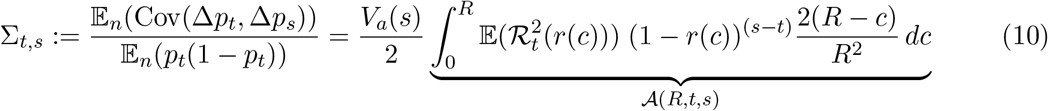

where 𝔼_*n*_(·) indicates an expectation taken over the position of the randomly placed neutral sites, and we define 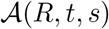 as the average linkage disequilibrium between selected and neutral sites that persists from generations *t* to *s* (*t* ≤ *s*). As is common with estimating the expected values of other ratios like *F*_*ST*_ (Bhatia et al. 2013), we use a ratio of expectations rather than the expectation of the ratio.

We can also use this expression to calculate the variance of allele frequency change. The standardized variance Var(Δ*p*_*t*_)/*p*_*t*_(1−*p*_*t*_) has two components: the drift term and the heritable variance in offspring number. Adding these independent contributions, the standardized variance is

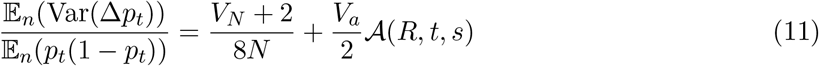

where *V*_*N*_ is the non-heritable variance in offspring number. Under a Wright–Fisher model of reproduction, *V*_*N*_ ≈ 2, this simplifies to

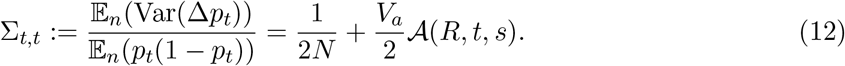

When combined, this expression for the variance in allele frequency change and our expression for temporal autocovariance are in agreement with Robertson (1961) and Santiago and Caballero (1995; 1998) when predicting the total variance in allele frequency change; see Appendix A.6. With the above expressions for the variances and covariances, we have a complete set of theoretic expressions for the variance-covariance matrix of allele frequency change, which we call **Σ**, with the diagonal variance elements Σ_*t,t*_ given by Equation (12), and the upper- and lower-triangle covariance elements Σ_*t,s*_ (*t* ≠ *s*) given by Equation (10).

### 2.3 Modeling the dynamics of additive genic and genetic variation

Our expressions for temporal autocovariance (Equation 10) requires an expression for *V*_*a*_(*t*), the additive genic variation through time. However, we lack general expressions for the dynamics of the additive genic variation during selection, as these dynamics are quite complex for a few reasons. First, since our theory considers polygenic selection at a finite number loci in a region, additive genic variation is not constant as it would be under an infinitesimal model (Bulmer 1980). Second, we allow for arbitrary levels of recombination from very tight linkage to loose linkage. Previous work has shown that predicting the dynamics of additive genic variation in a system with an arbitrary level of recombination is difficult, as both the additive genic and genetic variances depend on the higher-order moments of linkage disequilibrium (see Barton and Turelli 1987, and p. 607 of Turelli and Barton 1990).

A primary determinant of the additive genic variation is the heterozygosity of the selected sites. Assuming effect sizes are constant through time and across loci, we can rewrite the additive genic variation, *V*_*a*_(*t*) = 2*α*^2^∑_*l*_*p*_*l*_(*t*)(1 − *p*_*l*_(*t*)), as

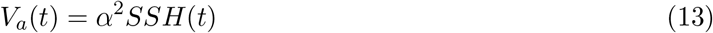

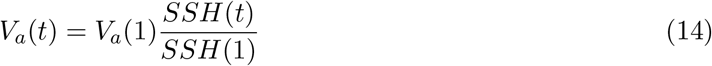

where *SSH*(*t*) = 2∑_*l*_*p*_*l*_(*t*)(1 − *p*_*l*_(*t*)) is the sum of site heterozygosity at time *t*. Ideally, we would directly use *SSH*(*t*) in a region; however, this would require knowing *a priori* which sites are being selected. Instead, we assume that the trait is sufficiently polygenic that frequency changes due to selection are weak, and that the change in heterozygosity at neighboring neutral polymorphic sites approximately mirrors that at selected polymorphisms. Then, using the sum of site heterozygosity at neutral sites, *SSH*_*n*_(*t*), as a proxy for the sum of site heterozygosity at selected sites,

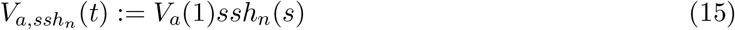

where we define *ssh*_*n*_(*s*) = *SSH*_*n*_(*s*)/*SSH*_*n*_(1) as the factor by which *V*_*a*_(1) decreases at time *s*, approximated by neutral sites’ allele frequency changes. Under this approximation, the dynamics of genic variation are determined by one free parameter, *V*_*a*_(1), and the directly measurable sum of site heterozygosity at neutral sites through time.

Our focus here is on the short-term response of a population, and so we look at the decay of genetic backgrounds present at the onset of directional selection. In reality, new mutations consistently create additive genetic variation for fitness; thus, an equilibrium level of additive genetic variance in the population can be maintained. The long-run effect of linked selection under this equilibrium model is handled by Santiago and Caballero (1995, 1998); see Appendix A.6.

### 2.4 Multilocus Simulation Details

To test our theoretical expressions, we have conducted extensive forward simulations of directional selection on a polygenic trait. We vary four critical parameters in these simulations: (1) the level of additive genetic variance at the onset of selection (*V*_*A*_), (2) the level of recombination (*R* in Morgans), (3) the number of selected sites in the region (*L*), and (4) the population size (*N*). We choose our grid of the selection and recombination parameters based on the levels we would expect across a wide variety of organisms; see Appendix Section A.7 for details. We used three different population sizes (*N* ∈ {100, 500, 1000}), but note that we use *N* = 1000 and a subset of the other parameters in our figures.

Before the onset of selection, we create the initial diploid population from a pool of gametes created by `msprime` (Kelleher et al. 2016), such that the initial allele frequency distribution and linkage disequilibria between sites is at mutation-drift-recombination balance. Details of how `msprime` was called are available on the project’s Github repository, in `R/simpop.r`. Then, we pass this pool of gametes into a forward Wright–Fisher-with-recombination simulation routine and let it evolve for four generations neutrally before initiating selection on the fifth generation. These first four generations of neutral evolution (without mutation) serve as a control to validate that the variance in neutral allele frequency change is as expected under a Wright–Fisher model and that temporal autocovariance between a generation before selection and during selection is zero.

We generate genetic variation for fitness by choosing *L* random loci from the neutrally evolved sites, and randomly assign an effect size of −*α* or +*α*, such that the expected total amount of additive genic variation is *V*_*A*_ (note that the initial additive genic and genetic variance are equal, *V*_*A*_ = *V*_*a*_, as the LD contribution is zero for randomly chosen sites). The details of this are given in Appendix Section A.7. This approach creates some additional variance around the target level of additive genic variation, as the sum of site heterozygosities will vary stochastically across simulation replicates. At the onset of selection, an individual *i*’s trait value is calculated as 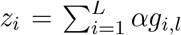 where *g*_*i,l*_ is their number of alleles with effect size *α* at locus *l*. Then, their absolute fitness is calculated using an exponential fitness function 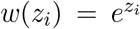 (Turelli and Barton 1990, p. 17). Under our Wright–Fisher model, we sample the parents of the next generation according to a multinomial distribution, where the probability of individual *i* being a parent is 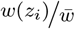.

We record 50 generations of simulated evolution, after which we compute the standardized sample temporal variance-covariance matrix **Q** (this is the sample analog of our theoretic variance-covariance matrix **Σ**), for each replicate as follows. First, we mark frequencies reaching fixation or loss as missing values. This allows the frequency changes before fixation/loss to contribute to the measured covariance, rather than removing the entire locus’s trajectory, which would act to condition the covariance on more intermediate frequencies. Note that one cannot ignore fixations or losses, as these have Δ*p*_*t*_ = 0 and thus an autocovariance of zero, which would act to underestimate the true level of autocovariance at segregating sites. Having marked fixations/losses as missing, we take the frequency matrix and calculate a vector of allele frequency changes 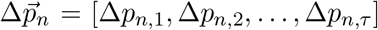 using each neutral locus *n*’s *τ* + 1 observed generations. Finally, we calculate the *τ* × *τ* sample standardized variance-covariance matrix **Q**, averaging over *M* neutral loci such that element *Q*_*t,s*_ is calculated as

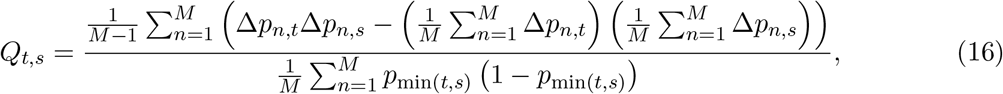

though see Appendix Section A.8 for a bias-corrected version when sample, rather than population allele frequencies are used. Sums over missing values only use pairwise-complete observations, implemented by R’s `cov()` function’s `use=‘pairwise.complete’` argument.

We have extensively validated our simulation procedure in a neutrally evolving population, ensuring that the decay of linkage disequilibrium and the allele frequency change match expectations (see Supplementary Figures 3, 4, and 5).

### 2.5 Comparing theory to simulation results

To validate our expressions for temporal autocovariance, we compare the levels of autocovariance and variance predicted by Equation (10) and (12) to the average levels observed across simulation replicates. To calculate the theoretic values of temporal autocovariance and variance, our expression requires the additive genic variation at *s*, *V*_*a*_(*s*); however, as described in Section 2.3, we lack an analytic expression for the dynamics of genic variation to plug into *V*_*a*_(*s*). Following the approach of others in evolutionary quantitative genetics (Turelli and Barton 1994, p. 930), we substitute the numerical values calculated directly from the simulation data for *V*_*a*_(*s*). Additionally we consider two other numerical values related to the additive genic variance: the observed additive genetic variance from our simulations (*V*_*A*_(*s*) = Var_*i*_(*z*_*i*_) at time *s*, which includes the contribution of LD between selected sites), and the additive genic variation at time s as approximated by the observed decay in the sum of site heterozygosity at neutral sites (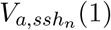, as described in Section 2.3).

Figure 2 compares the fit of our theory with differing additive genetic variances with the empirical covariances from our multilocus simulations. In each panel, we plot the level of temporal autocovariance between the allele frequency change across the first two generations of selection (Δ*p*_5_) and some later allele frequency change Δ*p*_*s*_ where *s* varies along the x-axis. Each point represents the temporal autocovariance (calculated across all sites in a region according to Equation 16) averaged across 100 replicate simulations, with the color of the point indicating the number of selected sites in the region. Within each panel, the temporal autocovariance predicted by Equation (10) is plotted as a set of three lines, one for each of the three different types of variance we have substituted in for *V*_*a*_(*s*). Overall, the fit is close but varies depending on the type of variance used for *V*_*a*_(*s*); we discuss each in turn below.

**Figure 2:**
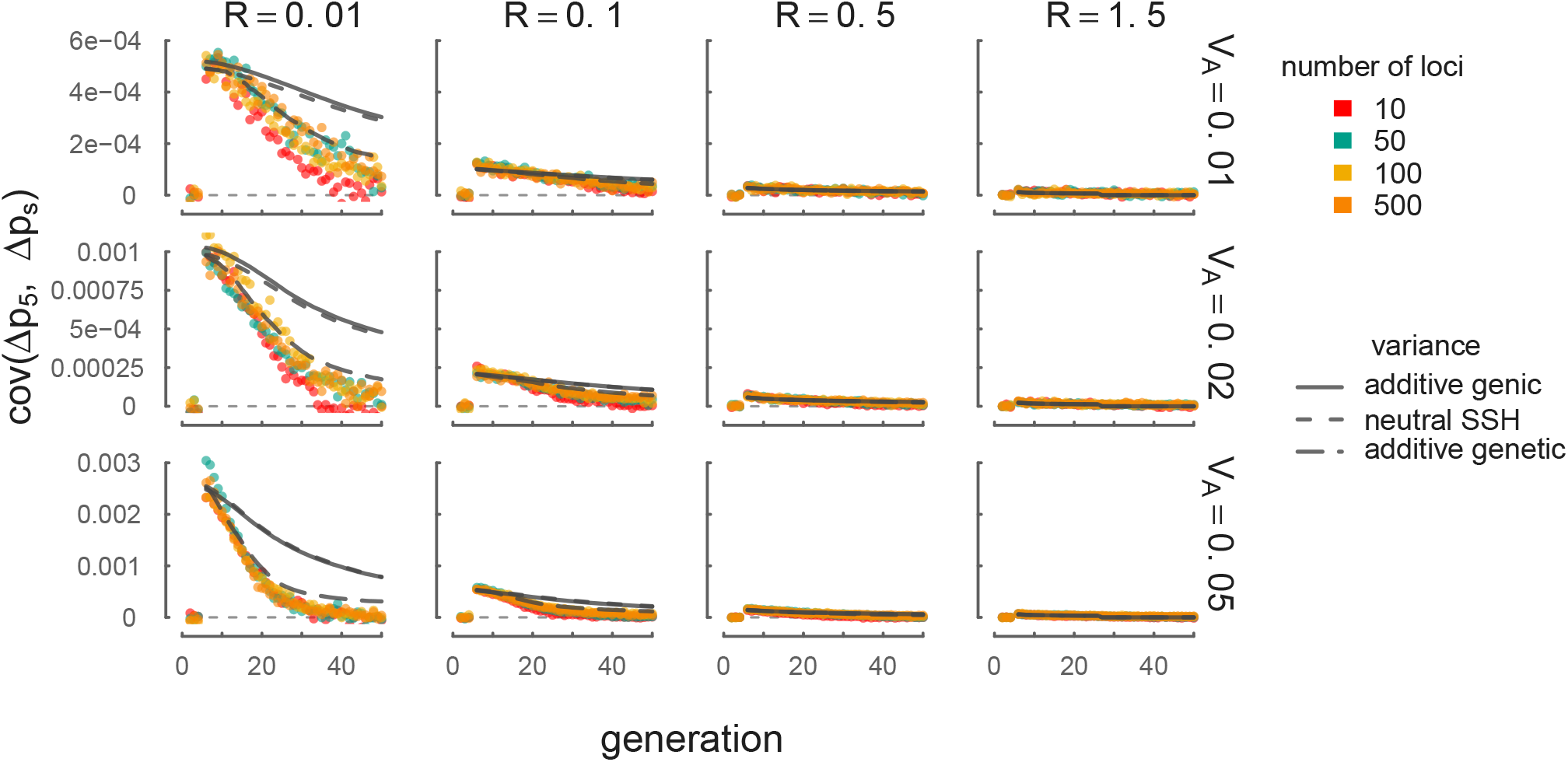
In each panel the temporal autocovariance Cov(Δ*p*_5_, Δ*p*_*s*_) is shown on the y-axis while generation s varies along the x-axis. Selection is initiated on the 5^th^ generation, so Δ*p*_5_ is the neutral allele’s frequency change across the first generation of selection. Each point is the temporal autocovariance between Δ*p*_5_ and the Δ*p*_*s*_ in a region, averaged over 100 simulation replicates, with the colors indicating the number of selected loci. The gray curves indicate the theoretic predictions (for *L* = 500 loci only) using Equation (10), with the equation’s variance provided by the empirically observed additive genic (solid), the additive genetic (long dashes), and the neutral sum of site heterozygosity approximation (short dashes). A thin horizontal dashed line indicates *y* = 0. Across the columns, the level of recombination (in Morgans) is varied; across rows, the initial level of additive genetic variation is varied. Note that while our results here are between the frequency change at the onset of selection Δ*p*_5_ and some later change Δ*p*_*s*_, our covariance theory matches simulation results between any two arbitrary frequency changes Δ*p*_*t*_, and Δ*p*_*s*_; see Supplementary Figure 7.

Using empirical additive genic variation (solid lines), our theory provides a good fit to the simulation results for a short period after selection is initiated (~5 generations) in regions with tighter linkage (*R* = 0.01 Morgans) across a range of additive genetic variation parameters (0.01 ≤ *V*_*A*_ ≤ 0.05; see Supplementary Figure 8 for *V*_*A*_ varying over orders of magnitude). With looser linkage (*R* ≥ 0.1 Morgans), our theory using the empirical additive genic variation fits much more closely over a longer duration (~10-15 generations). Note that some variability is caused by the noise of each simulation replicate around the target initial additive genetic variation *V*_*a*_ (see Section 2.4), as each replicate samples sites from a neutral coalescent. Our theory also accurately predicts the temporal autocovariance for different choices of reference generation, i.e. varying *t*, see Supplementary Figure 7.

When we use the sum of site heterozygosity at neutral sites (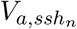, shown as a short-dashed lines) as a proxy for additive genic variation, the theory fits simulations over the same timespan as using the empirical additive genic variation. This is because (1) the SSH at neutral sites closely matches the SSH at selected sites, and (2) both closely follow the dynamics of additive genic variation through time (see Supplementary Figure 6). Using 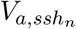 has the advantage that we can directly measure neutral SSH, which proves useful later in Section 3, as we use this approach to help infer the initial additive genic variation at the onset of selection.

Finally, we find using the additive genetic variance *V*_*A*_(*s*) = Var_*i*_(*z*_*i*_) accurately predicts the dynamics of temporal autocovariance over tens of generations (see the long-dashed lines in Figure 2). Furthermore, calculating the temporal autocovariance using the empirical additive genetic variation better fits simulation data in regimes with tight recombination, where using genic variation performs poorly after the first few generations (e.g. the column of panels where *R* = 0.01). Thus using the additive genetic variance in our framework provides a good fit to the temporal dynamics over relatively long time spans.

What differentiates *V*_*A*_(*s*) from *V*_*a*_(*s*) that could explain this better fit? The additive genic variation *V*_*a*_(*s*) ignores the contribution of linkage disequilibria between selected sites. We can write the additive genetic variance as

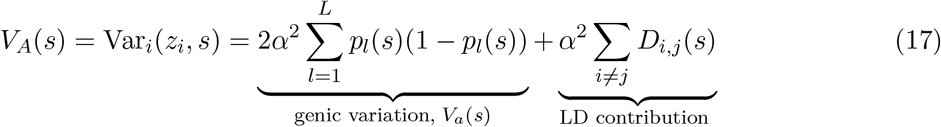

where *D*_*i,j*_(*s*) is the LD between selected sites *i* and *j* at time *s*. At the onset of selection, there is no expected linkage disequilibria between selected sites since the sites and effect sizes were randomly sampled — in other words, 𝔼(*D*_*i,j*_(*s*)) = 0. We see this in Figure 2, as the temporal autocovariance predicted with *V*_*a*_(*s*) match those of *V*_*A*_(*s*) when *s* = 6 (see also Supplementary Figure 6, which plots the empirical additive genic and genetic variances over time). Over time, these two quantities diverge as negative linkage disequilibria build up. While negative linkage disequilibria between selected sites build up due to epistasis under some forms of selection (known as the Bulmer effect, Bulmer 1971; Bulmer 1980), this is known not to happen under multiplicative selection (Bürger 2000, p. 50, 177) that is equivalent to the exponential directional selection fitness surface we have used in our simulations. Instead, the build up of negative linkage disequilibria between selected sites is likely due to Hill-Robertson interference (HRi) between selected sites (Hill and Robertson 1966), which affects the total additive genetic variation that selection is acting on. HRi refers to the creation of negative LD among beneficial alleles in finite population resulting from the fact that beneficial alleles that are on the same haplotype move more quickly through the population than beneficial alleles on deleterious backgrounds, resulting in negative LD. This negative LD among beneficial alleles lowers *V*_*A*_ compared to the genic *V*_*a*_ (Barton and Otto 2005; Crouch 2017; Good et al. 2014; Hill and Robertson 1966). In the derivation of our expression for temporal autocovariance, we greatly simplified the multilocus dynamics by ignoring the second term in Equation (6). This term includes the expected product of two LD terms; each is the LD between the neutral site and a selected site. Using full multilocus theory, one may find that by including these LD products, that *V*_*A*_ rather than *V*_*a*_ factors out the expression in Equation (7), but we leave this for future work. Importantly, our simulation results suggest that the negative linkage disequilibria created by selective interference only affects the temporal autocovariances through the variance term *V*_*a*_(*s*), and that the actual variance determining temporal autocovariance is the additive genetic variance, *V*_*A*_(*s*).

In addition to modeling autocovariance through time, our theory can predict the total temporal variance in allele frequency, Var(*p*_*t*_ − *p*_0_), when there is heritable variation for fitness. Furthermore, from Equation (3) recall that we can decompose Var(*p*_*t*_ − *p*_0_) into variance and covariance components. The variance components are determined by both the magnitude of drift (1/2*N*) and selection according to Equation (11), and the covariance components are determined solely by selection according to Equation (10) (assuming no inheritance of environmental factors). Using our theory, we have predictions for each of these components given the amount of additive genic/genetic variation for fitness, the population size (*N*), and the amount of recombination (*R*). In Figure 3, we compare the magnitudes of these components (averaged over the replicates of our simulations) to our theoretic predictions. We depict the predictions for the variance and covariance components using both the empirical additive genetic variance (*V*_*A*_) and the neutral sum of site heterozygosity proxy 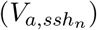 as adjacent bars, each around a point range with the point representing the average value over simulation replicates, and the bars indicating the lower and upper quartiles over simulations.

**Figure 3:**
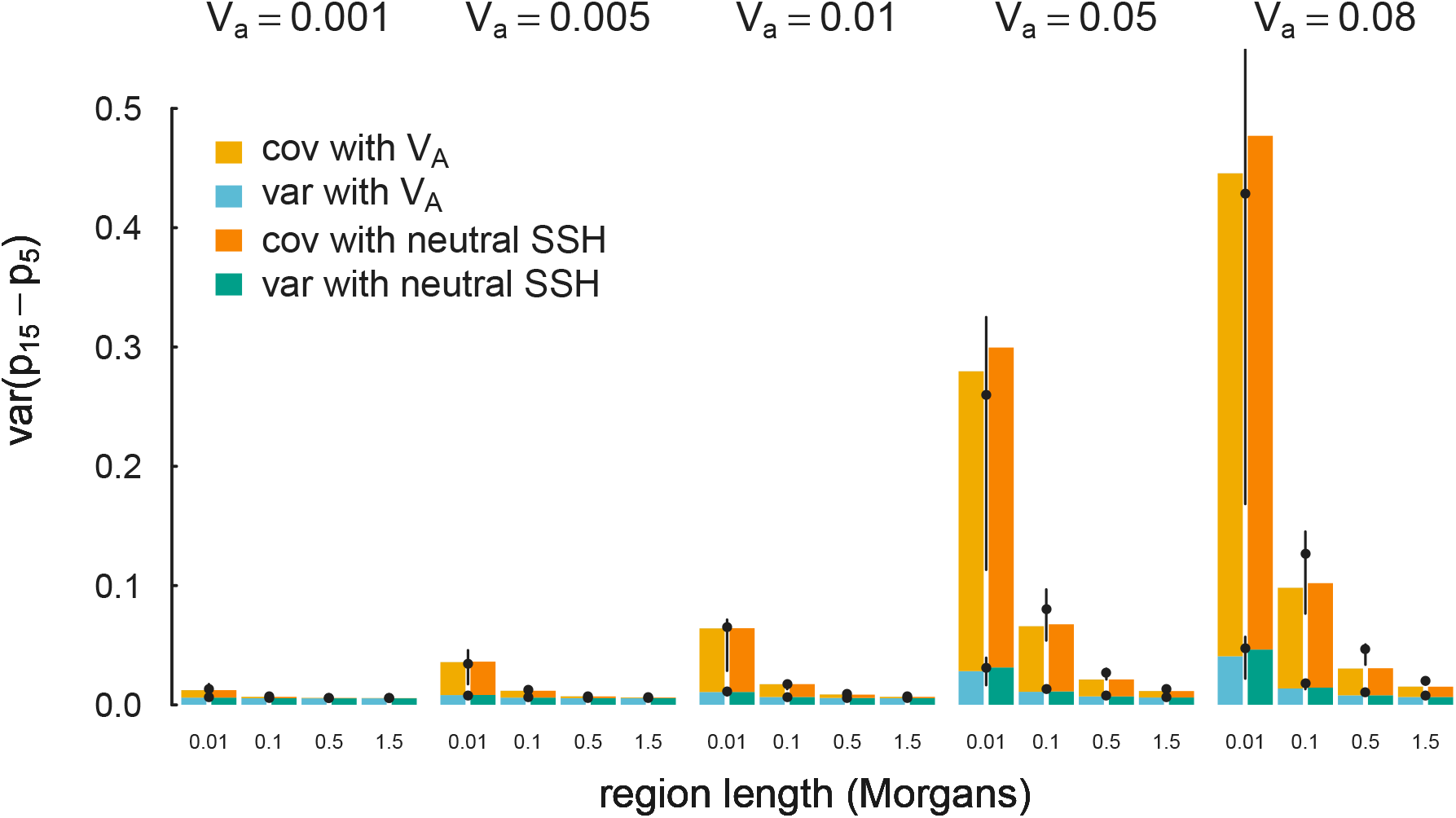
Summing over generations, Equations (6) and (12) accurately predicts the total variation in allele frequency change due to variance and covariance components. The predicted cumulative variance in allele frequency change across the ten generations after selection (Var(*p*_15_ − *p*_5_)) is shown as bars, using both the empirical additive genetic variation *V*_*A*_ (bars to the left of the pointrange), and the empirical neutral sum of site heterozygosity (bars to the right of the pointrange). The variance and covariance components are represented by blue/green and orange/yellow tones respectively. Finally, we show the averaged results of our simulations as pointranges, with the point depicting the average and the bars representing the lower and upper quartiles.

Finally, we have found that across a wide range of recombination and additive genetic variation parameters, the temporal autocovariance Cov(Δ*p*_*t*_, Δ*p*_*s*_) is largely determined by the compound parameter *V*_*A*_/*R* and the number of generations between *t* and *s*, which is a factor in Equation (9). We show in Figure 4 that the temporal autocovariance Cov(Δ*p*_5_, Δ*p*_*s*_) from simulations across a wide range of *V*_*A*_ and *R* parameters fall roughly on the same curve for each number of elapsed *s* − *t* generations.

**Figure 4:**
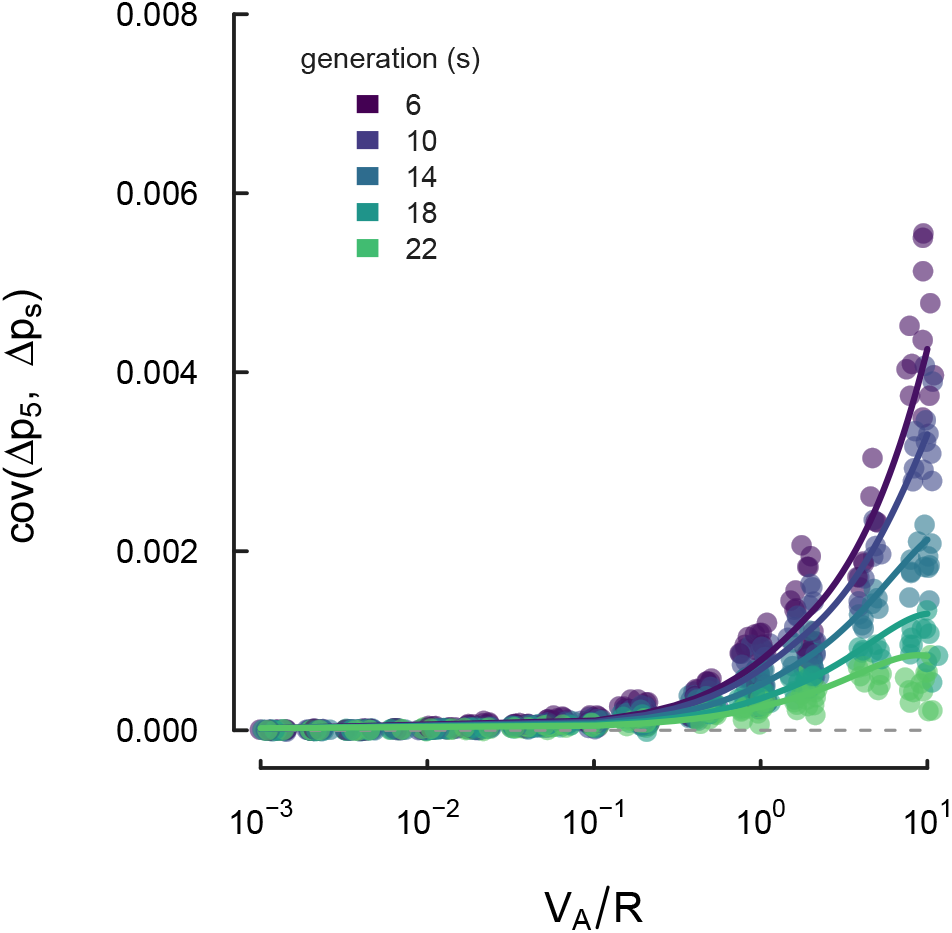
The compound parameter *V*_*A*_/*R* and the number of generations between the temporal autocovariance *s* − *t* largely determines the magnitude of the temporal autocovariance across a wide spectrum of *V*_*A*_ and *R* parameters. Each point is a simulation replicate with its x-axis position given by *V*_*A*_/*R*, the y-axis position equal to the temporal autocovariance, and the number of elapsed generations (*s* − *t*). Each line is a loess curve fit through each set of points for a particular generation (with smoothing parameter *α* = 0.9).

## 3 Estimating linked-selection parameters from temporal autocovariance

Our multilocus theory provides analytic expressions for the expected variances and covariances of a neutral allele’s frequency; thus a natural approach to parameter estimation is to equate these expectations to averages from the data and apply the method-of-moments. We describe a method-of-moments procedure below to estimate the initial additive genetic variance at the onset of selection (*V*_*A*_(1)) in the first generation, and the drift-effective population size (*N*) from temporal data within a single region *R* Morgans long, and then show a simple extension that allows this to be applied to genome-wide data. Our basic approach is to calculate first the sample variances and covariances of the *τ* observed generation-to-generation allele frequency changes, averaging over all of the putatively neutral sites in a region. We then equate these sample variances and covariances to our analytic expressions for the variance and covariances, leaving us with an overdetermined system of equations, which we solve using least squares. We demonstrate this simple estimation procedure provides accurate estimations of initial additive genetic variance and the drift-effective population size. We focus on this procedure, as it is simple and handles incomplete trajectories due to missing data or fixation/loss well. Calculating pairwise-complete covariances can leave sample covariances matrices non-positive definite, which makes maximum likelihood estimation perhaps much more difficult. Throughout, we use population allele frequencies (i.e. there is no sampling noise) which simplifies the description of the method; in Appendix Section A.8 we describe how the method is changed by finite sampling of chromosomes from a population.

From our multilocus theory, we have analytic expressions for each element of the *τ* × *τ* covariance matrix of allele frequency changes in a region. To model the additive genic variance through time, we use the empirical neutral sum of site heterozygosity approximation as described in Section 2.3. This approximates the rate that additive genic variation decreases through time from some initial level, *V*_*A*_(1), which we wish to estimate. In total, we have *τ* + *τ* (*τ* −1)/2 unique moment equations, which for the variance and covariance are defined as

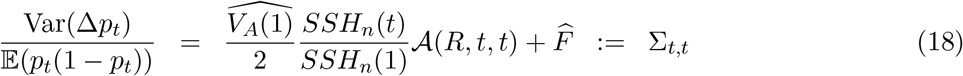

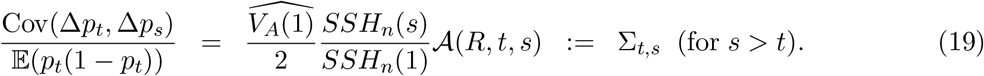

Here the first line gives the form of *τ* equations for the variance of allele frequency changes between subsequent generations, which includes the effect of genetic drift, 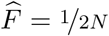. The second line gives the form of the covariances of allele frequency changes among different generations. The term 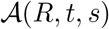 is the average level of linkage disequilibrium after the *s* − *t* generations that have elapsed, given there are *R* Morgans of recombination. In our multilocus theory section and simulations, this is equal to the integral in Equation (10). However, we can also directly calculate a sample 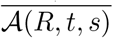 from observed linkage disequilibrium in a region (for details, see Appendix Equation (53)).

Following the method-of-moments, we equate each of these independent *τ* + *τ* (*τ* −1)/2 equations for Σ_*t,s*_ to the observed sampling moments, the elements *Q*_*t,s*_ of the upper triangle of the observed heterozygosity-normalized covariance matrix 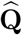 described in Equation (16). This yields *τ* + *τ* (*τ* −1)/2 equations with 2 unknown parameters: 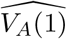 and 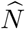. We solve this overdetermined system of equations using least squares, an approach similar to the generalized method-of-moments in econometrics (Hansen 1982). This approach finds parameter estimates that minimize the squared error between the moment-based parameter estimate and the true parameter value, with respect to the true parameter value. We write the elements *Q*_*t,s*_ of the upper triangle of the observed covariance matrix in the vector 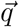, and write the method-of-moments equations as,

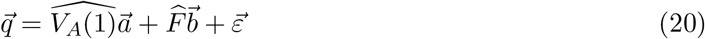

where the elements of 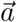 and 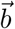, in the same order as 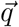, are given by

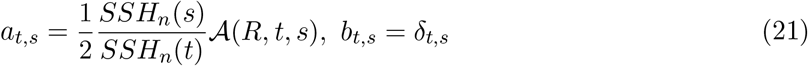

where *δ*_*t,s*_ is an indicator variable that is 1 when *s* = *t* and zero otherwise.

Then, we can readily estimate the parameters 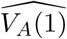 and 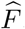 using least squares. We then obtain assume an estimate of 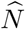 by talking 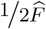. Since these equations are not statistically independent, we cannot assume 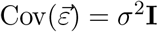. However, this does not affect our estimates 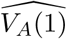 and 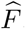, as the least squares procedure is unbiased regardless of the covariance structure between the error terms (Christensen 2011, p. 26).

Using this method-of-moments approach, we sought to infer the parameters of 20 of the replicates across the 254 parameter combinations (the same as used in Figure 2). We use the first five generations after the onset of selection to infer 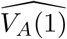 and 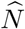, as for this short timespan the additive genetic variance is well approximated using sum of neutral site heterozygosity approach (see Section 2.5). Each simulation replicate includes around 500 neutral sites (the exact number is random, see Appendix A.7 for details).

Applying our approach to these simulations, we find we can infer both the initial level of additive genetic variation *V*_*A*_(1), and the effective population size *N* from multilocus temporal data. In Figure 5A, we show our method-of-moments gives reasonable estimates for the initial level of additive genetic variance over orders of magnitude of additive genetic variation, and different recombination regimes. As additive genetic variation for fitness becomes weaker (the left side of the figure), our estimates become more noisy. In Figure 5B we show the simultaneously estimated population size *N* against the true population size value. We also plot the estimated *N*_*e*_ (not accounting for selection) from a simple temporal estimator, *N*_*e*_ = − *t*/(2 log(1 − *F*)) (Krimbas and Tsakas 1971; Waples 1989) where *F* is Wright’s standardized variance (Wright 1931). While with high *V*_*A*_ and low *R*, the method-of-moments approach still underestimates *N*, it performs far better than a standard temporal *N*_*e*_ estimator that does not account for selection. In Appendix Figure 1, we include a version of this figure calculated using the method of moments on sample allele frequencies for a sample of size *n* = 100 chromosomes.

**Figure 5:**
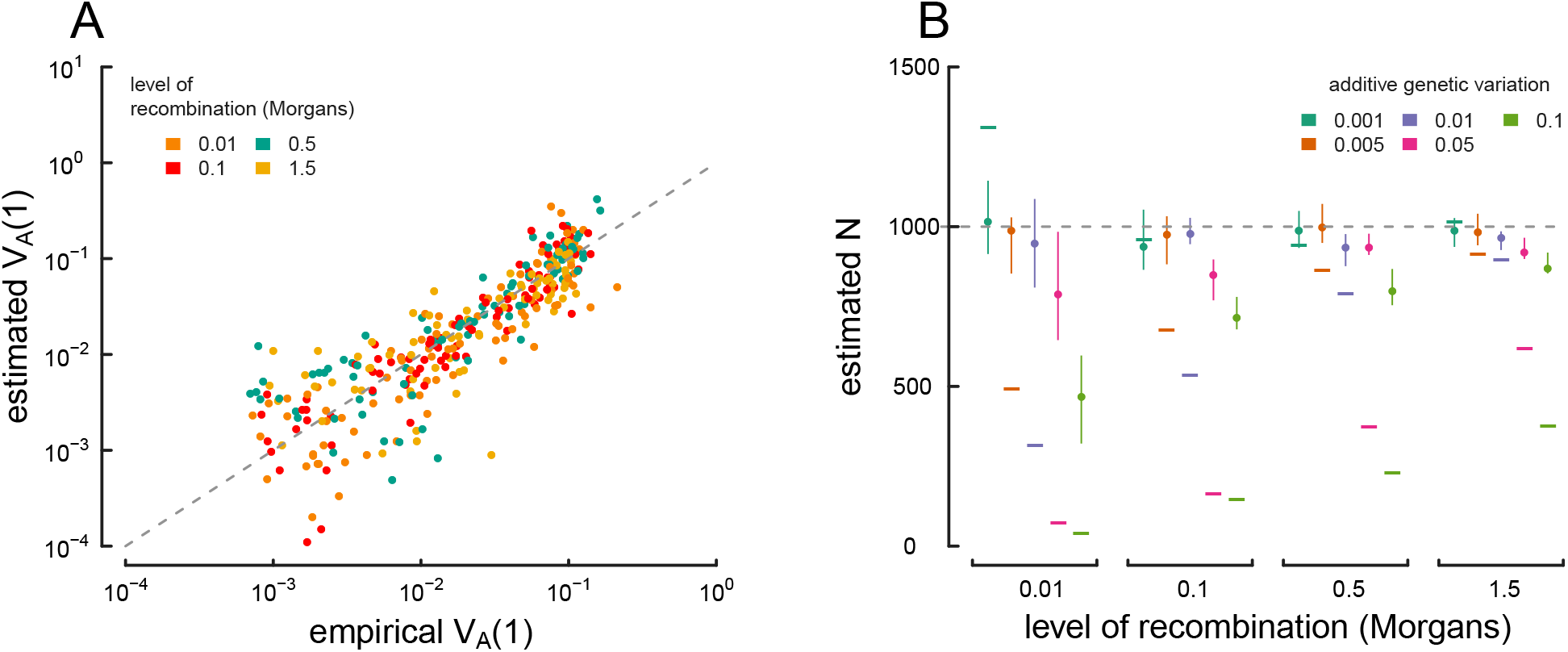
True parameter values and estimates using the method-of-moments approach on multilocus simulation data. (A) The true *V*_*A*_(1) (x-axis) and 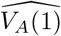 estimated from the variance/covariance matrix (y-axis) for each simulation replicate across different levels of recombination (indicated by each point’s color). The dashed gray line shows the *y* = *x* line where an estimate is exactly true to its real value. Note that the plot is on a log-log scale, as *V*_*A*_ varies across orders of magnitude in our simulations. (B) Estimated drift-effective population size 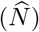 across a range of simulations with different levels of additive genetic variance and recombination. Each point denotes the median, with lines denoting the interquartile range. A simple temporal estimate of the effective population size, estimated with accounting for the effects of selection, is averaged for each replicate and plotted as a dash. The true value (*N* = 1, 000) is shown with the dashed gray line. Population frequencies (without sampling noise) are used in this figure; see Appendix Figure 1 for an analogous figure calculated with sample frequencies.

We can extend this approach to whole-genome data by imagining partitioning the genome into *B* non-overlapping windows of length *w*_BP_ in basepairs (e.g. megabase windows). We first assume that windows contribute uniformly to the genome-wide level of additive genetic variance for fitness, and show how our method-of-moments approach can be used to estimate a global *V*_*A*_(1). Assuming a uniform distribution of genetic variance across basepairs, the total additive variance is *V*_*A*_(1) = *v*_*A*_(1)*Bw*_BP_, across our B windows, where *v*_*A*_(1) is the additive genetic variance per basepair. As each window *i* contributes to *v*_*A*_(1)*w*_BP_, our least squares approach given by Equation (20) becomes

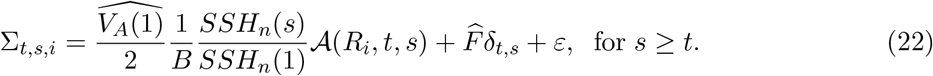

However, we expect *a priori* that windows containing more coding bases might disproportionately contribute to the total additive genetic variance. This suggests an alternative model to fit where partitions of the additive genetic variance across windows are proportional to the number of coding bases, similar to background selection and other linked selection models (CorbettDetig et al. 2015; McVicker et al. 2009; Rockman et al. 2010). Thus, we could write total 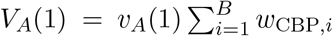 where *w*_CBP*,i*_ is the number of coding or exonic basepairs in window *i* (this could be any quantifiable annotation feature in the window), and 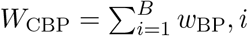 is the total number of coding bases in the genome. With window *i* contributing *v*_*A*_(1)*w*_CBP*,i*_ to the additive genetic variance and having map length *R*_*i*_, we now define 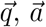, and 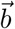 as having elements given by the equations

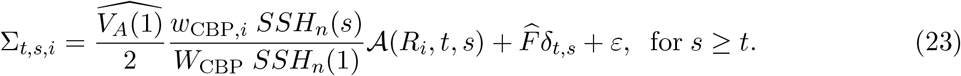

Again, the parameters of this model, 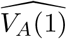 and 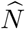, can be estimated with least squares. When analyzing genome-wide data, these various models could potentially be compared to an out-of-sample procedure, using inferred parameters to estimate the mean-squared predictive error between the two models for the remaining windows (Elyashiv et al. 2016). The confidence intervals for our method-of-moments estimates could be obtained through bootstrapping genomic windows since the errors are not identically and independently distributed.

### 3.1 Estimating the proportion of allele frequency change due to linked selection

We can also estimate what fraction of allele frequency change over *t* generations (Var(*p*_*t*_−*p*_0_)/*t*) is due to linked selection acting to perturb the frequency trajectories of neutral alleles. We have developed two approaches: first, a more conservative approach that considers only the contribution of selection to the temporal autocovariance, and second, a more exact approach that uses the estimated effective population size to include the contribution of selection to both variances and covariances of allele frequency change.

First, a simple estimate of the total fraction of the variance in allele frequency change (*G*) caused by linked selection is

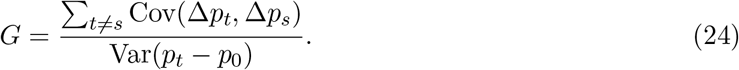

However, this estimator is conservative because it ignores the contribution that linked selection has on the variance in allele frequency change across a single generation (the Var(Δ_*H*_*p*_*t*_) term in Equation 2). If we include these variance terms, we have a less-conservative estimator we call *G*’,

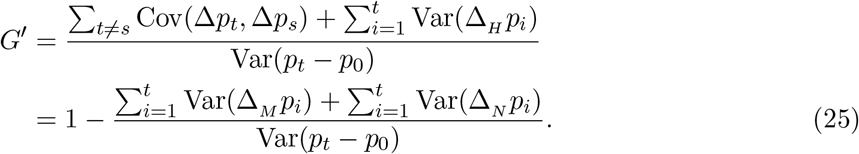

We can think of the numerator of the second term in Equation (25) as the variance in allele frequency change in a Wright–Fisher population without selection. Recall that under a Wright– Fisher model, the standardized variance across t generations is approximately (1 − exp(−*t*/2*N*)) ≈ *p*_0_(1 − *p*_0_) × *t*/2*N*, where this second approximation works for short time spans (*t*/2*N* ≪ 1). This suggests that we can use our method-of-moments estimate of the effective population size without the effects of selection, 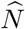, and compare the fraction of standardized variance we expect under this rate of drift to the empirical standardized variance,

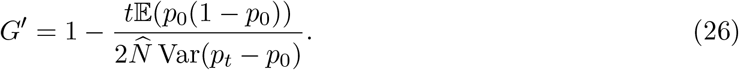

Figure 6 shows the estimated *G*’ from the method-of-moment 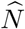 estimates using 20 replicates of simulated data. We learn three important points about our *G*’ estimator. First, for low *V*_*A*_, or *V*_*A*_ = 0, the estimator is quite noisy. Second, although the signal can be noisy for low *V*_*A*_, the relationship between *G*’ and level of recombination is consistent with selection affecting the total variance in allele frequency changes across the genome. Finally, this suggests that a negative relationship between *G*’ and recombination rate, calculated in windows across the genome, is a robust signal of linked selection impacting the total variance in allele frequency change. Furthermore, we would expect a positive relationship between the number of coding basepairs per window (when such information is available) and *G*’, which could serve as another robust signal of linked selection impacting the total variance in allele frequency change.

**Figure 6:**
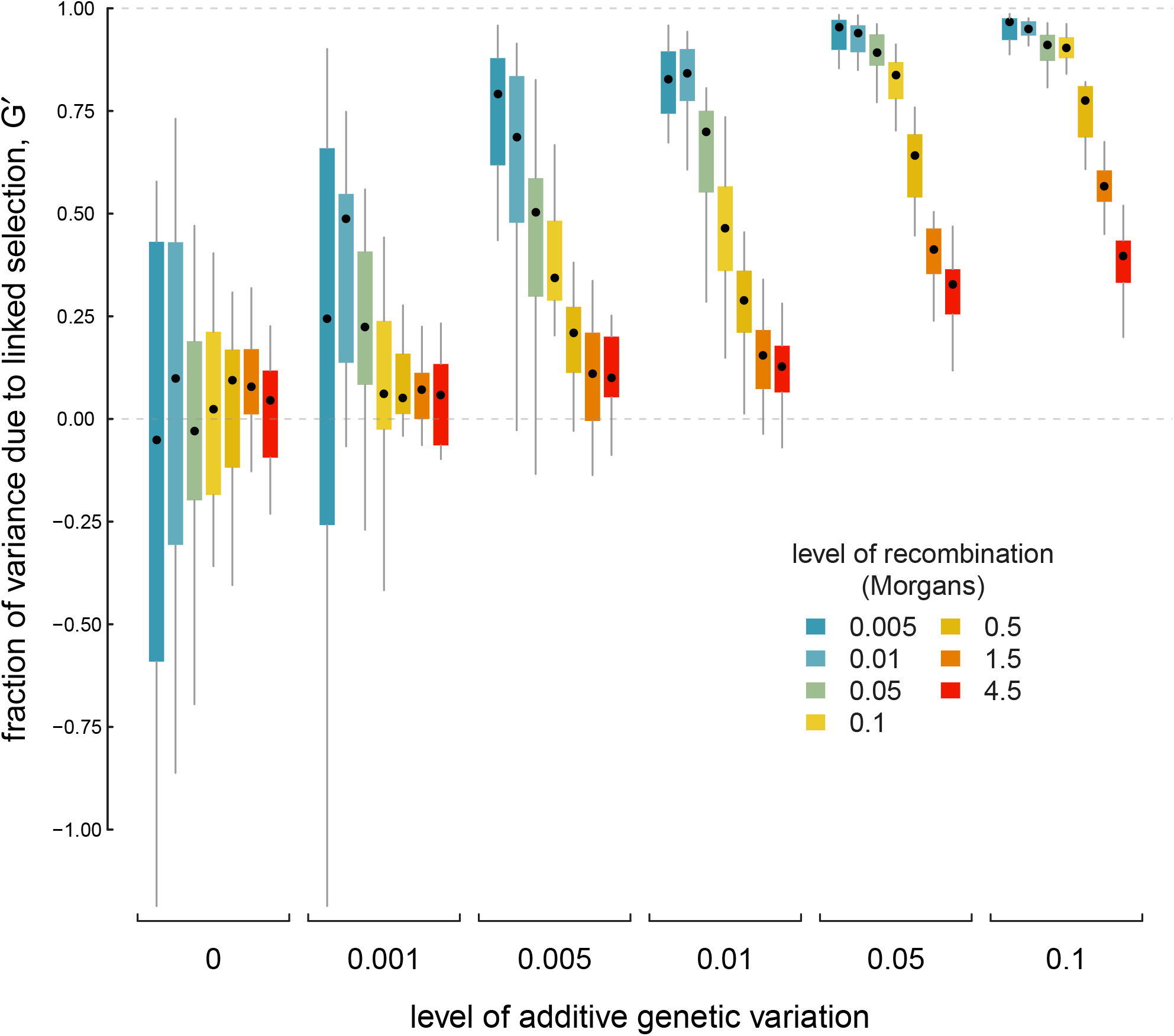
The proportion of total variance in allele frequency changes caused by linked selection, *G*’, across a variety of different levels of additive genetic variance (each group of boxplots), and different levels of recombination (each colored boxplot within a group). Each boxplot shows the spread of values across 20 replicates, with 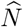 being calculated across each replicate.

### 3.2 Fluctuating Selection

Thus far we have assumed that fitness effect sizes are constant through time, that is *α*_*t,l*_ = *α*_*s,l*_, for all *t*, *s*. In natural populations, changes in the environment or composition of the population may cause these effect sizes to change through time, due to changing selection pressures and changes in the epistatic environment experienced by alleles. If these changes occur within the timeframe of recorded allele frequency changes, the levels of temporal autocovariance will differ from the levels predicted from our directional selection theory. However, from Equation (6) we can see that the magnitude of temporal autocovariance is determined in large part by 𝔼(*α*_*t*_*α*_*s*_).

Here, we discuss how temporal autocovariance behaves under an example of strong fluctuating selection: when selection on a trait changes direction at some point. Specifically, we change the fitness function 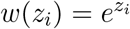 to 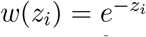 after some timepoint *t*^∗^; this is equivalent to changing *α*_*s,l*_ = − *α*_*t,l*_ iff *s* ≥ *t*^∗^, and *α*_*s,l*_ = *α*_*t,l*_ otherwise for all other *s* < *t*^∗^.

When such a strong change in the direction of selection occurs, the temporal autocovariance between timepoints before and after the change becomes negative, since temporal autocovariance is determined by the product *α*_*t*_*α*_*s*_ for *t* ≠ *s* (here we are holding effects constant across loci). We have validated this using the same simulation procedure as described in Section 2.4, except on generation 15 we reverse the direction of selection on the trait by changing the fitness function *w*(*z*) = *e*^*z*^ to *w*(*z*) = *e*^−*z*^. In Figure 7A, we show the temporal autocovariance Cov(Δ*p*_5_, Δ*p*_*s*_) for varying *s* along the x-axis (in this case, *V*_*a*_ = 0.05, *R* = 0.1, and *L* = 500). During the first five generations, the temporal autocovariance behaves as it does under directional selection, decaying due to the decrease in additive genetic variance and the breakdown of linkage disequilibrium. Then, on the 15^th^ generation, the direction of selection on the trait with breeding value *z*_*i*_ reverses and temporal autocovariance becomes negative since for all *α*_*s,l*_*α*_*t,l*_ < 0 for all *s* ≥ *t*^∗^ and *t* < *t*^∗^. Under this simple flip in the direction of selection pressure, the genic variance *V*_*a*_(*s*) = 2*α*^2^∑_*l*_*p*_*s,l*_(1 − *p*_*s,l*_), the expressions for the temporal autocovariance can be replaced with *V*_*a*_(*s*) = 2*α*_*s*_*α*_*t*_∑_*l*_*p*_*s,l*_(1 − *p*_*s,l*_, akin to a genetic/genic covariance. In Figure 7A, the gray line is our predicted level of temporal autocovariance proportional to 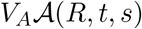 given by Equation (8) before generation 15, and after generation it is proportional to 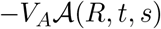 (using the empirical additive genetic variance). Note, however, that the dynamics of additive genetic variance under fluctuating selection are more complex than under directional selection. Whereas under directional selection the genetic variance decays as selection proceeds, under fluctuating selection there can be a transient increase in the additive genetic variance (seen in Figure 7A between generations 15-24). This transient inflation of the additive genetic variance is caused by the increase in the heterozygosity of haplotypes that had experienced reduced heterozygosity due to directional selection. With the direction of selection reversed, previously selected haplotypes move to more intermediate frequencies, which increases the additive genetic variance until selection proceeds and this variance decays (generations 25 and onwards). Overall, the dynamics of additive genetic variance under fluctuating selection are more complicated than under directional selection which makes inference using our sum of site heterozygosity approximation infeasible.

**Figure 7:**
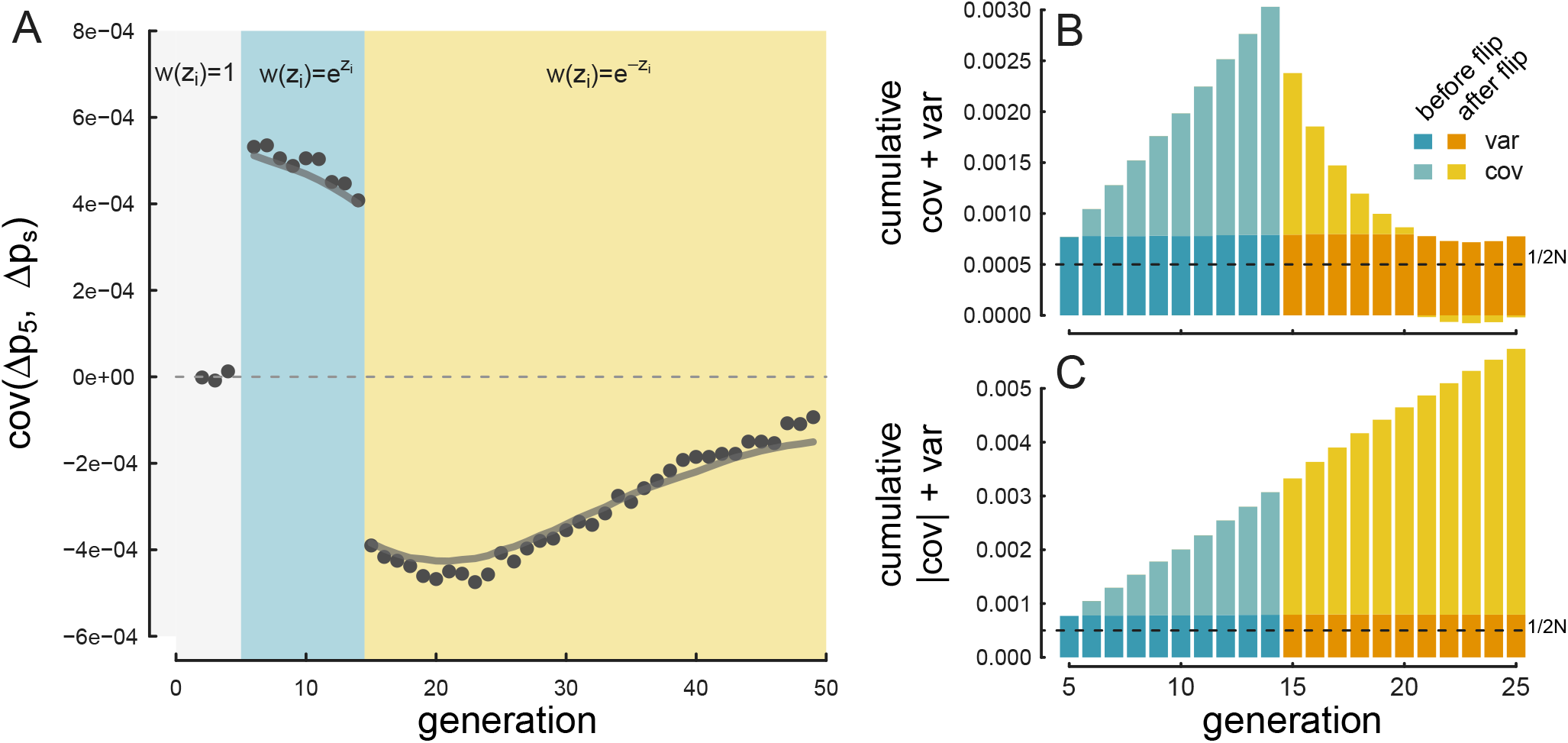
A: The covariance Cov(Δ*p*_5_, Δ*p*_*s*_), where *s* is varied on the x-axis, for *V*_*a*_ = 0.05, *R* = 0.1, and *L* = 500 averaged over 100 replicates. Selection begins in generation 5 (with the fitness function 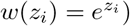 and on generation 15 the direction of selection flips (and the fitness function becomes 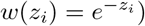. The gray line shows our directional selection temporal autocovariance prediction modified so that after the flip in the direction the trait is selected, we plot the *negative* theoretic level of temporal autocovariance. B: The average cumulative variance and covariances through the generations for the same simulation parameters, with the height of the bar representing the total cumulative variation Var(*p*_*t*_ − *p*_0_)/*t*. Since the direction of selection flips, the covariances terms after generation 15 become negative, leading the total variance to decrease (the negative covariances are plotted below the x-axis line). After generation 21, the total covariance is negative, leading the total variance to dip below the level of variance alone (determined by drift and heritable fitness variation). The dark gray dashed line shows the level of variance expected by drift alone (Var(*p*_*t*_−*p*_0_)/*t* = 1/2*N*). C: The effect of using the absolute value of covariance, which prevents the negative autocovariances from canceling out the effects of other covariances before the direction of selection changed.

**Figure 8:**
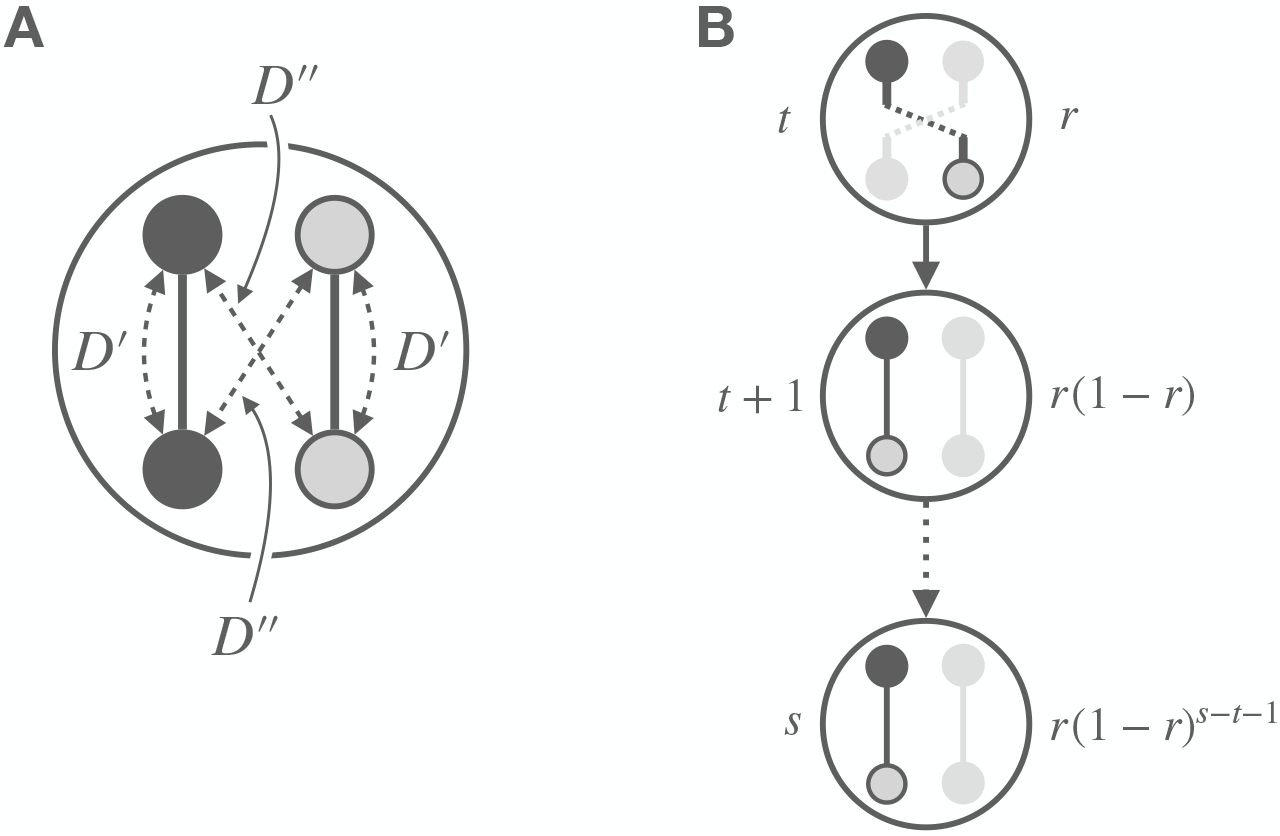

A: An illustration of gametic (*D*^*’*^) and non-gametic (*D*^*’’*^) linkage disequilibria between two loci in a diploid. B: An illustration of how non-gametic linkage disequilibria in generation t is converted to gametic linkage disequilibria through recombination (which happens with probability r, the recombination fraction between the two loci), and is then maintained until generation s with probability (1 r)^*s−t−*1^. The gray loci on the gray gamete indicate the homologous, but not tracked focal association. Overall, the covariance created by the conversion of non-gametic LD to gametic LD is E(*Ds’ Dt’’*) = r(1 − r)^*s−t−*1^E ((*Dt*’’)^2^).

Since fluctuating selection can create *negative* temporal autocovariance, the total amount of autocovariance over time (e.g. ∑_*t≠s*_ Cov(Δ*p*_*t*_, Δ*p*_*s*_)) can misrepresent the actual amount that linked selection is affecting allele frequencies on shorter timescales. This, in turn, leads our estimator *G*, for the fraction of variance in allele frequency due to linked selection, to underestimate the total contribution of linked selection to variation in allele frequencies over the time period. We show an example of this in Figure 7B, which depicts the total variance in allele frequency change Var(*p*_*t*_−*p*_0_)/*t* through time, partitioned into variance and covariance components and colored according to whether the generation was before and after the reverse in directional selection. The covariances ∑_*t≠s*_ Cov(Δ*p*_*t*_, Δ*p*_*s*_) increase as they would under directional selection, but after generation 15, the contribution of covariance begins to decrease as negative autocovariances accumulate. By generation 20, the total covariance terms have a net negative effect, and actually act to decrease the total variance Var(*p*_*t*_−*p*_0_)/*t* for a few generations below the constant level expected under drift and heritable variation.

To more fully capture the contribution of selection allele frequency change, we modify G, using the absolute value of the covariances,

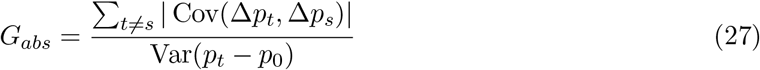

which prevents negative temporal autocovariances from canceling out the effects of positive temporal autocovariances, since both are a reflection of linked selection acting on neutral allele frequency changes. We show *G*_*abs*_ in Figure 7C, where even after the change in the directional selection we see a steady accumulation of covariance contributing to total variance Var(*p*_*t*_−*p*_0_)/*t*. This also suggests one plausible way to check for genomically wide-spread fluctuating selection would be to test if *G*_*abs*_ > *G*. However, we note that in contrast to our simulations, it is likely that in natural populations only a subset of alleles change their relationship to fitness. This may act to dampen the magnitude of, but not completely reverse the direction of genome-wide temporal autocovariance, and so different approaches may be needed to identify fluctuations.

## 4 Discussion

Currently, the prevailing empirical approach to studying linked selection relies on using samples from a single timepoint and modeling the patterns of diversity subject to different functional constraints and in different recombination environments. The early theoretic work underpinning empirical analyses of linked selection’s effects on diversity were primarily full sweep, recurrent hitchhiking models, where beneficial mutations arise in the population and then sweep to fixation (Kaplan et al. 1989; Maynard Smith and Haigh 1974; Stephan et al. 1992). Furthermore, by looking at the patterns of diversity around amino acid substitutions or the site frequency spectrum in low recombination regions, researchers have teased apart the effects of background selection and hitchhiking in *Drosophila* (Begun et al. 2007; Elyashiv et al. 2016), humans (Hernandez et al. 2011; McVicker et al. 2009), and some plants species (Beissinger et al. 2016; Nordborg et al. 2005; Schmid et al. 2005; Williamson et al. 2014). Yet, as other theoretic models of hitchhiking incorporating changes in the environment (Kopp and Hermisson 2007, 2009a,b), sweeps originating from standing variation (Hermisson and Pennings 2005), and multiple competing beneficial haplotypes (Pennings and Hermisson 2006) have been developed, it has become rather more difficult to detect the signal of these hitchhiking phenomenon in empirical data. We’ve proposed here that temporal autocovariance offers a unique and measurable signal of linked selection over shorter timescales that provides a fuller picture of the ways in which genome-wide diversity has affected by these other hitchhiking phenomena.

### 4.1 Empirical Applications and Future Directions

Here, we’ve developed expressions for temporal variances and autocovariances, and applied these to model temporal variances and covariances during directional selection on a trait. We’ve demonstrated how one can (1) estimate the additive genetic variance for fitness and the drift-effective population size, (2) estimate the fraction of variance in allele frequency change due to linked selection, and (3) evaluate whether fluctuating selection is operating from these temporal variances and autocovariances. However we recognize a series of limitations and difficulties when applying these methods to empirical data and natural populations.

First, one difficulty with temporal sampling of natural populations is the risk that the genetic composition may change drastically due to migration or biased sampling across timepoints. Since our theory assumes a constant-sized and isolated population, migration into the sampled population presents a serious potential confounder. For example, seasonal migration could create an influx of new alleles that could at best dampen signals of directional selection across seasons, or at worst create a signal of artificial covariance between like seasons. Similar biases could occur if a sampling method incidentally preferred certain subgroups in a stratified population. This could occur, for example, if individuals differ in some behavior affecting their likelihood of being sampled across different temporal environments, which could cause spurious covariances. While such sampling issues and migration might be able to be detected *post hoc* from exploratory data analysis approaches like PCA, studying isolated natural populations and carefully designing sampling schemes would lead to the best inference. Importantly, the effects of gene flow and other temporal inhomogeneities could differ across recombination environments, as among population differentiation will be more pronounced in regions of low recombination and high functional density (Burri 2017; Keinan and Reich 2010; Nachman and Payseur 2012). In situations where migration is a factor, one way forward might be to study the contribution of linked selection after partitioning out the effects of gene flow across recombinational and functional environments by extending admixture inference approaches that estimate the genome-wide effect of drift and admixture.

Second, our method-of-moments approach relies on assuming we can approximate the dynamics of additive genetic variance through time by using the empirically observed changes in sum of site heterozygosity in a region as a proxy for its decay. While we have shown this model works under directional selection in a relatively idealized setting, inference in natural populations may be complicated by changes in the environment that induce the effect sizes across loci to vary across timepoints. While our fluctuating selection results show that our directional selection theory extends to changes in the direction of selection with minor adjustments (e.g. Figure 7), having the effect sizes vary across each generation would complicate the dynamics of additive genetic variance through time and make inference of *V*_*A*_ difficult. Similarly, we assume that effect sizes are constant across sites. Variance in effect sizes across a region will not bias our results unless there is covariance between effect size and local recombination rate. Further work is required to develop statistical methods to test for violations of our assumptions about effect sizes remaining constant across time, and to potentially incorporate these complications into inference.

Along similar lines, we have focused on directional selection under a multiplicative model. However, selection experiments, a natural place to apply our method, often use truncation selection, which generates systematic epistasis for fitness and thus linkage disequilibria among loci (Bürger 2000; Walsh and Lynch 2018). Similarly, in natural populations stabilizing selection will act on many traits, which can also generate LD among loci. These selective processes will act to rapidly reduce the additive genetic variance for fitness across time, especially in low recombination regions, and to reduce the initial additive genetic variance in low recombination regions. Simulating the effect of truncation selection and stabilizing selection on temporal covariances in allele frequencies seems a useful direction. Our model may prove to be a useful null model of selection that the complications of epistasis could be tested against.

Finally, while we demonstrate our method-of-moments estimation approach is simple and leads to unbiased estimators, we see there are opportunities for simple extensions and other inference procedures. Differences in recombination rates and coding density are relatively easily accommodated (Equation 23). In fitting our model we assumed a parametric form to the initial LD between neutral and selected loci; however, in practice the initial linkage disequilibria between neutral polymorphisms and putatively functional sites could be estimated empirically. Using the empirical LDs in Equation (23) could make the inference somewhat analogous to LD Score regression (Bulik-Sullivan et al. 2015). The LD Score of a SNP is simply the sum of the *R*^2^’s of the focal SNP to all SNPs within some large physical genomic window, which could be used in place of the integral in Equation (10). Using this equation, *V*_*A*_ could be estimated by regressing the temporal covariance of a SNP on its LD Score. An alternate approach would be to use likelihood methods to model each neutral site’s frequency changes using the set of pairwise linkage disequilibria and recombination distances between the neutral site and all neighboring polymorphic sites. Then, genome-wide or region-wide estimates could be found via composite likelihood methods, in a similar manner to McVicker et al. (2009) and Elyashiv et al. (2016). Furthermore, one could include different *V*_*A*_ parameters for neighboring polymorphic sites with specific functional annotations, such as those in genic regions, introns, exons, etc. to see how different classes of sites contribute to the additive genetic variance for fitness. Our hope is that statistical methods to quantify the effects of linked selection over short timescales will improve and be combined with measures of phenotypic change, leading to a more synthetic view of how selection on ecological timescales occurs at the genetic and phenotypic levels.

As the number of empirical temporal genomic studies continue to increase, it is worth mentioning how our study of temporal autocovariance suggests a few ways to optimize experimental design to increase the power to differentiate the effects of selection from drift. First, one should ideally sample frequencies from consecutive generations for at least some timepoints during the duration of the experiment. This is because the variance in allele frequency change Var(Δ*p*_*t*_) between adjacent generations is only impacted by the heritable variance in offspring number, Var(Δ_*H*_p_*t*_) (see Equation 2), and not by the accumulation of temporal autocovariance terms (e.g. Equation 3); this allows for more accurate estimates of *G*. In cases where a long study duration is needed but sequencing is limited to only a subset of generations, a mixed duration sampling design, such as sampling generations 1 through 4, and then 10 through 14, and so forth could serve as a compromise. Second, as described in Appendix Section A.8, the shared sampling noise between adjacent timepoints creates a negative bias in autocovariance that must be corrected for. As described in Equations (96) and (97), we can estimate this bias from data, but this introduces additional uncertainty into our parameter estimates. In cases where the experimenter suspects *a priori* that fluctuating selection is occurring, e.g. between two seasons, we recommend at least two temporal samples per season. This allows one to differentiate negative covariance occurring from the bias correction procedure underestimating bias from negative covariance caused by fluctuating selection through comparing non-adjacent timepoints that differ in season. Finally, once can directly remove the effects of the technical sampling noise created by variation in sequencing by dividing up temporal samples and barcoding them into two groups (e.g. A and B). Then, the sample covariance estimate Cov(Δ*p*_*t,A*_, *p*_*s,B*_) does not share the technical sampling noise, reducing the bias (but note some bias remains due to the sampling process where individuals are sampled from the population.)

### 4.2 Connecting Temporal Linked Selection with Single Timepoint Studies

Our goal in this paper is to suggest that quantifying variance and autocovariance using temporal data sets can help us understand the impact linked selection has across the genome on short timescales, which supplements our current view informed mainly by single timepoint studies. A range of approaches to estimate the parameters and impact of models of linked selection from a single contemporary timepoint have been developed (Begun and Aquadro 1992; Elyashiv et al. 2016; Hudson 1994; McVicker et al. 2009; Sella et al. 2009; Wiehe and Stephan 1993). These estimates necessarily reflect linked selection over tens to hundreds of thousands of generations. One question is whether these estimates of the proportion of allele frequency change due to linked selection should line up with those over shorter time periods? Some forms of linked selection may be fairly uniform over time, whereas rare, strong sweeps will have a huge impact on long-term patterns of variation but may be hard to catch in temporal data. Conversely, as we discuss below, fluctuating selection may lead to stronger signals of linked selection on short timescales than seen in long-term snapshots.

Studies of contemporary data have revealed multiple lines of evidence for the effect of linked selection in a variety of taxa. If linked selection is pervasive across the genome, diversity could be severely dampened as most sites would be in the vicinity of selected sites, thus reducing the genomewide level of diversity without leaving strong local signals differentiated from the background. This is one proposed resolution of Lewontin’s paradox, the observation that diversity levels occupy a narrow range across taxa with population sizes that vary by orders of magnitude (Gillespie 2001; Leffler et al. 2012; Lewontin 1974; Maynard Smith and Haigh 1974). Elyashiv et al. (2016) estimated a 77 - 89% reduction in neutral diversity due to selection on linked sites in *Drosophila melanogaster*, and concluded that no genomic window was entirely free of the effect of selection. Similarly, Corbett-Detig et al. (2015) has found evidence of a stronger relative reduction in polymorphism due to linked selection in taxa with larger population sizes. However, these reductions fall short of the many orders of magnitude required for linked selection to explain Lewontin’s paradox (Coop 2016).

One limitation of these approaches is that they require estimating *π*_0_, the level of diversity in the absence of linked selection, usually from the diversity in high-recombination regions with low gene content. The average genome-wide reduction of diversity can then be judged relative to *π*_0_. Ideally, *π*_0_ would be a measure of the average diversity due entirely to drift and demographic history, i.e. unaffected by heritable fitness variation. However, there are two complications with this. First, as Robertson (1961) first showed, even a site completely unlinked from sites creating heritable fitness variation experiences a reduced effective population size due to the total additive genetic variance for fitness at these unlinked sites, and thus lower diversity (see also Santiago and Caballero 1995). The second complication is that if linked selection is sufficiently strong, the bases used to measure π_0_ may not be sufficiently unlinked from fitness-determining sites to plateau to the Robertson (1961) level of diversity, a known potential limitation (Coop 2016; Elyashiv et al. 2016). Overall, the empirical studies relying on present-day samples from a single timepoint could be underestimating the effects pervasive linked selection has on diversity. If linked selection can be observed over suitable timescales in temporal data, we might be able to disentangle some of these effects. For example, if high recombination regions still show temporal autocovariance in allele frequency change, we would have evidence that even these regions are not free of the effect of linked selection and we might be able to estimate its long term impact on levels of diversity.

Temporally or spatially fluctuating selection has long been discussed as a explanation for abundant, rapid phenotypic adaptation over short timescales, yet over longer timescales both phenotypic changes and molecular evolution between taxa are slow (Gingerich 1983; Hendry and Kinnison 1999; Messer et al. 2016). However, most of our approaches to population genomic data are built on simple models with constant selection pressures, as typically we have not had the data to move beyond these models (Messer et al. 2016). Currently, many approaches to quantify the impact of linked selection due to hitchhiking assume classic sweeps, where a consistent selection pressure ends in the fixation of a beneficial allele (Hernandez et al. 2011; Sella et al. 2009; Wiehe and Stephan 1993). However, fluctuating selection can have a larger effect on reducing diversity than classic sweeps (Barton 2000) depending on the timescales over which such fluctuations occur. In fact as Barton (2000) points out, the total effect of classic Maynard-Smith and Haigh-type sweeps on diversity is limited by the relatively slow rate of substitutions. We show that when the direction of selection on a trait abruptly reverses, this creates negative autocovariance between the allele frequency changes before and after the reverse in direction. We can observe the shift by plotting autocovariances over time and noting when they become negative, indicating a negative additive genetic covariance between fitness at two timepoints. Here we assume a simple form of fluctuating selection: where selection pressures on all of our sites flip at some timepoint. In reality, selection pressures will change on only some traits, and some of the genetic response will be constrained by pleiotropy, thus only some proportion of the additive genetic variance will change. Still, we expect some level of negative covariance after a reversal in the direction of selection, and there is additional signal of fluctuating selection by comparing how the strength of temporal autocovariance varies with recombination and the initial level of linkage disequilibrium in the genome.

### 4.3 Connecting Estimates of *V*_*A*_ from Temporal Genomic Data and Quantitative Genetic Studies

The temporal covariance of allele frequencies potentially offers a way to estimate the additive genetic variance for fitness, as illustrated by our method-of-moments approach across genomic windows. The additive genetic variance for fitness can, like any other trait, be estimated through quantitative genetics methods, which exploit the phenotypic resemblance between relatives and their known kinship coefficients (Kruuk 2004; Shaw and Shaw 2013; see Hendry et al. 2018 for a review), and these methods have been applied to estimate the additive genetic variance for fitness from natural populations (Burt 1995; Mousseau and Roff 1987). Ideally, one could reconcile quantitative genetic measures of fitness variance with estimates from allele frequency covariance. For example, Charlesworth (2015) undertook a similar analysis in *Drosophila melanogaster*, comparing population genetic estimates of fitness variance to quantitative genetics estimates, highlighting a discordance potentially consistent with undetected large-effect alleles that are likely maintained by some form of balancing selection. By allowing us to directly measure fitness variation from population genetic data over very short timescales, temporal data could help untangle the causes of this discordance. A natural extension of this would be to see which regions contain the greatest inferred levels of additive genetic variance for fitness, and test for functional covariates such as the number of coding bases, etc. Whereas previous temporal studies have focused on finding loci under selection, inferring the level of additive genetic variance could provide a more complete view of how much selection operates over short timescales.

### 4.4 Concluding Thoughts

With temporal data we can directly partition the total variance in allele frequency change across generations, Var(*p*_*t*_ − *p*_0_) into components according to the underlying process governing their dynamics: drift and linked selection. Since the trajectory of a neutrally drifting polymorphism does not autocovary, evidence of temporal autocovariance across neutral sites in a closed population is consistent with linked selection perturbing these sites’ trajectories. If we consider drift to be the process by which non-heritable variation in reproductive success and Mendelian segregation cause allele frequencies to change, then this is estimable from and separable from the effects of linked selection using temporal data. This helps frame the long-running debate about the roles neutral drift and linked selection have in allele frequency dynamics into a problem that can potentially be directly quantified by the contribution of each distinct process with temporal data.

## Supporting information

highlighted revisions from last bioRxiv version

## 5 Code and Data Availability

All code and simulation data to reproduce these results are available on GitHub at https://github.com/vsbuffalo/tempautocov.

## 6 Acknowledgments

We thank Nick Barton, Doc Edge, Matt Osmond, Enrique Santiago, Michael Turelli, and two anonymous reviewers for feedback on previous versions of the manuscript, and Aneil Agrawal, Dave Begun, Sarah Friedman, Bill Hill, John Kelly, Tyler Kent, Chuck Langley, Sally Otto, Jonathan Pritchard, Kevin Thornton, Anita To, and members of the Coop lab for helpful conversations. This research was supported by an NSF Graduate Research Fellowship grant awarded to VB (1650042), and NIH (R01-GM108779) and NSF (1353380) awarded to GC.

## A Appendix

**Table 1:**
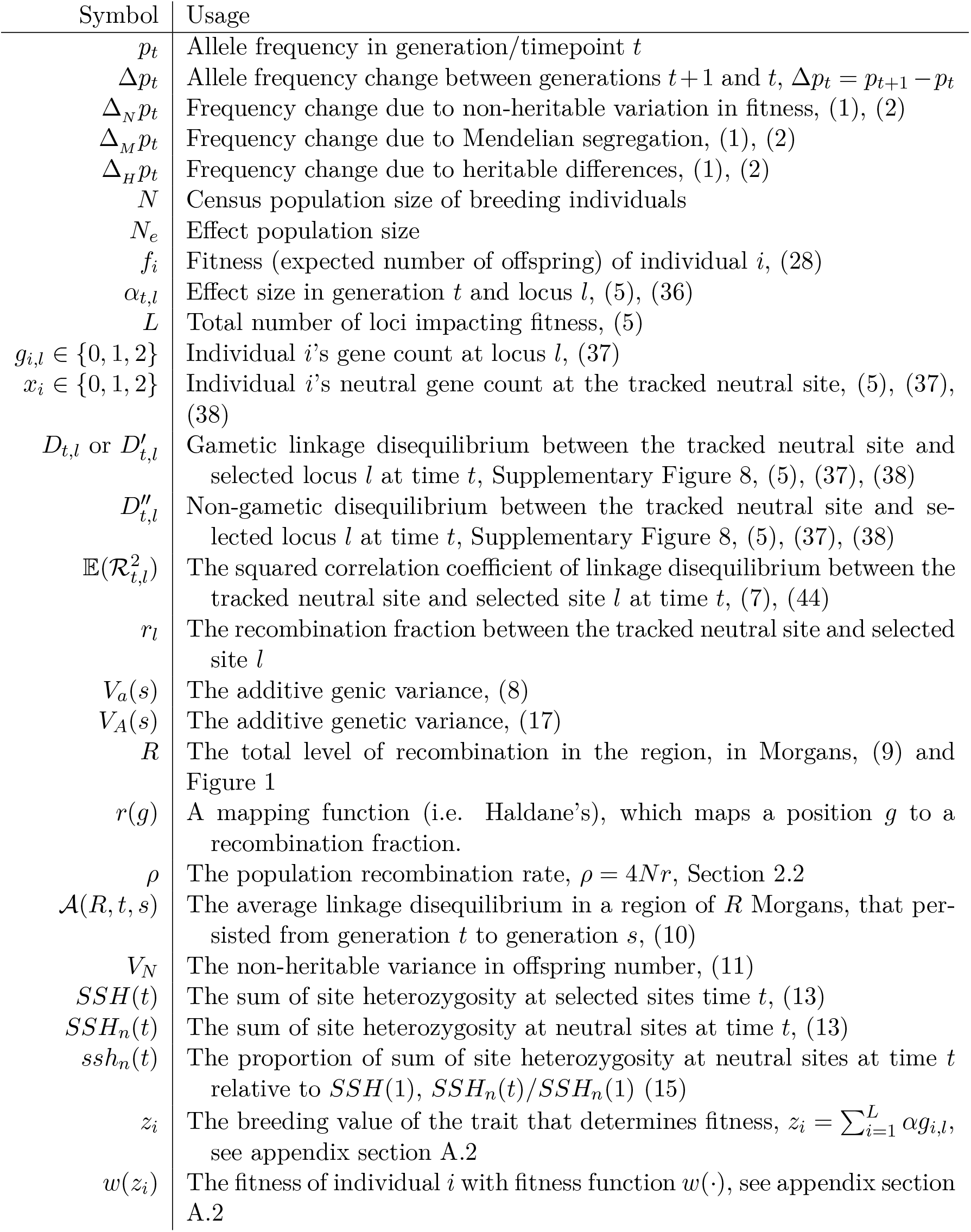
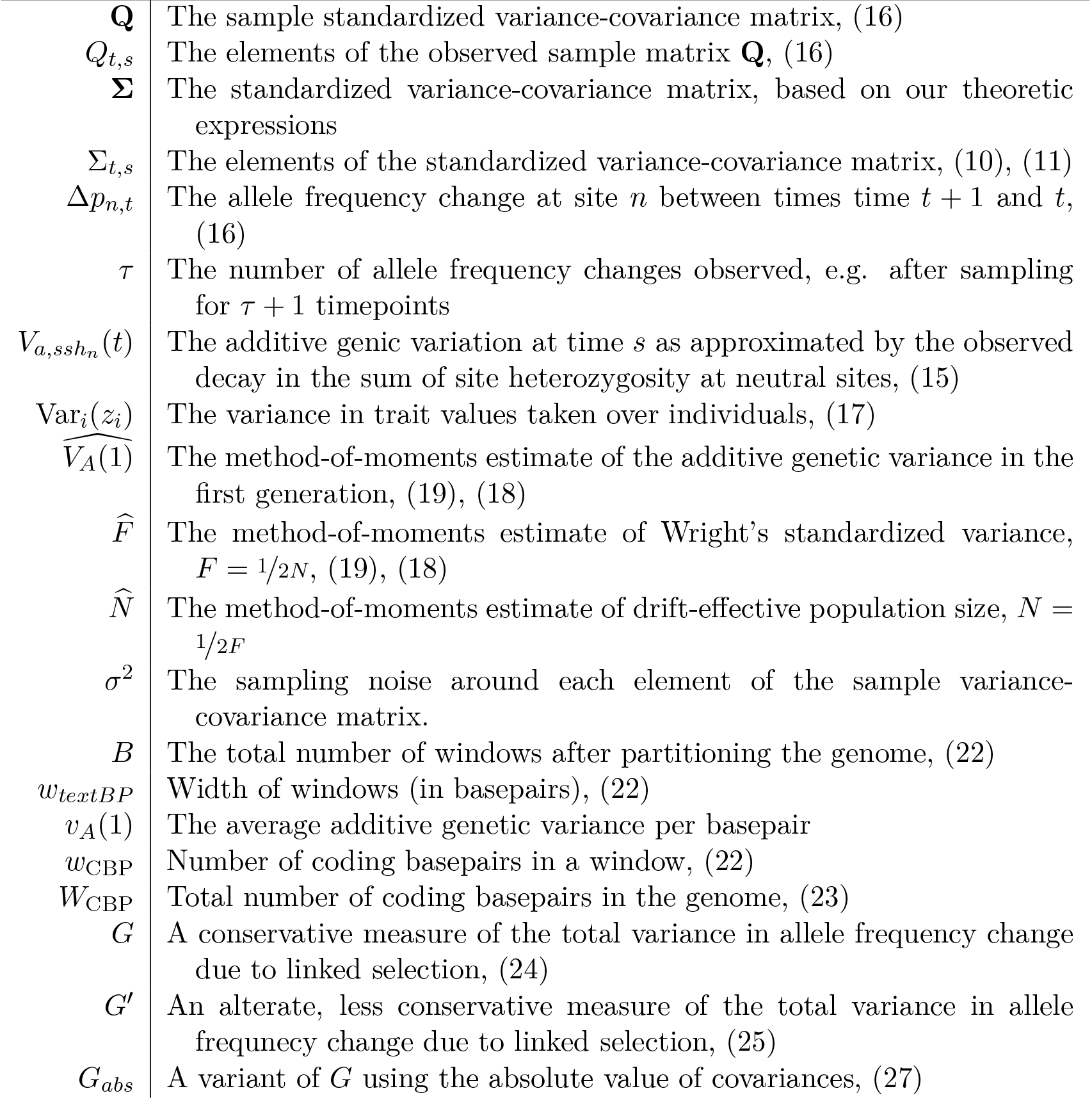
Notation

### A.1 Decomposition of Allele Frequency Change

This decomposition of *neutral* allele frequency change between two consecutive generations is based on that of Santiago and Caballero (1995). We imagine a closed Wright–Fisher population of *N* diploids, where each diploid *i* contributes *k*_*i*_ ~ Multinom(*f*_*i*_/*N*, 2*N*) gametes to the next generation. We assume the population size is constant, such that one diploid begets one diploid and thus 𝔼_*i*_(*f*_*i*_) = 1. The neutral allele frequency in the next generation can be thought of as each of the *N* parents passing their average genotype *x*_*i*_/2 (where *x*_*i*_ ∈ {0, 1, 2} is the number of tracked neutral alleles individual *i* carries) to their *k*_*i*_ gametes, plus a random Mendelian deviation *b*_*ij*_ ∈ {0, −1/2, 1/2} to each offspring j. Then, the frequency in the next generation can be written as

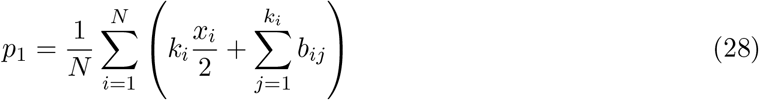

where 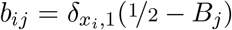, *B*_*j*_ ~ Bernoulli(1/2) and 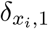 is an indicator function that is one when the individual *i* is a heterozygote (i.e. *x*_*i*_ = 1), and zero otherwise.

If we further decompose the number of offspring of individual *i* into the genetic and non-genetic contributions, *k*_*i*_ = *f*_*i*_ + *d*_*i*_, then

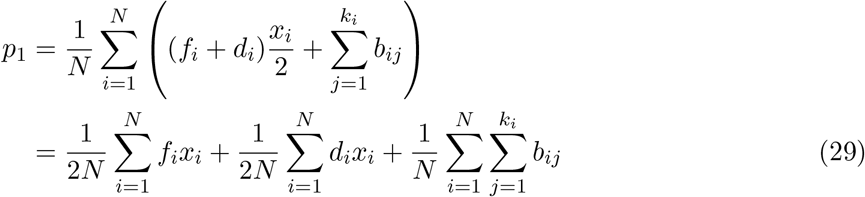

and the change of the neutral allele’s frequency is the difference Δ*p* = *p*_1_ − *p*_0_ where 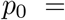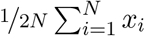. Then,

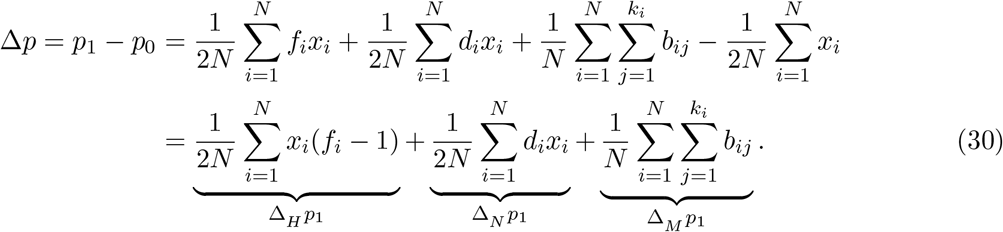

These *d*’s broadly capture non-heritable variation in an individual’s offspring number, with 𝔼[*d*_*i*_] = 0. In a quantitative genetics framework Var_*i*_(*d*_*i*_) can include non-genetic variation in the lifetime reproductive success of individuals (*V*_*E*_), in population genetic models Var_*i*_(*d*_*i*_) can accommodate the sampling of parents to form the next generation, e.g. multinomial sampling of individuals from fitnesses (Santiago and Caballero 1995). From now forwards, we assume that variation in these d’s is non-heritable. We note that in non-panmictic populations, chance covariances could be created between a neutral polymorphism and environmental component of their phenotype, especially as a population expands its range into new environments that affect the phenotype, and variants “surf” to higher frequencies (Edmonds et al. 2004; Excoffier and Ray 2008; Hallatschek and Nelson 2008).

Note that by construction, the allele frequency change components Δ_*N*_ *p*_*t*_ Δ_*M*_ *p*_*t*_, and Δ_*H*_ *p*_*t*_ are orthogonal within each individual, given the neutral allele frequency *x*. Partitioning the allele frequency change for an individual *i* into its components,

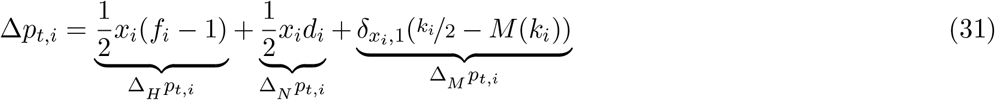

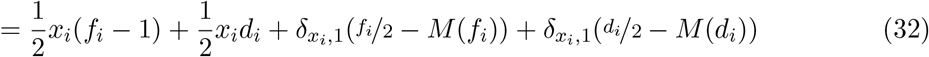

where 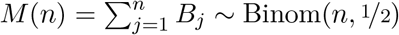.

Two random variables *X*, *Y* are uncorrelated if Cov(*X*, *Y*) = 𝔼(*XY*) − 𝔼(*X*)𝔼(*Y*) = 0, and are orthogonal if either has an expected value of zero such that 𝔼(*XY*) = 0. We show briefly that taking expectations *over conceptual evolutionary replicates*, the terms are orthogonal. First, the terms *x*(*f* − 1) and *xd* are orthogonal (dropping i subscripts),

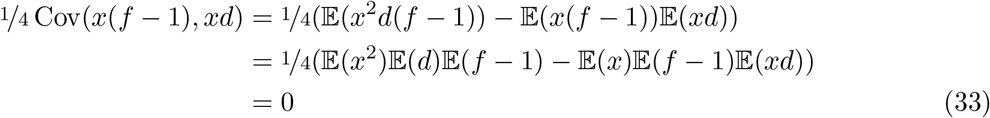

since 𝔼(*f*) = 1 across evolutionary replicates due to the assumption that population size is constant, and *x* ⊥ *f*, as across all evolutionary replicates, there is no dependence between a particular neutral allele an individual carries and their fitness (though in particular replicates, such associations occur). Similarly, for the case *x* = 1 (other cases are all zero, and can be ignored), it can be shown using the law of total expectation that *Cov*(*xd*, *d*/2 − *M*(*d*)) = 0 (and likewise with *x*(*f* − 1) and *f*/2 − *M*(*f*)). Note that across individuals within a population, there are weak covariances in their number of offspring as the total number of offspring must sum to *N*; under a Multinomial offspring distribution, these are of order 1/*N*.

### A.2 Temporal variance and autocovariance under multilocus selection

We assume the phenotype of an individual i has an additive polygenic basis, such that their breeding value is 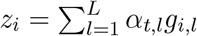 which deviates around a mean of zero, and *α*_*t,l*_ is the additive effect size at locus *l* in generation *t*, and *g*_*i,l*_ ε {0, 1, 2} is individual *i*’s allele count at this locus (note that effect of non-heritable environmental noise affecting the trait is accounted for in the *d*_*i*_ terms above). We impose directional selection on this trait using an exponential fitness function, such that individual *i*’s fitness is 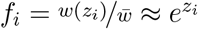 (assuming w̄ ≈ 1). If we assume individuals’ phenotypic values do not deviate too far from their mean value of zero, we can approximate *f*_*i*_ as: 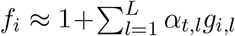. Then, we can write the change in neutral allele frequency due to only heritable variation in fitness (Δ_*H*_ *p*_*t*_) as a covariance between fitness and the neutral allele frequency across individuals in generation t,

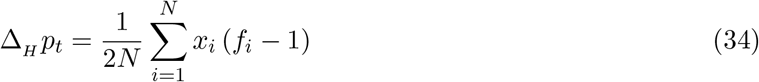

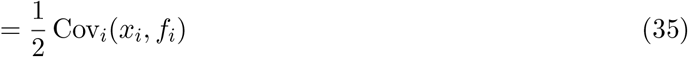

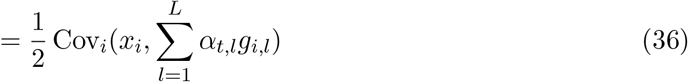

which is the is the Robertson-Price covariance (Lynch, Walsh, et al. 1998; Price 1970; Robertson 1966; Walsh and Lynch 2018).

Now, we break up the genotypic value *x*_*i*_ into the contributions of each of the two gametes that formed individual *i*, 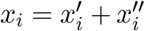, and likewise with the trait locus 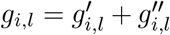, where 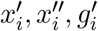, and 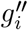 are all indicator variables. Expanding out the covariances, we have

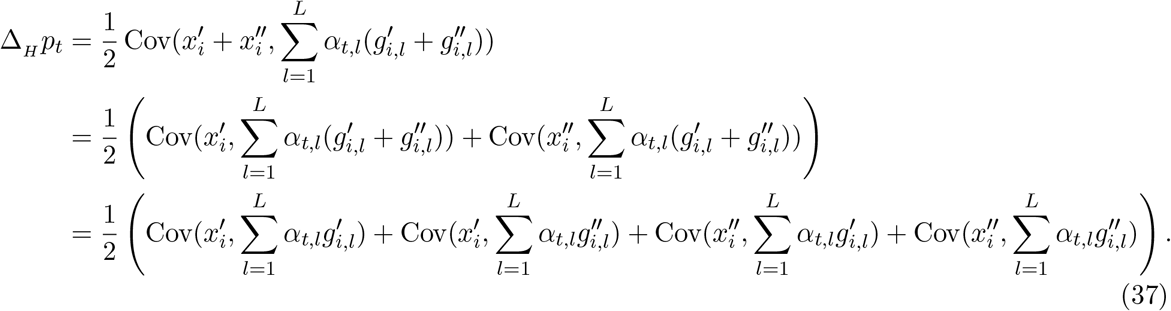

Each of these covariances is between the neutral allele and a selected allele, either on the same gamete (either maternal or paternal) or across gametes. These covariances can be written as linkage disequilibrium terms,

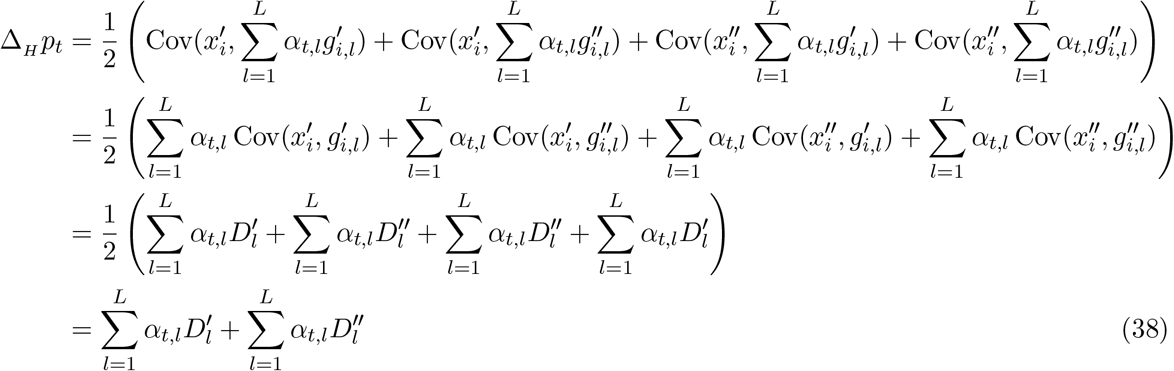

where D’_*L*_ is the linkage disequilibrium between alleles on the same gamete (the gametic LD), and the 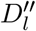 is the across-gamete LD (non-gametic LD); see Weir (1996), p. 121.

We ignore non-gametic linkage disequilibria 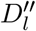 as these are weak under random mating, and write the multilocus temporal covariance between the allele frequency changes Δ*p*_*t*_ and Δ*p*_*s*_ as

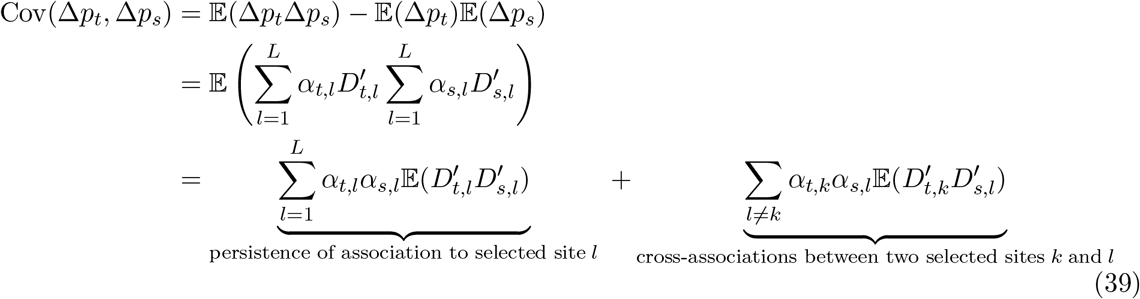

since 𝔼(Δ*p*_*t*_) = 0.

#### Modeling the dynamics of LD between selected and neutral sites

Here, we outline a model of the changes in linkage disequilibria between the focal neutral site and selected sites, which allows us to derive an expression for the first term in Equation (39). Typically, models of multilocus selection track the genetic changes in a population by transforming between a representation of haplotype frequencies to a representation of allele frequencies, linkage disequilibria, and higherorder linkage disequilibria (Barton and Turelli 1987, 1991; Turelli 1988; Turelli and Barton 1990). We for the moment avoid the difficulty of a full multilocus treatment by assuming that the linkage between selected sites is loose enough that one selected site’s frequency change is independent of the change at other selected sites, e.g. there is no selective interference; this was an assumption in past treatments (Santiago and Caballero 1995, 1998; see Barton 2000 for discussion of this). Specifically, we model the dynamics of the LD between the neutral site and each selected site as they would behave under a single sweep model.

We adapt Barton’s (2000) model for LD dynamics during a sweep. We imagine a polymorphic neutral locus has alleles *B*_1_ and *B*_2_ with frequencies *p* and 1 − *p*. We partition the allele frequency of *B*_1_ by conditioning on which allele at the selected site (either *A*_1_ or *A*_2_) is carried on the same background, e.g. *P*(*B*_1_|*A*_1_), and *P*(*B*_1_|*A*_2_), such that *P*(*B*_1_) = *P*(*B*_1_|*A*_1_)*P*(*A*_1_) + *P*(*B*_1_|*A*_2_)*P*(*A*_2_). Then, the linkage disequilibrium between neutral and selected sites can be expressed as *D* = *P*(*A*_1_)*P*(*A*_2_)(*P*(*B*_1_|*A*_1_) −*P*(*B*_1_|*A*_2_))). To simplify notation, we denote the frequency of the neutral allele on the different fitness backgrounds at time *t* by 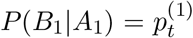 and 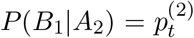 and the selected allele at locus *l* frequency as *p*_*t,l*_, we have the LD between the focal neutral site and selected site *l* at time *t* is

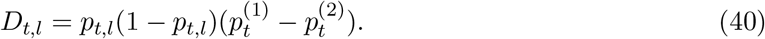

Selection changes the frequency *p*_*t,l*_ through time, and recombination acts to disassociate the neutral allele with its backgrounds. The expected difference 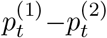 is maintained with probability (1 − *r*_*l*_) each generation (Barton 2000). If the initial generation is *t* and the future generation is *s* (*s* > *t*), and *r*_*l*_ is the recombination rate between the focal neutral locus and selected site l, this leads to

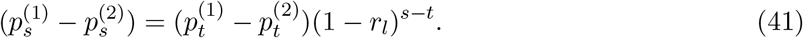

Then, we can use this to describe the dynamics of *D*_*s,l*_ to generation *t*,

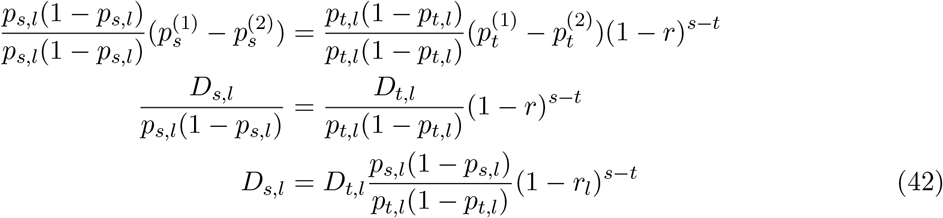

(c.f. Stephan et al. 2006, equations (30) and (31)). Now, we can find the expected product 𝔼(*D*_*t,l*_*D*_*s,l*_) by multiplying both sides by *D*_*t,l*_ and taking expectations. We treat the allele frequency trajectory as deterministic, giving us,

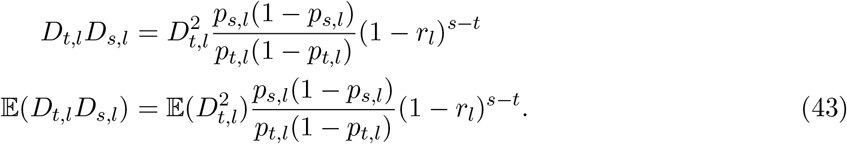

Then, we simplify this by replacing 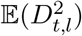 with 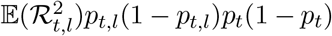 (where 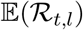 is the square of the correlation between the neutral site and selected site *l* at time *t*; Hill and Robertson 1968),

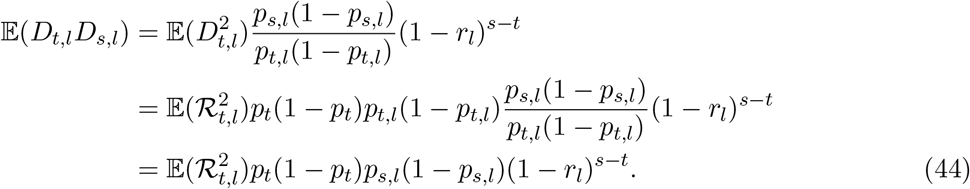

Returning to Equation (39) and replacing the 𝔼(*D*_*t,l*_*D*_*s,l*_) terms with our expression above,

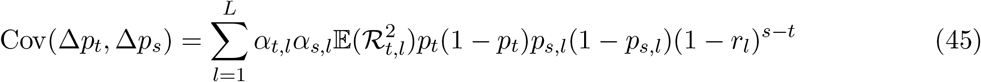

again ignoring the cross-associations between selected sites. Dividing our temporal covariance by the neutral site’s *p*_*t*_(1 − *p*_*t*_), we can write the multilocus temporal covariance in a standardized form (analogous to Wright’s F),

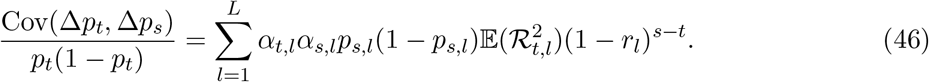

#### Using average additive genetic variation

We can approximate Equation (46) by noticing that the terms *α*_*t,l*_*α*_*s,l*_*p*_*s,l*_(1 − *p*_*s,l*_) are similar to an additive genic variation if effect sizes remain constant through time. We make that assumption here, writing *α*_*l*_ := *α*_*t,l*_ = *α*_*s,l*_, leading to

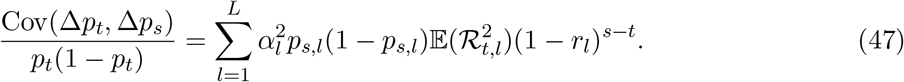

We can further simplify this by assuming that there is no covariance between the additive genic variation at a selected site, and the LD between that selected site and the neutral site. We write the additive genic variation at site *l* at time *s* as 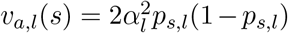, and the average additive genic variation across loci as 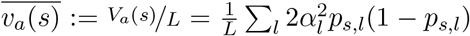. Then, each locus’s additive genic variation can be expressed as: *v*_*a,l*_(*s*) = *v*_*a*_(*s*) + ε_*l*_. Substituting this, the autocovariance is

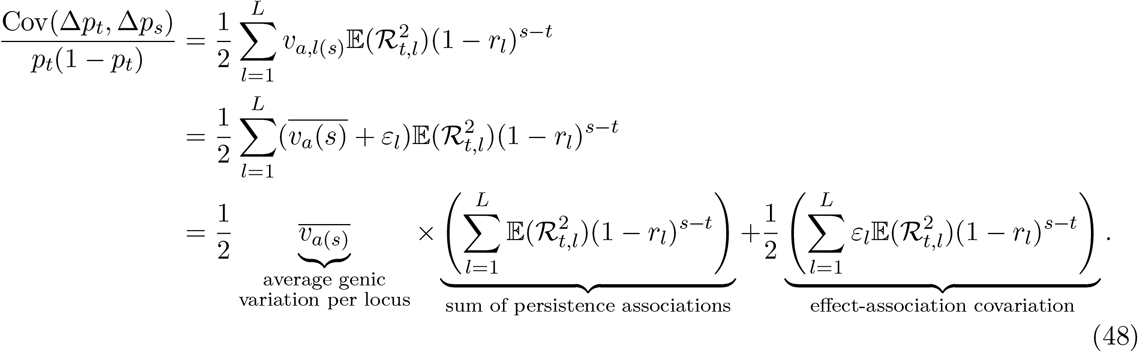

We assume that this last term, which is non-zero in expectation only if there is covariance between the additive genic variation at a selected site and the expected LD between the selected and the neutral sites is zero. Rewriting the total genic variation as *V*_*a*_(*s*),

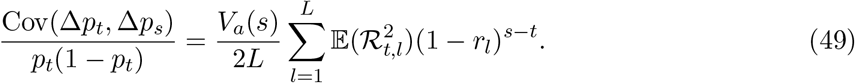

#### Continuous approximation to chromosomes

Currently, we have treated the positions of the selected sites are fixed, conditioning on knowing that a selected site *l* has a recombination fraction *r*_*l*_ away from the focal neutral site site. Here, we make a few further assumptions. First, we assume the *L* selected loci are each independently and identically uniformly distributed along a *continuous* region of *R* Morgans in length (*g* ~ *U*(−*R*/2, *R*/2)), and we now calculate the covariance at a focal neutral site at the origin by taking expectations over the random positions of these sites. Second, we assume that the LD between the neutral and selected site *l* only depends on the recombination fraction between the sites. Since *g* is the genetic distance, the recombination fraction is now provided by a mapping function *r*(*g*) that maps genetic distances to recombination fractions. Throughout the paper and in the simulations, we use Haldane’s mapping function, 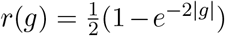 (note the absolute value translates positions on [−*R*/2, *R*/2] to distances to the focal neutral site). Next, we assume the LD between two sites can be completely determined by the distance between the focal neutral site and a random selected site, allowing us to rewrite 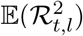 as the function 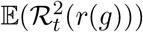. Now, letting 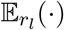 represent the expectation taken over the random positions of selected sites on the genetic map,

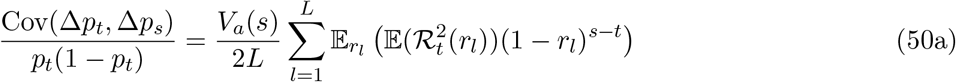

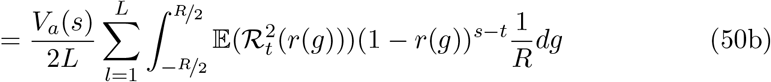

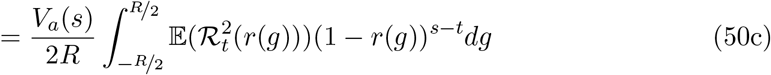

since the 1/*L* cancels with the *L* from the sum of expectations.

As our trait is neutrally evolving before directional selection starts we use the expected neutral LD, 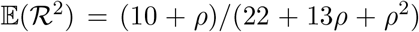 where ρ = 4*Nr*(*g*) (Hill and Robertson 1968; Ohta and Kimura 1969) when *t* is the first generation selection begins.

### A.3 Averaging covariance across multiple loci

Thus far, our covariance assumes that a single neutral site is positioned in the center of a region R Morgans long, with selected sites uniformly distributed along this region. In our simulations, however, we simulate a region that contains many neutral sites which we average over in calculating the temporal autocovariance. In this case, we average over the random distance between a neutral site’s position n and a selected site’s position g, which is *c* = |*n* − *g*|, where *n*, *g* ~ *U*(0, *R*). This random variable *c* is distributed according to the triangle distribution, *f*(*c*) = 2(*R* − *c*)/*R*^2^; we replace the uniform PDF in Equation (50b) with the triangle density PDF and average over the distance between sites,

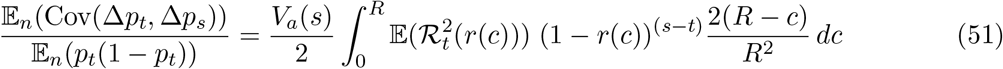

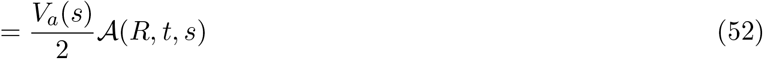

where 𝔼_*n*_(·) indicates we take the expectation also over neutral sites, and we use 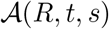 to denote the average linkage disequilibrium between selected and neutral sites that persists from generations *t* to *s* (*t* ≤ *s*). Note that in calculating the standardized covariance above, we use a ratio of expectations rather than the expectation of the ratio (Bhatia et al. 2013).

### A.4 Empirically Calculating the Average LD Persisting Across Generations

In the previous expressions for temporal autocovariance, we stepped through a conceptual model for the average levels of linkage disequilibria between neutral and selected sites that persists across |*s* − *t*| generations 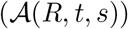, where the positions of selected and neutral sites are randomly distributed along a chromosomal region. In systems with a known recombination map and studies where linkage disequilibria can be calculated, we have the recombination fraction *r*_*i,j*_ and the pairwise linkage disequilibria 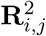 between between two loci *i* and *j* (where **R**^2^ is the M × M matrix of pairwise LD calculated at time *t*). Since we do not *a priori* know whether a site is selected or not, we sum over all polymorphic *M* loci, thus characterizing the average linkage disequilibria in a region as

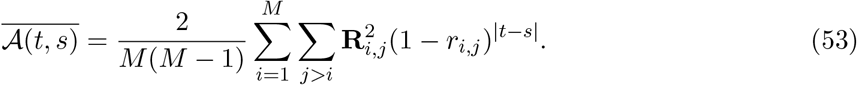

This sum is the empirical analog to the integral in Equation (10).

### A.5 The Strength of Unlinked and Non-gametic Associations

Here, we characterize the contribution of completely genetically unlinked loci segregating for fitness variation to the change in frequency of our neutral allele. Across evolutionary replicates, there is no expected covariance between the neutral allele an individual carries and their fitness (𝔼(Δ*H p*_*t*_) = 𝔼(Cov(*x*_*i*_, *f*_*i*_)) = 0); rather, for unlinked loci, chance associations are created from the variance around this sampling process of neutral alleles into individuals with varying fitness (Var(Δ*H p*_*t*_) = Var(Cov(*x*_*i*_, *f*_*i*_))). As the neutral allele and fitness variation independently assort themselves into individuals, the chance associations that form have a variance given by Var(Cov_*i*_(*x*_*i*_, *f*_*i*_)). This has the form of the sampling variance of a covariance, which for random variables X and Y is given by Kendall et al. (1994, p. 472),

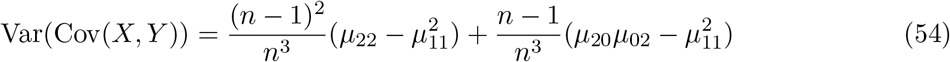

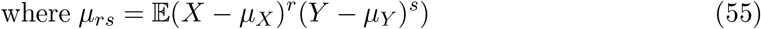

where μ_*X*_ and μ_*Y*_ are the means of *X* and *Y* respectively, and the variance is taken over conceptual replicate populations and the covariance is calculated over the individuals in a population. Then, applying this to our covariance Δ*H p*_1_ = Cov_*i*_(*x*_*i*_, *f*_*i*_),

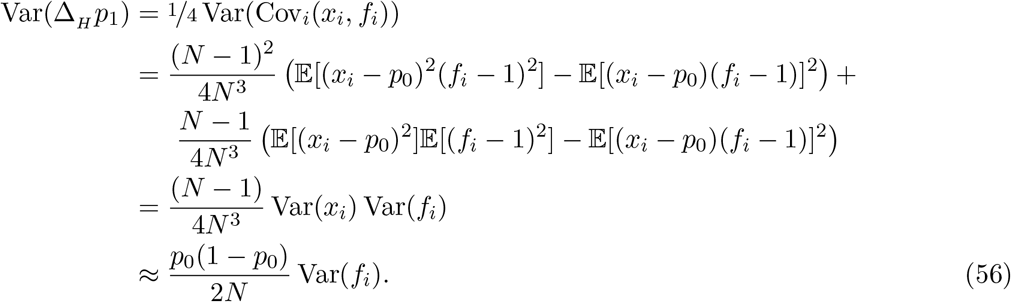

Thus the chance covariances that form between the neutral alleles individuals carry and their fitness have a variance proportional to Var(*f*_*i*_)/2*N*.

#### Non-gametic linkage disequilibrium’s contribution to temporal autocovariance

Throughout the paper, we ignore the effects of non-gametic linkage disequilibria, 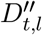, the disequilibria that occurs between the two gametes (maternal and paternal) at two loci (see Figure 8 (A) for an illustration of gametic LD *D*’ and non-gametic LD *D*^’’^). Following Weir’s equation for the sampling variance of non-gametic LD *DA/B* (p. 124, 1996), the chance non-gametic disequilibrium that builds up sampling 2N gametes into N individuals is,

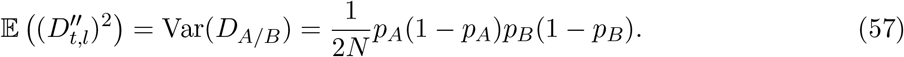

Note the similar form to Equation (56) as both the unlinked and non-gametic LD arise from the random sampling of alleles at different loci into individuals.

There is no expected covariance between our gametic and non-gametic LD within a generation 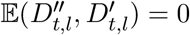, assuming random mating. However, 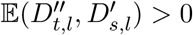 for *s* > *t*, as a fraction of the non-gametic LD may be converted into gametic LD in the next generation. Specifically, following Santiago and Caballero (1995), we can write the product of the non-gametic LD in generation *t* with the gametic LD in generation *s* as

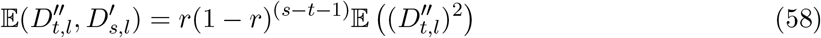

where a proportion *r* the non-gametic LD in generation *t* is converted into gametic LD, and a proportion (1–*r*)^(*s*−*t*−1)^ of this is carried forward unbroken by recombination over the remaining *s*–*t*–1 generations (see Figure 8B for an illustration of this process).

In our analysis in the main text, we ignore these terms as 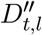 is expected to be small due to its inverse dependence on *N*. However, these terms are necessary for the analysis of looser linkage (Santiago and Caballero 1995, 1998).

### A.6 Connecting our model with the models of Robertson (1961) and Santiago and Caballero (1995, 1998)

Here, we describe the models of Santiago and Caballero (1995, 1998), relating their work on the long-run effective population size experienced by a neutral allele where there is (1) unlinked heritable fitness variation (their 1995 paper), or (2) linkage, where fitness-determining sites are randomly scattered along a chromosome (their 1998 paper). Overall, their models are formulated in a quantitative genetics tradition, where the population genetic dynamics at the selected loci are not explicitly modeled (although these links are made more explicitly in their ’98 paper). In contrast, in deriving our expressions for temporal autocovariance and variances, we use a population genetic approach, modeling the dynamics at selected sites (though we simplify from the full multilocus treatment, e.g. we assume selected loci experience independent sweeps and we ignore the LD between selected sites). We show that we can reconcile the two approaches, and demonstrate that the temporal autocovariance expressions we develop in our models are implicit in their model. We also work through their expressions for *N*_*e*_ with heritable variance, because it represents quite useful result but their original presentation was spread across two papers (and a change in notation).

#### Santiago and Caballero’s 1995 and 1998 models for *N*_*e*_

While our goal in the main text of our paper was to develop expressions for the temporal variances and autocovariances in allele frequency change when there is heritable fitness variation in the population, the goal of both the 1995 and 1998 Santiago and Caballero papers is to derive an expression for the long-run *N*_*e*_ when there is heritable variation for fitness in the population. In their 1995 paper, they find the effective population size for large t (see p. 1018, equation 16) to be

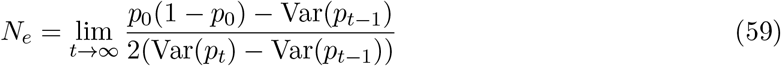

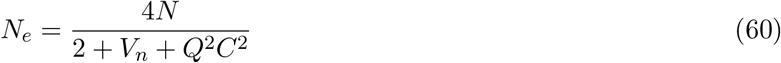

where *C*^2^ is the heritable variation for fitness (*V*_*A*_ in our notation), and *V*_*n*_ is the non-heritable variation in offspring number (i.e. under a Wright–Fisher model, *V*_*n*_ ≈ 2). For a neutral locus completely unlinked from fitness variation (the situation first considered by Robertson 1961), 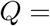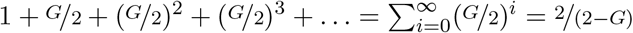 (Santiago and Caballero 1995, equation 17). Note that we’ve simplified their expressions by assuming no assortative mating, and that we try to follow their notation as closely as possible (consequently, the Q here is unrelated to the *Q*_*t,s*_ of the main text). Santiago and Caballero (1995) assume that continual artificial selection maintains a constant level *C*^2^ of fitness variation in the population each generation, yet the *particular* fitness backgrounds the neutral allele is stochastically associated only contributes a fraction G in the next generation, *G*^2^ in the generation after, and so on, as selection reduces genetic variation for fitness (note: in their 1998 paper, they use *Z* instead of *G*). Similarly, the associations between the neutral and fitness backgrounds decay at a rate 1/2 due to independent assortment. Note that Robertson (1961) assumed that the fitness backgrounds that become stochastically associated with the neutral allele do not experience any decay in their fitness variation (*G* = 1); in this case, *Q* = 2 as Robertson’s work found. With an arbitrary amount of linkage between the focal neutral and fitness backgrounds, Santiago and Caballero (1998) show that only *Q* is affected, and derive an expression for *Q* that only depends on *G* and the size of the genome in Morgans, *L* (equation 6, 1998).

In our main text, we model the temporal autocovariance created by heritable fitness variation, which also impacts the cumulative variance in allele frequency change Var(*p*_*t*_ − *p*_0_). To illustrate how our model connects with theirs, below is the cumulative variance in allele frequency change for three generations in their 1995 notation, with the corresponding changes in allele frequency below,

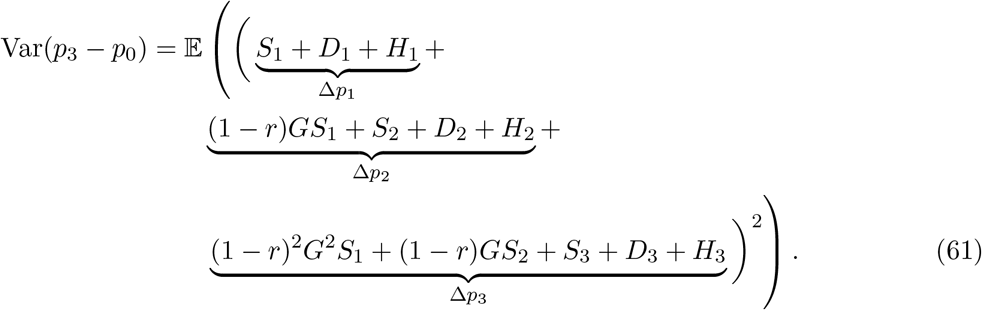

Grouping terms by the generation that the initial association was formed, we see how Santiago and Caballero (1995) define the *Q*_*i*_ terms in their notation,

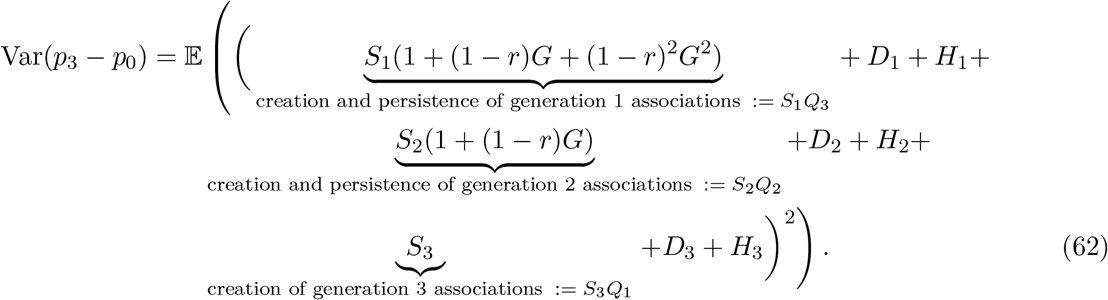

since the associations created in generation *i* (0 < *i* ≤ *t*) persist with probability (1 – *r*)^*t*−*i*^, with proportion *G*^*t*−*i*^ of its original fitness variation in generation *t*. In general, the cumulative impact of the associations formed *i* generations ago has coefficient 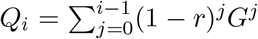. Using these *Q*_*i*_terms simplifies this equation to

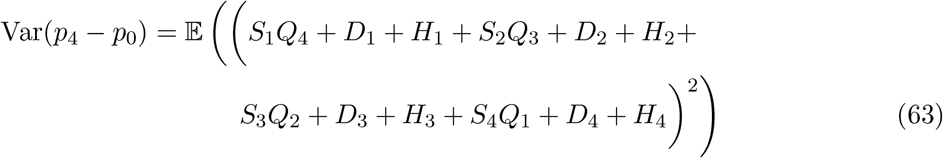

or, in general,

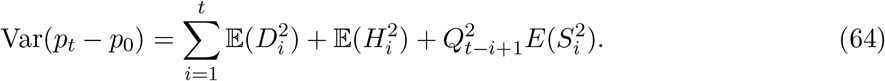

Then, Santiago and Caballero (1995) note that assuming *V*_*n*_, *C*^2^, and population size *N* are constant across generations, the magnitude of all of the effects 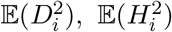, and 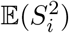 are constant across all generations (for all *i*, so we omit the *i* subscript for these terms), except for a geometric decay due to drift at a rate (1 – 1/2*N*_*e*_) per generation that effects all terms. Such that, when we include the decay in the variance due to drift,

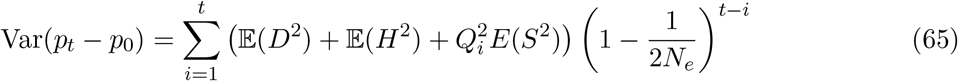

(cf. p. 1018 of Santiago and Caballero 1995). In the long run, the variance in the neutral allele’s frequency change hits a balance. Many copies of the neutral allele segregating in the population are on fitness backgrounds it has recently become stochastically associated with, as segregation and recombination have not broke these associations apart. A few copies of the neutral allele are on fitness backgrounds they became associated with many generations ago, that have by chance survived to remain associated. In all cases, the effect these associations have on present-day allele frequency change is weakened by the fact that natural selection has reduced the genetic variance of these fitness backgrounds. Since the long-run variance in allele frequency under drift in a Wright–Fisher population is

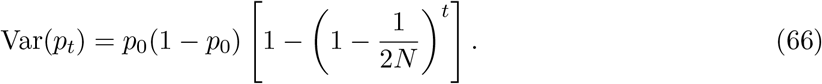

One can estimate the effective population size *N*_*e*_ using the observed difference in variances Var(*p*_*t*_) and Var(*p*_*t*_–_1_). (Note that this is a different long-run effective population size used by others, *N*_*e*_ = *p*_0_(1 – *p*_0_)t/(2 Var(*p*_*t*_ − *p*_0_)), Crow and Kimura 1970) Santiago and Caballero use Equation (66), taking the difference Var(*p*_*t*_)–Var(*p*_*t*_–_1_) and rearranging to end up with the large *t* estimator of *N*_*e*_,

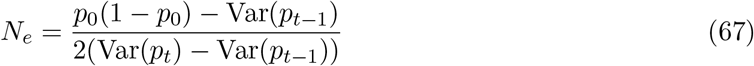

(cf. p. 1018, Santiago and Caballero 1995). Rearranging,

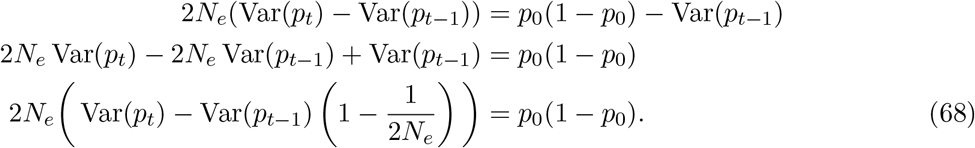

This very conveniently simplifies the sum in Equation (65), as we can show with the case of *t* = 3,

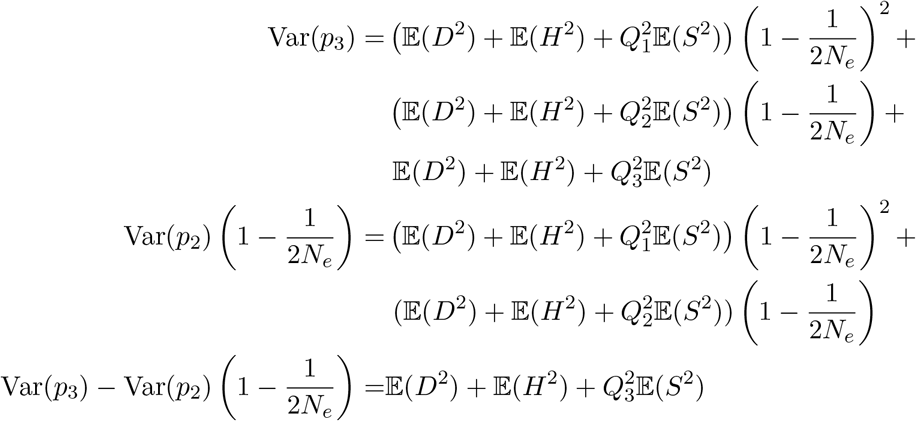

or, generally,

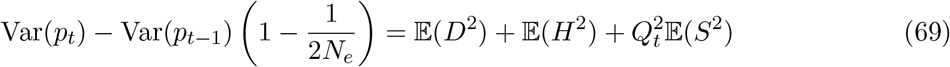

Inserting this into Equation (68), the long-run effective population size can be written as

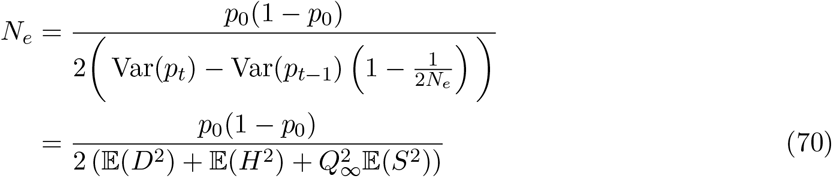

where *Q*_∞_ = 1 + ½ + ¼ + . . . = 2 in Robertson’s (1961) model, and *Q*_∞_ = 1/(1 – *G*(1 –*r*)) in Santiago and Caballero’s (1995) model. (Note: r here represents the recombination fraction between fitness variation and neutral sites, which differs from Santiago and Caballero (1995) equation 17, where r represents the correlation between parental fitness).

Then, Santiago and Caballero (1995) show,

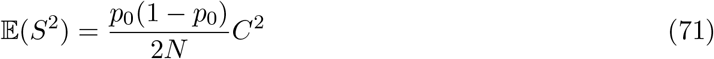

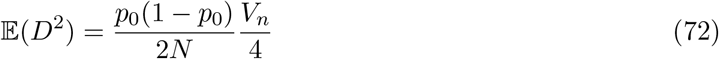

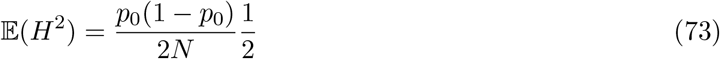

(cf. Santiago and Caballero 1995, equation 11). Inserting these into Equation (70), we have

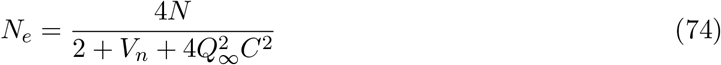

which is a simplified version of Santiago and Caballero’s equation 16, and which further simplifies to *N*_*e*_ = *N* when *V*_*n*_ = 2 and *C*^2^ = 0, the effective population size of a Wright–Fisher population of *N* hermaphroditic individuals.

#### The covariances caused by fitness associations

With an understanding now of the basics of Santiago and Caballero’s (1995, 1998) models, how they connect to our notation, and how they reach their expression for the long-run effective population size, we turn now to finding the temporal autocovariances implicit in their model. We start by looking at the variance in allele frequency between generations 0 and 4 (Equation (61)), including an additional generation so the pattern is clearer later,

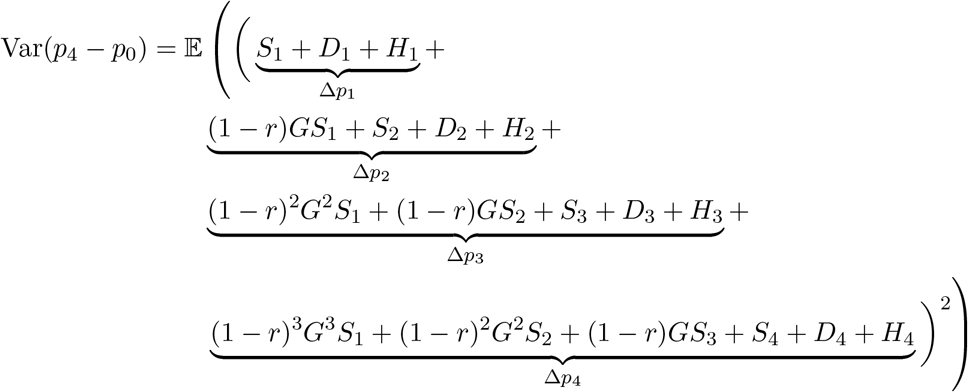

The cross-terms like 𝔼(*D*_1_*D*_2_), 𝔼(*H*_1_*D*_1_) and 𝔼(*S*_1_*S*_2_) are all expected products of independent random variables, where the expectation of each random variable is zero, and consequently are all zero. The only non-zero cross terms are products of 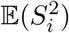. When we look at the covariances with the allele frequency change in the initial generation and a later generation *s*, Cov(Δ_*H*_ *p*_1_, Δ_H_*p*_*s*_),

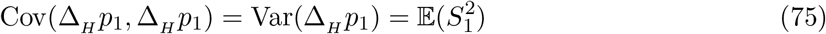

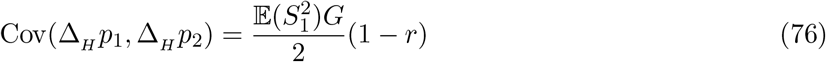

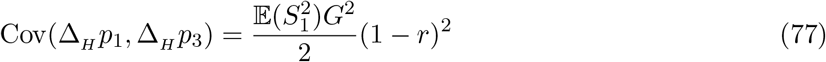

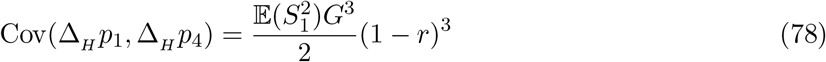

where the 1/2 coefficient comes from the fact that in Santiago and Caballero’s work, the 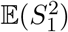 products represent *both* the Cov(Δ_*H*_ *p*_*t*_, Δ_*H*_ *p*_*s*_) and Cov(Δ_*H*_ *p*_*s*_, Δ_*H*_ *p*_*t*_) terms, so a single temporal autocovariance in our notation is half their joint covariance term.

However, if our reference generation is different, say 2, the associations from earlier generations that have persisted to that generation can also lead to covariances to later generations. Looking at the covariances

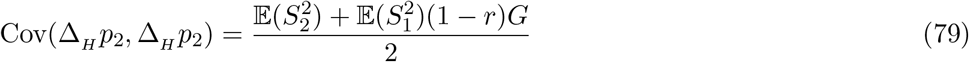

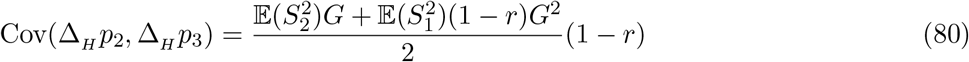

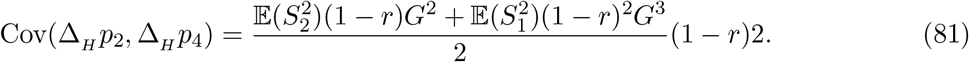

Likewise, the covariances Cov(Δ_*H*_ *p*_3_, Δ_*H*_ *p*_*s*_) include the associations that persist from earlier generations. In general,

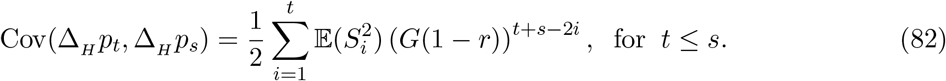

This expression is more complex than our expression for temporal autocovariance because its modeling the LD in generation *t* as it builds up from generations 1 to *t*. In contrast, our expressions for covariance incorporate all of this build up of LD as the initial LD term 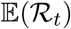 (for a single locus case). This expression for autocovariance implied by Santiago and Caballero’s (1995) work matches our expression for autocovariance (for a single locus) when the arbitrary first generation is *t* rather than 1

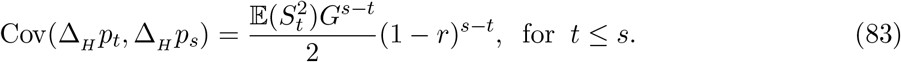

Using the expression for 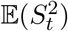 (equivalent to Var(Δ_*H*_ *p*_*t*_) in our notation) derived in Appendix Section A.5,

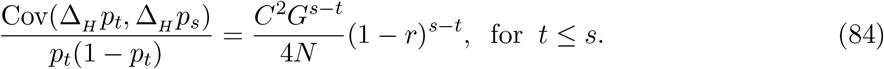

This is analogous to Equation (8) for a single locus, where *C*^2^*G*^*s*−*t*^ is the additive variation in generation s (equivalent to our *V*_*a*_(*s*)), and the factor 1/2*N* represents the chance build up of LD between the neutral site and an unlinked fitness background. In our expression, we condition on existing linkage disequilibrium 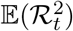 between the neutral site and its fitness background, whereas they assume a buildup of linkage disequilibria to a drift-recombination equilibrium. We can see this by returning to the Δ_*H*_ *p*_4_ term of Equation (61),

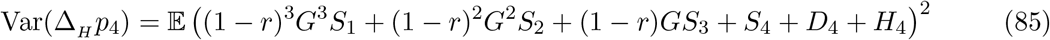

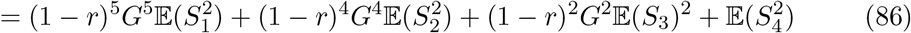

where following Santiago and Caballero’s (1995, p. 1018) approach, we can replace each 𝔼(*S*_*i*_) with 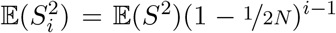 and let *G* = 1 as we focus on the buildup of LD. This gives us the general equation,

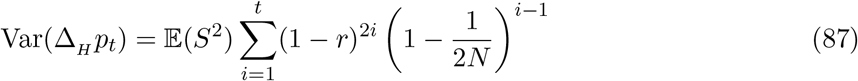

and taking this geometric series to infinity converges (since (1–*r*)^2^(1–1/2*N*) < 1) and replacing 𝔼(*S*^2^) with the chance associations that build up gametes sampled into individuals (Equation (56)),

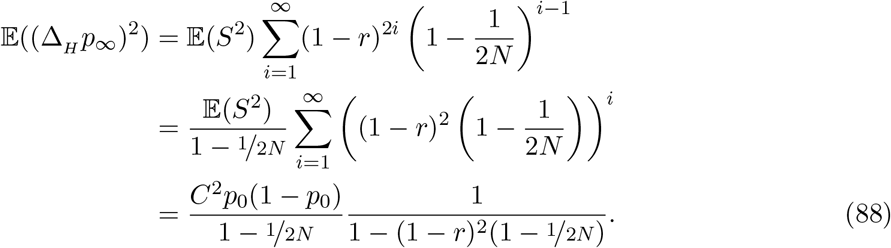

When we assume *r* → 0 and 1/*N* → 0 and *Nr* is a constant, this gives us

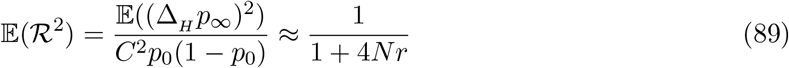

which is analogous to the 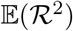 measure of linkage disequilibrium, standardized to rescale the fitness variation. The right-hand side is identical to Sved’s identity by descent equilibrium 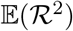 under drift-recombination balance (1971). Note that our expression can be recovered from Equation(82) when the reference generation *t* → ∞ such that the LD hits its equilibrium level,

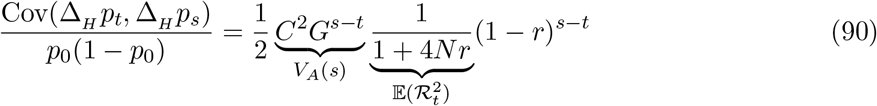

which is identical to our expression for temporal autocovariance when initial LD is due to a neutral drift-recombination balance, and the change in *V*_*A*_ during selection is modeled as a geometric decay at rate *G*. Note that the terms in underbraces indicate the corresponding terms in Equation (8) for a single locus.

### A.7 Multilocus simulation details

#### Targeting an initial level of Additive Genetic Variation

We choose *θ* for the coalescent simulations to target a total number of segregating sites *L* + *M*, where *L* is the number of selected sites (a parameter we vary in our multilocus simulations), and at least *M* = 200 randomly placed neutral sites over which we can calculate the temporal autocovariance. Then, the total number of target sites *M* + *L* is then inflated by a factor of 1.5 to ensure a sufficient number of sites given the random mutation process. The *L* selected sites are randomly chosen from the segregating sites, and all remaining mutations are neutral. Thus, using Watterson’s expression for the expected number of segregating sites under the coalescent (1975), we have *θ* = 1.5(*M* + *L*)/(γ + log(2*N*)) where γ ≈ 0.577 is Euler’s Gamma. Each of the *L* selected sites is given a random effect size of with equal probability, where we choose by targeting a specific level additive genic variation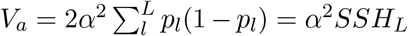 where 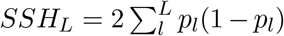 is the sum of the sum of site heterozygosity of the *L* neutrally evolving sites that will be selected once selection begins. Under neutrality, 𝔼(*SSH*_*L*_) = θ_*L*_ = *L*/(γ + log(2*N*)). Then, we set 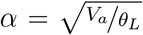. We empirically validate that our target genic variation is close to the empirically observed level.

#### Choosing the simulation parameter range

We simulate over a grid of parameter ranges, varying the number of loci *L*, the target additive genic variation *V*_*a*_ in the region (by varying *α*), and the recombination map length of the region *R* in Morgans. We have varied these parameters over ranges that encompass a wide range likely to be encountered in natural populations. To do this, we found that additive genetic variation for lifetime reproductive success varied over orders of magnitude, from 0 (e.g. in male red deer, Kruuk et al. (2000)) to 1.1 in male red-billed gull (Teplitsky et al. 2009). These values of additive genetic variation are for the entire genome (we write these as *V*_*A,GW*_, where *GW* indicates genome-wide); our simulations model a region of varying map length. We expect most recombination map lengths to be roughly over the scale of 5 50 Morgans in length, and we chose to investigate how temporal autocovariance behaves across a spectrum of recombination, from a completed linked region (*R* = 0), to the scale over which a strong classic hardsweeps could affect diversity (0.5 centiMorgans) to a region where the ends are approximately unlinked (4.5 Morgans), overall giving us a parameter range of *R* ε {0, 0.005, 0.01, 0.05, 0.1, 0.5, 1.5, 4.5}. Over this grid of our region recombination map lengths, and the total organism recombination map lengths, we get a rough estimate of the number of regions we would expect with this level of recombination in the organism’s total genome, imagining homogeneous recombination rate, by taking *G*/*R*. Using this estimate of the number of regions in the genome, we calculate the genetic variation per region over our grid of parameters as *V*_*A,GW*_ *R*/*G*, and target a level of additive genic variation per region *V*_*a*_ equal to this regional additive genetic variation *V*_*A*_. From preliminary simulations, we had found we can not detect much temporal autocovariance below *V*_*a*_ < 0.001 with the initial level of linkage disequilibrium from mutation-drift balance, so we ignore parameters less than this value (other than *V*_*a*_ = 0 as a control). Additionally, to reduce the number of simulations, we exclude *V*_*a*_ > 0.1 as this only excludes a small region of the parameter grid and preliminary simulations demonstrated the behavior of temporal autocovariance with high *V*_*a*_ is evident with the included values. Overall, this gives us a spectrum of target additive genic variation per region of *V*_*a*_ ε {0.001, 0.002, 0.005, 0.01, 0.02, 0.05, 0.08, 0.1}. Note that to prevent our plots from being too dense, we include only a representative subset of this parameter grid in our figures.

### A.8 Accounting for allele frequency sampling noise

In practice, one will calculate the temporal variance-covariance matrix on allele frequency trajectories calculated from sampled chromosomes from the population. We assume a binomial sampling process, where *n* chromosomes are sampled from the population such that 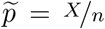, and *X* ~ Binom(*p*, *n*). We can then write 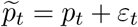, and our covariances can be written as

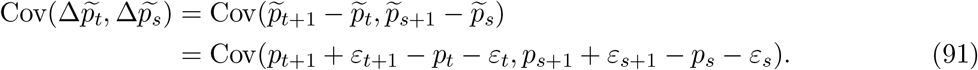

Note this simplifies to

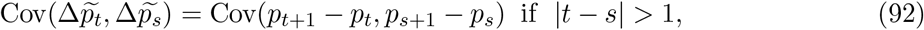

since in these cases, the sampling noise at a timepoint is not shared between the estimated allele frequency changes. However, if |*t*–*s*| = 1 the sampling noise from timepoint *t*] + 1 is shared, biasing the sample estimate of covariance:

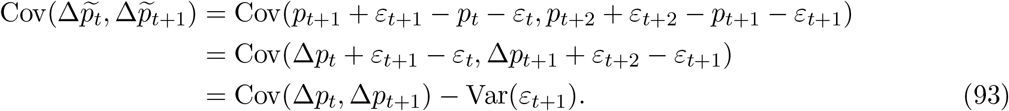

Similarly, the variance (*t* = *s*) is biased, as it is impacted by the binomial sampling noise too,

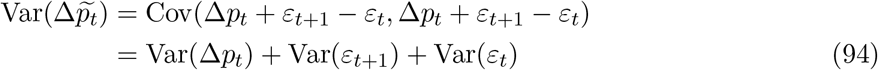

In practice, these covariances are calculated over loci in a region or across the entire genome. We assume that the tracked allele has been randomly swapped (e.g. the tracked allele frequency is not systematically the minor, major, or reference allele), such that 𝔼(Δ*p*_*t,l*_) = 0 for all *t* and *l*. Then, the unbiased covariance estimate as calculated over loci is

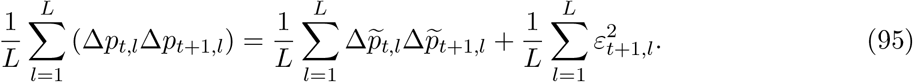

Then, we can use an unbiased plugin estimate of the frequency sampling variance 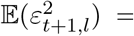 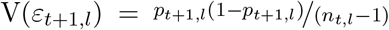 (Nei 1987, p. 191) to estimate these bias terms, and add or subtract them from the estimator accordingly. Accounting for finite sampling, the unbiased sample variance-covariance matrix now has elements:

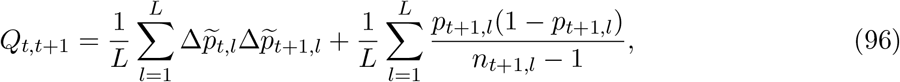

and variance

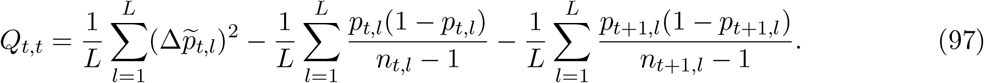

In Section 2.5, we used population frequencies in introducing the method of moments estimators of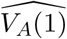and 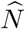. Here, we discuss the performance of these estimators with sample allele frequencies. Our simulations are identical to those described in Section 2.4, except we have increased the target number of neutral sites in each region so it is around 10, 000. We mimic sampling of *n* = {50, 100, 200, 500} chromosomes from the population, and use the bias-corrected sample variance-covariance matrix in the method-of-moments approach described in Section 3 to estimate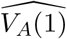 and 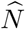.

In Figure 1, we show the performance of our estimators in the case where chromosomes *n* = 100 have been sampled from the population. Overall, there are two important differences compared with Figure 5 of the main text. First, while the estimator 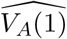 performs well for high levels around *V*_*A*_ ~ 0.1, the variance around the estimator increases significantly as *V*_*A*_ grows weaker. As the covariances are proportional to *VA*/*R*, sampling noise grows larger than the theoretic temporal autocovariance as *V*_*A*_ become weaker. Then, one cannot discriminate against the chance covariances formed by the sampling process from the temporal autocovariance created by linked selection without either a large sample size or more timepoints. Second, the approach underestimates *V*_*A*_(1) for very loose linkage (*R* > 1/2). This is another consequence of the first problem; as sampling noise grows equal, or larger than the magnitude of temporal autocovariance, the estimation procedure performs poorly. As the linkage becomes more loose, the magnitude of temporal autocovariance grows weak relative to the sampling noise, and this noise can be partially absorbed by 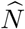. This effect can be somewhat ameliorated by calculating the sample variance-covariance matrix over a shorter region of the genome such that *R* is smaller (as long as SNP density is sufficient), or by increasing the sample size. Finally, the estimation of effective population size shown in Figure 1 are also affected by *V*_*A*_, as discussed in Section 3 (though the effect is obscured by the sampling noise): high levels of *V*_*A*_ in regions with low recombination generally lead to underestimates of 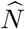.

**Figure 1:**
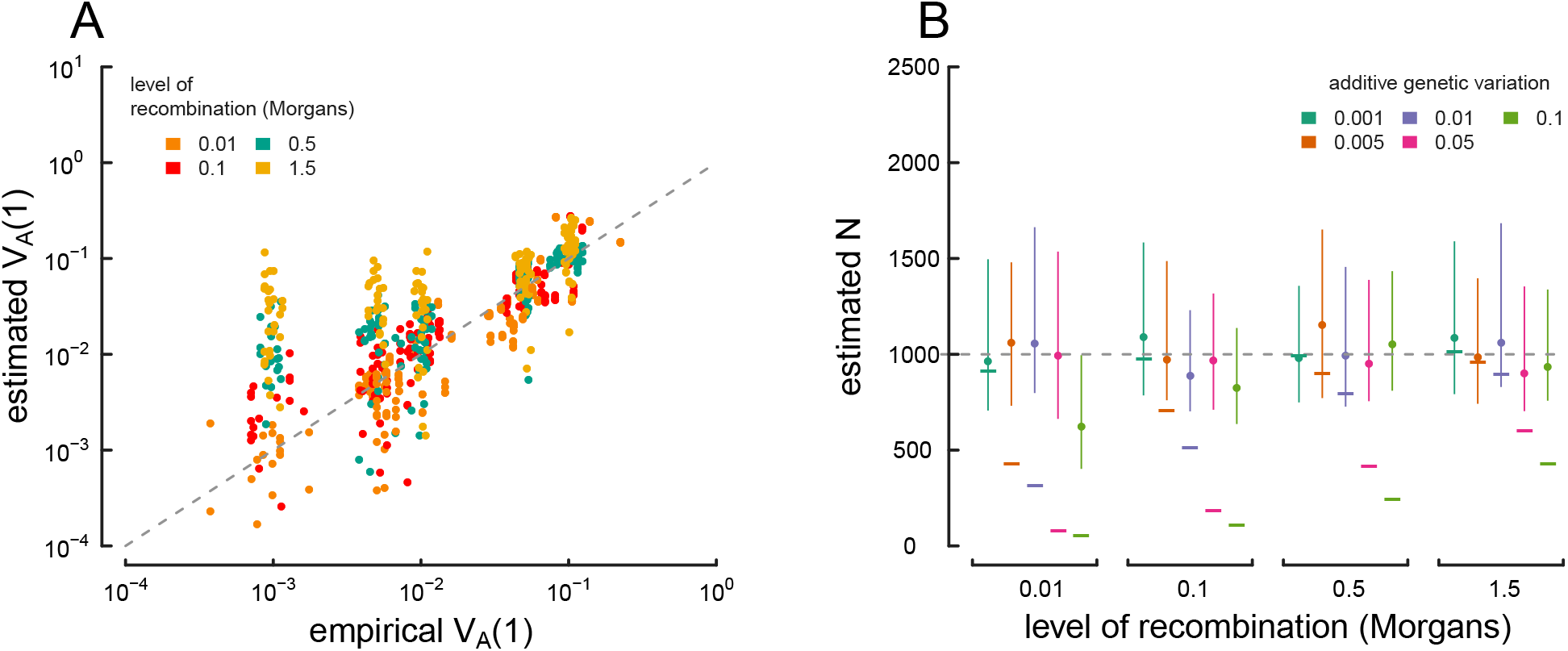
True parameter values and estimates using the method-of-moments approach on sample (n = 100 chromosomes) multilocus simulation data; these figures are analogous to Figure 5 in the main text, except the estimators have been calculated on sample, rather than population frequency data. (A) The true *V*_*A*_(1) (x-axis) and 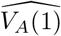 estimated from the sample variance/covariance matrix (y-axis) for each simulation replicate across different levels of recombination (indicated by each point’s color). (B) Estimated drift-effective population size 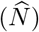 across a range of simulations with different levels of additive genetic variance and recombination. Each point denotes the median, with lines denoting the interquartile range. A simple temporal estimate of the effective population size, estimated with accounting for the effects of selection, is averaged for each replicate and plotted as a dash. The true value (N = 1, 000) is shown with the dashed gray line.

To understand how sample size affects the method-of-moments estimators, Figure 2 depicts median relative error of 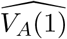 and 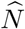 for various sample sizes.

**Figure 2:**
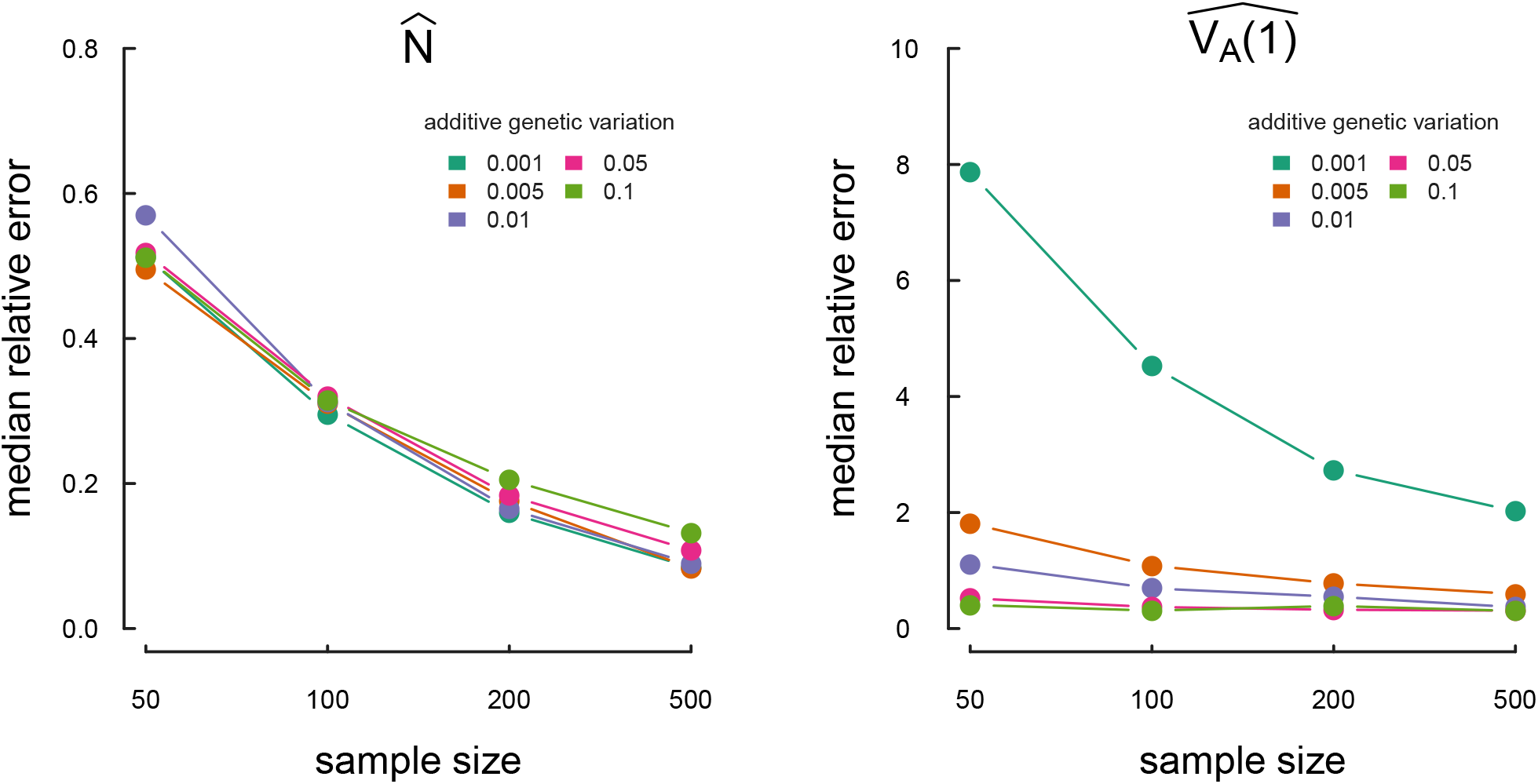
The median relative estimation error, over 30 replicate simulations, of the method of moments estimator for drift effective population size 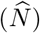 and the initial additive genetic variance 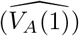.

## B Appendix Figures

### B.1 Validation of simulation routine

**Figure 3:**
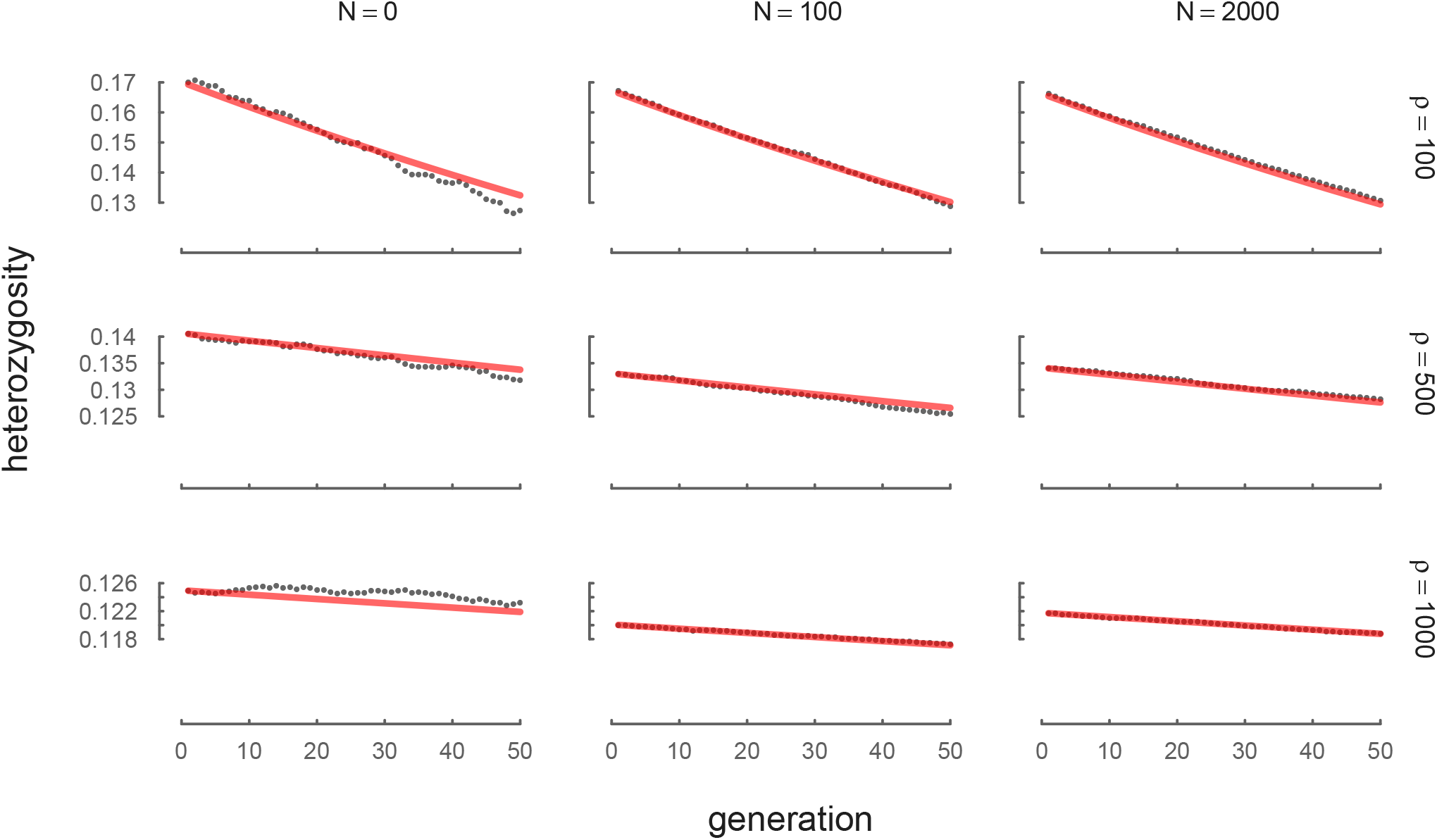
The neutral decay of heterozygosity due to drift, averaged across 100 replicates and a variety of *N* and ρ levels. The red line is the theoretic expectation *H*_*t*_ = *H*_1_(1 − 1/(2N))^*t−*1^.

**Figure 4:**
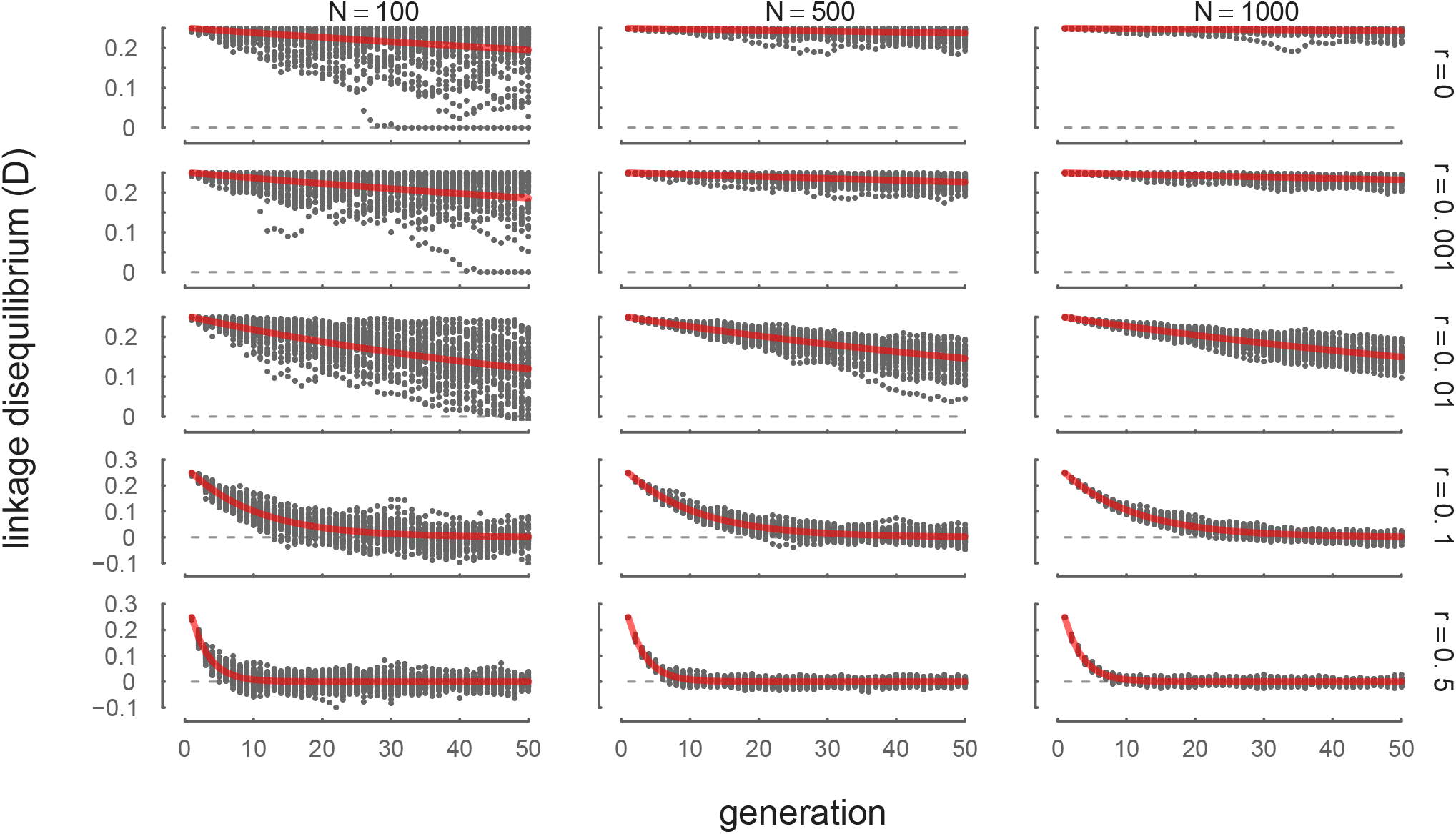
The decay of LD between neutral sites due to recombination, across 100 replicates and a variety of *V*_*A*_ and R levels. The initial population is created to have an artificial level of initial LD, with half the gametes carrying all derived alleles, and the other half carrying all ancestral alleles. The red line is the theoretic expectation *D*_*t*_ = *D*_1_(1 − 1/(2N))^*t−*1^(1 − r)^*t−*1^.

**Figure 5:**
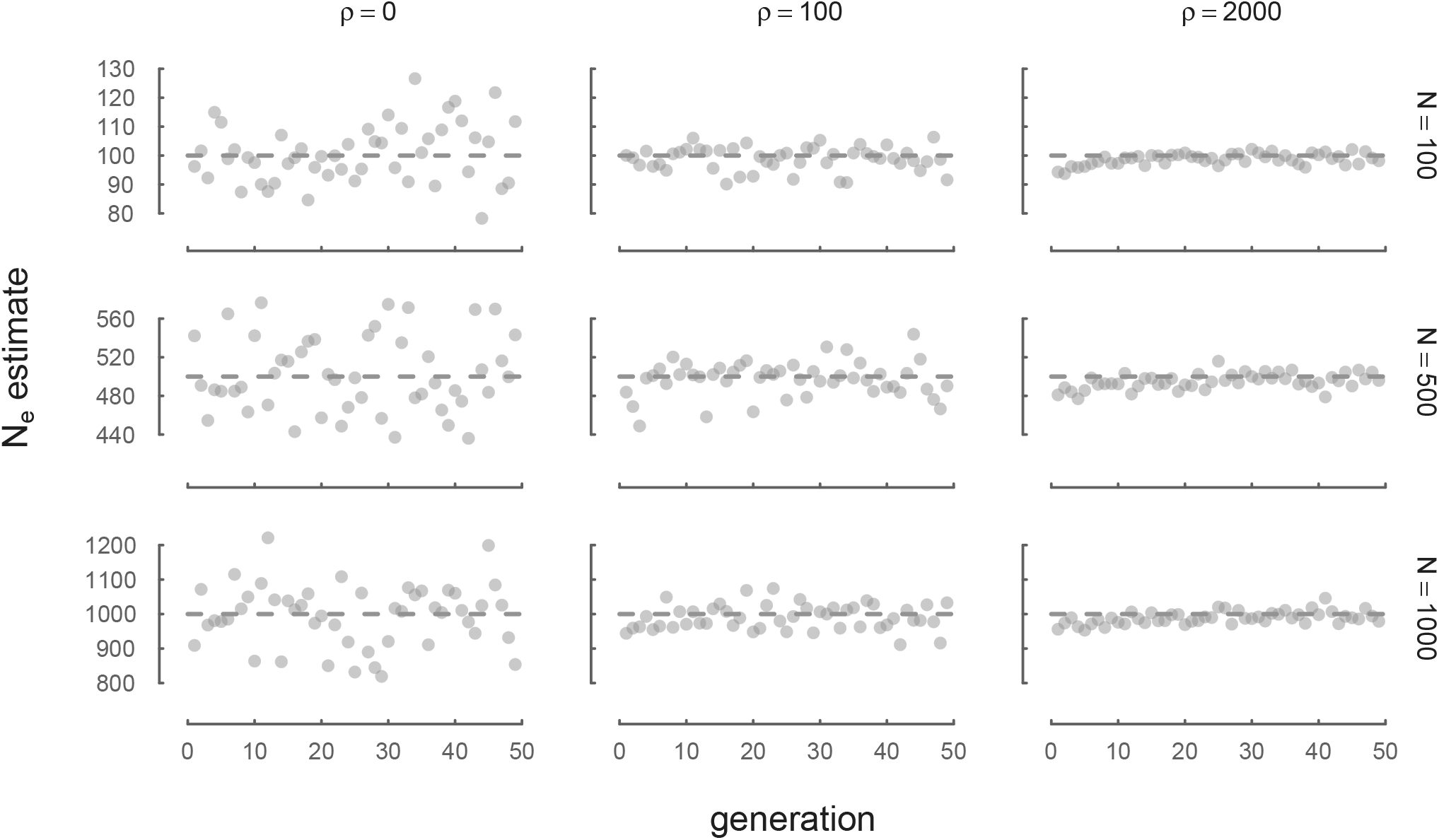
N_*e*_ estimated through time from neutral forward simulations is consistent with true *N* across a variety of recombination ρ and *N* parameters. Here, we estimate *N*_*e*_ with 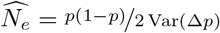 from neutral allele frequency changes.

### B.2 Dynamics of Variances

**Figure 6:**
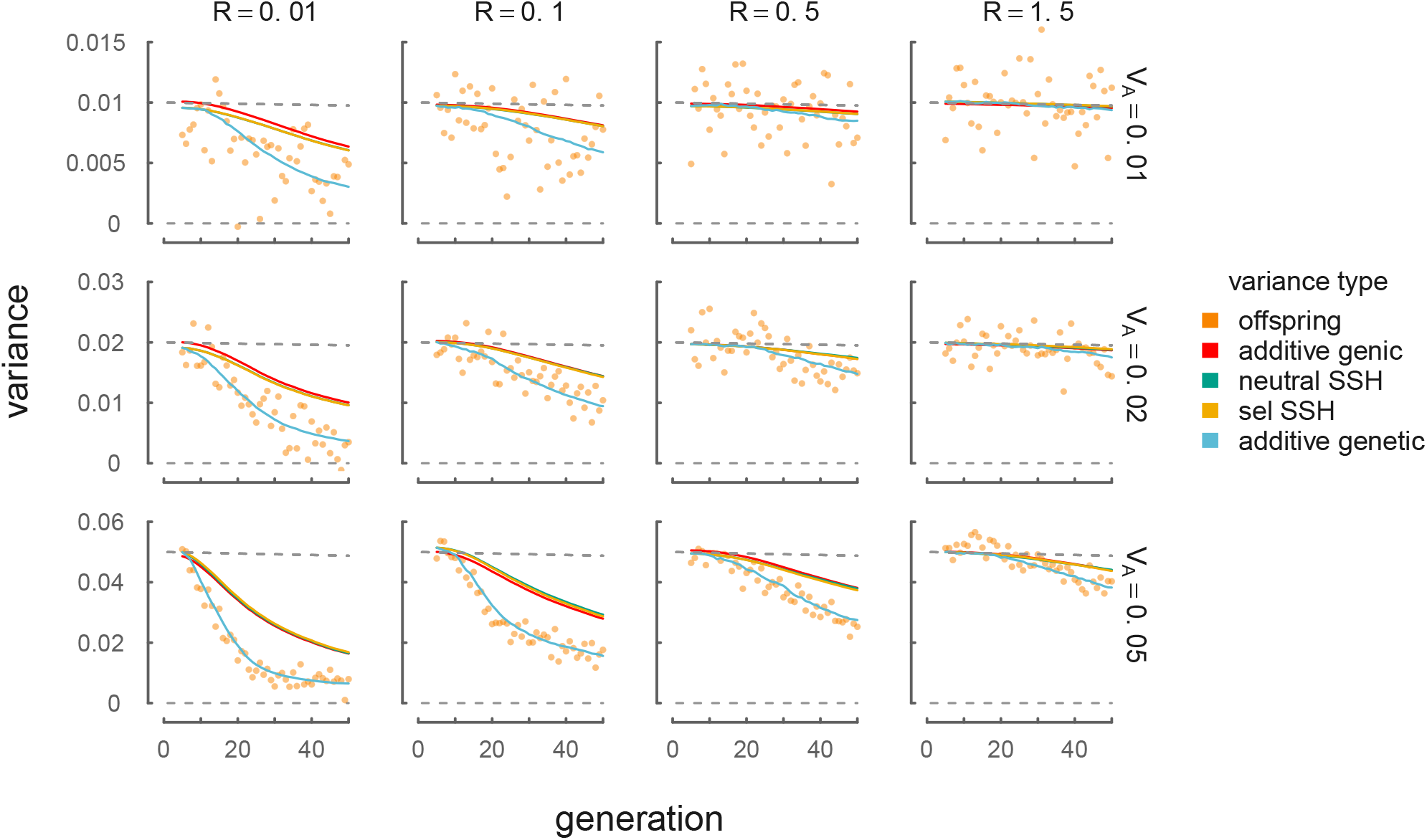
The dynamics of different variances in our simulations, across a variety of initial target *V*_*A*_ and R levels. Orange points represent the empirical heritable variance in offspring (e.g. with the noise of the Wright–Fisher reproduction process removed). These closely track the additive genetic variance for the trait undergoing directional selection, *V*_*A*_ = Var(z). The red line shows the dynamics of additive genic variance, 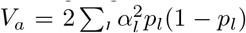, which is closely tracked by both the selected (yellow line) and neutral (green line)

### B.3 Supplementary Temporal Autocovariance Figures

**Figure 7:**
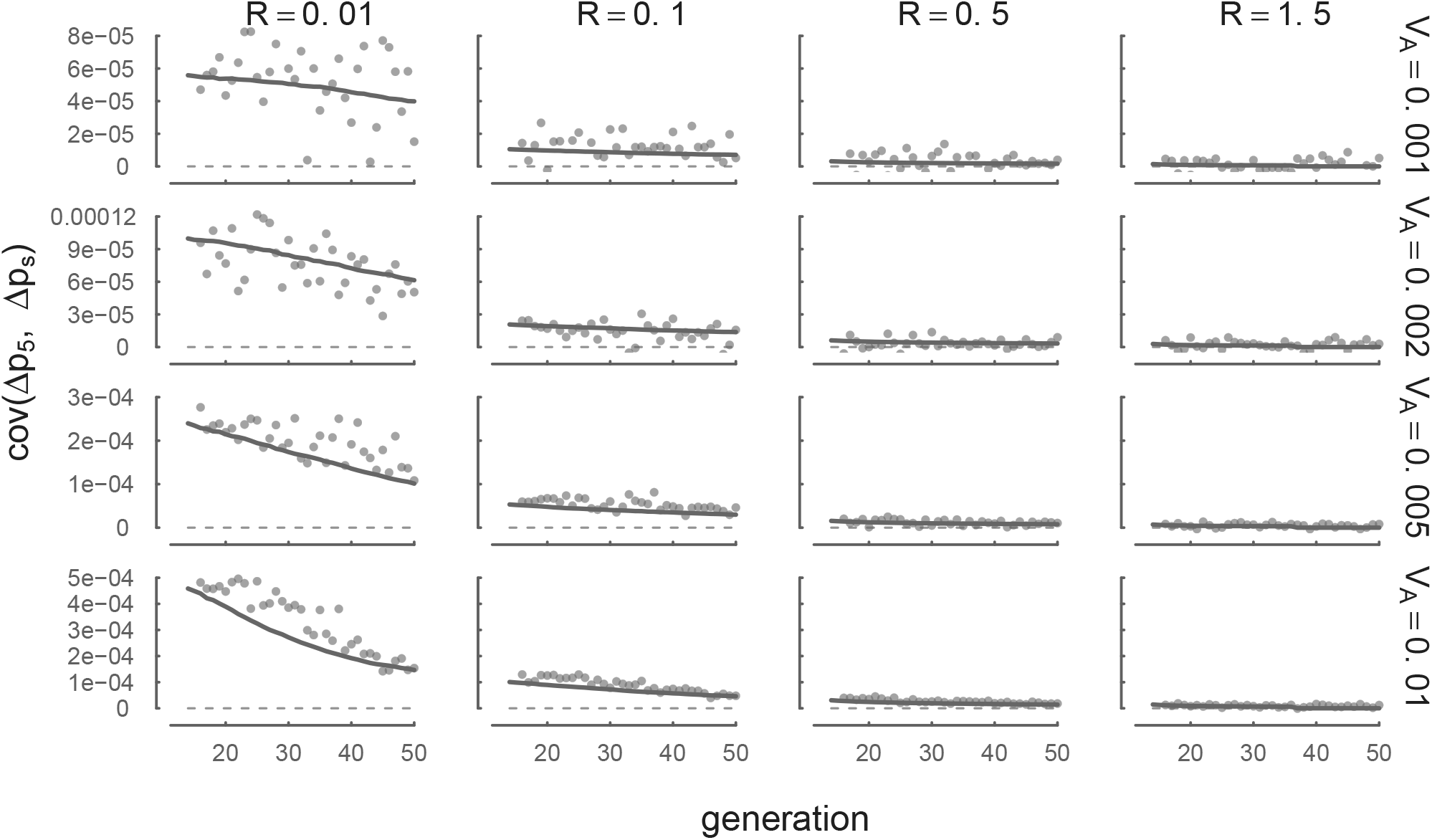

Each panel shows the temporal autocovariance Cov(Δp_13_, Δp_*s*_) on the y-axis, where s varies along the x-axis. This is analogous to Figure 2 with a different reference generation (t = 13), and compares the averaged simulation results (points) with the temporal autocovariance predicted by Equation (10) using the empirical additive genetic variance (curve). The covariances in these panels are weaker compared to those in Figure 2 because by generation 13, additive genetic variance for fitness and the linkage disequilibria between neutral and selected sites has decayed.

**Figure 8:**
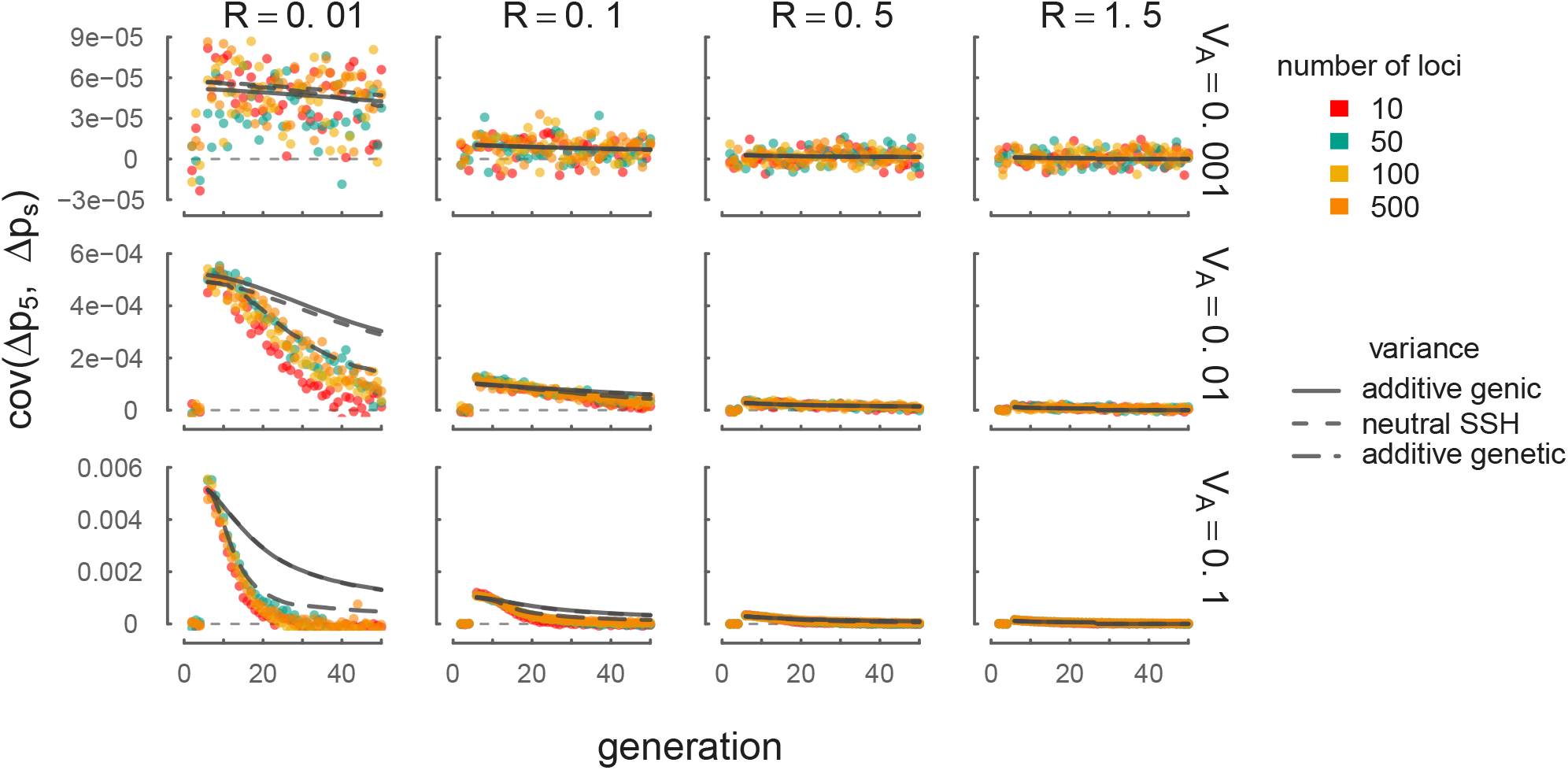

A version of Figure 2 with a subset of *V*_*A*_ parameters used in our simulations that vary over orders of magnitude. This demonstrates that our theory using the empirical additive genic variance (solid gray line), additive genetic variance (long dashed gray line), and the neutral SSH proxy (short dashed gray lines) performs as described in Section 2.5 even when *V*_*A*_ varies over orders of magnitude in a region. Higher variance in the empirical covariances with weak selection (V_*A*_ = 0.001) are due to chance covariances due to drift. The light gray dashed line depicts Cov(Δp_5_, Δp_*s*_) = 0.

**Figure 9:**
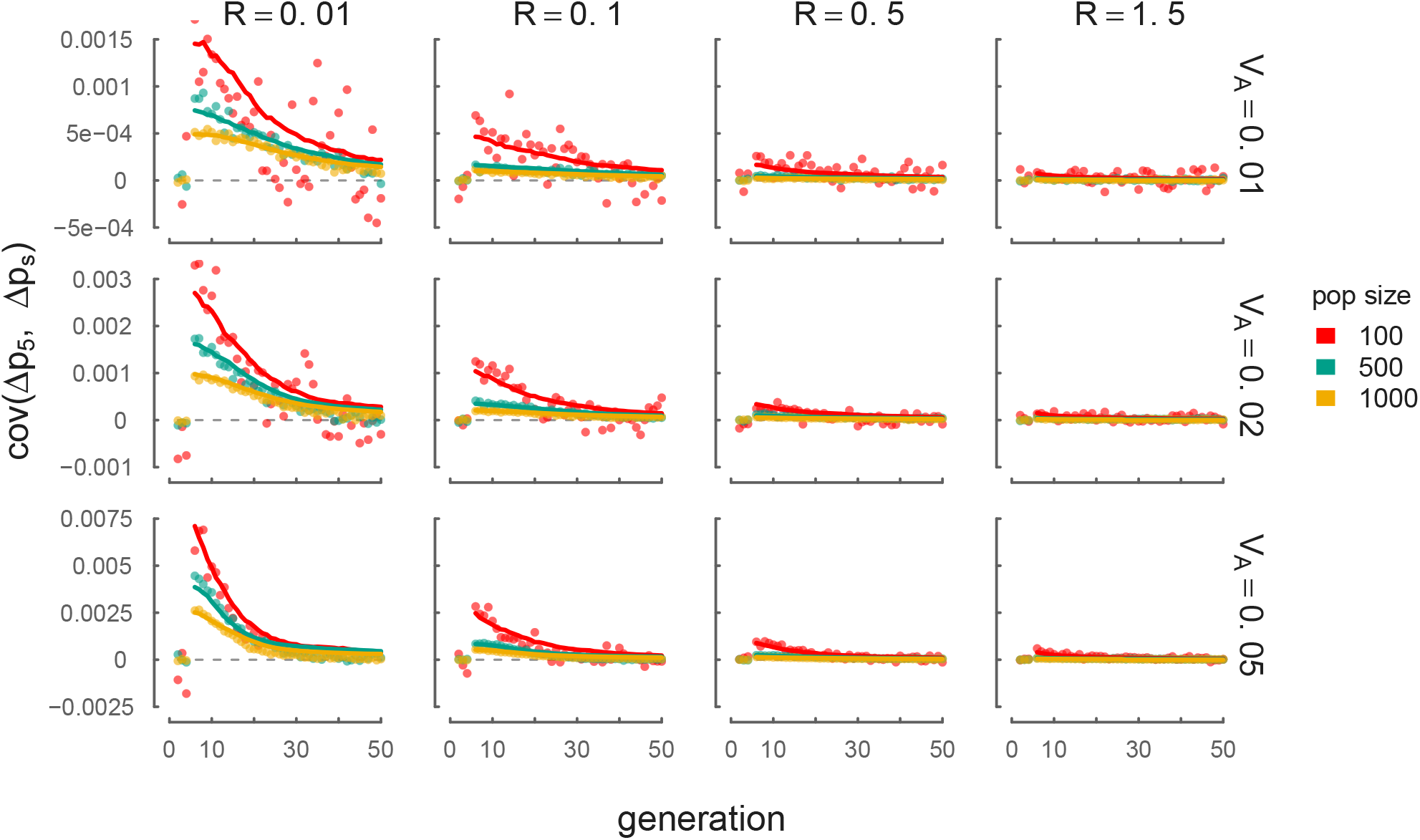

A version of Figure 2 demonstrating temporal autocovariance simulation results and theoretic predictions with varying *N*. This demonstrates that our theory using the empirical additive genetic variance (lines) fits simulations across a variety of *N* parameters. The light gray dashed line depicts Cov(Δp_5_, Δp_*s*_) = 0. Note that the initial LD varies due to differing equilibrium levels of LD from our burnin across varying *N*.

## References

Aguade, M, N Miyashita, and C H Langley (1989). “Reduced variation in the yellow-achaete-scute region in natural populations of Drosophila melanogaster”. en. In: Genetics 122.3, pp. 607–615.

Barghi, Neda et al. (2019). “Genetic redundancy fuels polygenic adaptation in Drosophila”. en. In: PLoS Biol. 17.2, e3000128.

Barton, N H (2000). “Genetic hitchhiking”. In: Proceedings of the Royal Society of London B: Biological Sciences 355.1403, pp. 1553–1562.

Barton, N H and Sarah P Otto (2005). “Evolution of recombination due to random drift”. en. In: Genetics 169.4, pp. 2353–2370.

Barton, N H and M Turelli (1987). “Adaptive landscapes, genetic distance and the evolution of quantitative characters”. en. In: Genet. Res. 49.2, pp. 157–173.

Barton, N H and M Turelli (1991). “Natural and sexual selection on many loci”. In: Genetics 127.1, pp. 229–255.

Begun, D J and C F Aquadro (1992). “Levels of naturally occurring DNA polymorphism correlate with recombination rates in D. melanogaster”. en. In: Nature 356.6369, pp. 519–520.

Begun, David J et al. (2007). “Population genomics: whole-genome analysis of polymorphism and divergence in Drosophila simulans”. en. In: PLoS Biol. 5.11, e310.

Beissinger, Timothy M et al. (2016). “Recent demography drives changes in linked selection across the maize genome”. en. In: Nat Plants 2, p. 16084.

Bergland, Alan O, Emily L Behrman, Katherine R O’Brien, Paul S Schmidt, and Dmitri A Petrov (2014). “Genomic Evidence of Rapid and Stable Adaptive Oscillations over Seasonal Time Scales in Drosophila”. In: PLoS Genet. 10.11, e1004775–19.

Bhatia, Gaurav, Nick Patterson, Sriram Sankararaman, and Alkes L Price (2013). “Estimating and interpreting FST: the impact of rare variants”. en. In: Genome Res. 23.9, pp. 1514–1521.

Bulik-Sullivan, Brendan K et al. (2015). “LD Score regression distinguishes confounding from poly-genicity in genome-wide association studies”. en. In: Nat. Genet. 47.3, pp. 291–295.

Bulmer, M G (1971). “The Effect of Selection on Genetic Variability”. In: Am. Nat. 105.943, pp. 201–211.

Bulmer, Michael George (1980). The Mathematical Theory of Quantitative Genetics. Clarendon Press.

Bürger, R (2000). The Mathematical Theory of Selection, Recombination, and Mutation. John Wiley & Sons.

Burke, Molly K et al. (2010). “Genome-wide analysis of a long-term evolution experiment with Drosophila”. In: Nature 467.7315, pp. 587–590.

Burri, Reto (2017). “Interpreting differentiation landscapes in the light of long-term linked selection: Differentiation And Long-Term Linked Selection”. In: Evolution Letters 1.3, pp. 118–131.

Burt, Austin (1995). “The Evolution of Fitness”. en. In: Evolution 49.1, pp. 1–8.

Cannings, C (1974). “The latent roots of certain Markov chains arising in genetics: a new approach, I. Haploid models”. In: Adv. Appl. Probab. pp. 260–290.

Charlesworth, Brian (2009). “Fundamental concepts in genetics: effective population size and patterns of molecular evolution and variation”. In: Nature Publishing Group 10.3, pp. 195–205.

Charlesworth, Brian (2015). “Causes of natural variation in fitness: evidence from studies of Drosophila populations”. en. In: Proc. Natl. Acad. Sci. U. S. A. 112.6, pp. 1662–1669.

Charlesworth, Brian, M T Morgan, and D Charlesworth (1993). “The effect of deleterious mutations on neutral molecular variation”. In: Genetics 134.4, pp. 1289–1303.

Chevin, Luis-Miguel and Frédéric Hospital (2008). “Selective sweep at a quantitative trait locus in the presence of background genetic variation”. en. In: Genetics 180.3, pp. 1645–1660.

Christensen, Ronald (2011). Plane Answers to Complex Questions | SpringerLink. Springer Science & Business Media.

Coop, Graham (2016). Does linked selection explain the narrow range of genetic diversity across species? Tech. rep. Cold Spring Harbor Labs Journals.

Corbett-Detig, Russell B, Daniel L Hartl, and Timothy B Sackton (2015). “Natural Selection Constrains Neutral Diversity across A Wide Range of Species”. In: PLoS Biol. 13.4, e1002112–25.

Crouch, Daniel J M (2017). “Statistical aspects of evolution under natural selection, with implications for the advantage of sexual reproduction”. en. In: J. Theor. Biol. 431, pp. 79–86.

Crow, James Franklin and Motoo Kimura (1970). An Introduction to Population Genetics Theory. New York, Evanston and London: Harper & Row, Publishers.

Cutter, Asher D and Bret A Payseur (2013). “Genomic signatures of selection at linked sites: unifying the disparity among species”. In: Nature Publishing Group 14.4, pp. 262–274.

Dobzhansky, Theodosius (1943). “Genetics of Natural Populations IX. Temporal Changes in the Composition of Populations of Drosophila Pseudoobscura”. In: Genetics 28.2, pp. 162–186.

Dobzhansky, Theodosius (1971). “Evolutionary Oscillations in Drosophila pseudoobscura”. In: Ecological Genetics and Evolution. Boston, MA: Springer US, pp. 109–133.

Edmonds, Christopher A, Anita S Lillie, and L Luca Cavalli-Sforza (2004). “Mutations arising in the wave front of an expanding population”. en. In: Proc. Natl. Acad. Sci. U. S. A. 101.4, pp. 975–979.

Elyashiv, Eyal et al. (2016). “A Genomic Map of the Effects of Linked Selection in Drosophila”. In: PLoS Genet. 12.8, e1006130–24.

Endler, John A (1986). Natural selection in the wild. Princeton University Press.

Excoffier, Laurent and Nicolas Ray (2008). “Surfing during population expansions promotes genetic revolutions and structuration”. en. In: Trends Ecol. Evol. 23.7, pp. 347–351.

Feder, A F, S Kryazhimskiy, and J B Plotkin (2014). “Identifying signatures of selection in genetic time series”. In: Genetics.

Fisher, R A and E B Ford (1947). “The spread of a gene in natural conditions in a colony of the moth Panaxia dominula”. In: Heredity 1, pp. 143–174.

Franssen, S U, R Kofler, and C Schlötterer (2017). “Uncovering the genetic signature of quantitative trait evolution with replicated time series data”. en. In: Heredity 118.1, pp. 42–51.

Fu, Qiaomei et al. (2016). “The genetic history of Ice Age Europe”. en. In: Nature 534.7606, pp. 200–205.

Gillespie, J H (2000). “Genetic drift in an infinite population. The pseudohitchhiking model”. en. In: Genetics 155.2, pp. 909–919.

Gillespie, John H (2001). “Is the population size of a species relevant to its evolution?” In: Evolution 55.11, pp. 2161–2169.

Gingerich, P D (1983). “Rates of evolution: effects of time and temporal scaling”. en. In: Science 222.4620, pp. 159–161.

Good, Benjamin H, Michael J McDonald, Jeffrey E Barrick, Richard E Lenski, and Michael M Desai (2017). “The dynamics of molecular evolution over 60,000 generations”. en. In: Nature 551.7678, pp. 45–50.

Good, Benjamin H, Aleksandra M Walczak, Richard A Neher, and Michael M Desai (2014). “Genetic Diversity in the Interference Selection Limit”. In: PLoS Genet. 10.3, e1004222.

Haldane, J B S (1919). “The combination of linkage values and the calculation of distances between the loci of linked factors”. In: J. Genet. 8.29, pp. 299–309.

Hallatschek, Oskar and David R Nelson (2008). “Gene surfing in expanding populations”. en. In: Theor. Popul. Biol. 73.1, pp. 158–170.

Hansen, Lars Peter (1982). “Large Sample Properties of Generalized Method of Moments Estimators”. In: Econometrica 50.4, pp. 1029–1054.

Hendry, A P and M T Kinnison (1999). “Perspective: the pace of modern life: measuring rates of contemporary microevolution”. In: Evolution 53.6, p. 1637.

Hendry, Andrew P, Daniel J Schoen, Matthew E Wolak, and Jane M Reid (2018). “The Contemporary Evolution of Fitness”. en. In: Annu. Rev. Ecol. Evol. Syst.

Hermisson, J and P S Pennings (2005). “Soft sweeps: Molecular population genetics of adaptation from standing genetic variation”. In: Genetics.

Hernandez, R D, J L Kelley, E Elyashiv, and S C Melton (2011). “Classic selective sweeps were rare in recent human evolution”. In: Science.

Heyer, Evelyne, Alexandre Sibert, and Frédéric Austerlitz (2005). “Cultural transmission of fitness: genes take the fast lane”. en. In: Trends Genet. 21.4, pp. 234–239.

Hill, W G and Alan Robertson (1966). “The effect of linkage on limits to artificial selection”. In: Genet. Res. 8.03, pp. 269–294.

Hill, W G and Alan Robertson (1968). “Linkage disequilibrium in finite populations”. In: Theor. Appl. Genet. 38.6, pp. 226–231.

Höllinger, Ilse, Pleuni S Pennings, and Joachim Hermisson (2019). “Polygenic adaptation: From sweeps to subtle frequency shifts”. en. In: PLoS Genet. 15.3, e1008035.

Hudson, R R (1994). “How can the low levels of DNA sequence variation in regions of the drosophila genome with low recombination rates be explained?” en. In: Proc. Natl. Acad. Sci. U. S. A. 91.15, pp. 6815–6818.

Hudson, R R and N L Kaplan (1995). “Deleterious background selection with recombination”. In: Genetics 141.4, pp. 1605–1617.

J A Woolliams, N R Wray, R Thompson (2008). “Prediction of long-term contributions and in-breeding in populations undergoing mass selection”. In: 62, pp. 231–242.

Jain, Kavita and Wolfgang Stephan (2015). “Response of Polygenic Traits Under Stabilizing Selection and Mutation When Loci Have Unequal Effects”. en. In: G3 5.6, pp. 1065–1074.

Jain, Kavita and Wolfgang Stephan (2017). “Rapid Adaptation of a Polygenic Trait After a Sudden Environmental Shift”. In: Genetics 206.1, pp. 389–406.

Johansson, Anna M, Mats E Pettersson, Paul B Siegel, and Örjan Carlborg (2010). “Genome-Wide Effects of Long-Term Divergent Selection”. In: PLoS Genet. 6.11, e1001188–12.

Kaplan, N L, R R Hudson, and C H Langley (1989). “The “hitchhiking effect” revisited”. In: Genetics.

Keinan, Alon and David Reich (2010). “Human population differentiation is strongly correlated with local recombination rate”. en. In: PLoS Genet. 6.3, e1000886.

Kelleher, Jerome, Alison M Etheridge, and Gilean McVean (2016). “Efficient Coalescent Simulation and Genealogical Analysis for Large Sample Sizes”. en. In: PLoS Comput. Biol. 12.5, e1004842.

Kendall, Maurice, Alan Stuart, J Keith Ord, and A O’Hagan (1994). “Kendall’s advanced theory of statistics, volume 1: Distribution theory”. In: Arnold, sixth edition edition.

Kettlewell, H B D (1958). A survey of the frequencies of Biston betularia (L.)(Lep.) and its melanic forms in Great Britain. Vol. 12. Heredity. Nature Publishing Group.

Kettlewell, H B D (1961). “The phenomenon of industrial melanism in Lepidoptera”. In: Annu. Rev. Entomol.

Kinnison, M T and A P Hendry (2001). “The pace of modern life II: from rates of contemporary microevolution to pattern and process”. en. In: Genetica 112-113, pp. 145–164.

Kopp, Michael and Joachim Hermisson (2007). “Adaptation of a quantitative trait to a moving optimum”. In: Genetics 176.1, pp. 715–719.

Kopp, Michael and Joachim Hermisson (2009a). “The genetic basis of phenotypic adaptation I: fixation of beneficial mutations in the moving optimum model”. en. In: Genetics 182.1, pp. 233–249.

Kopp, Michael and Joachim Hermisson (2009b). “The genetic basis of phenotypic adaptation II: the distribution of adaptive substitutions in the moving optimum model”. en. In: Genetics 183.4, pp. 1453–1476.

Krimbas, C B and S Tsakas (1971). “The genetics of Dacus oleae. V. Changes of esterase polymorphism in a natural population following insecticide control-selection or drift?” In: Evolution 25.3, p. 454.

Kruuk, Loeske E B (2004). “Estimating genetic parameters in natural populations using the ‘animal model’”. In: Philos. Trans. R. Soc. Lond. B Biol. Sci. 359.1446, pp. 873–890.

Kruuk, Loeske E B et al. (2000). “Heritability of fitness in a wild mammal population”. In: Proceedings of the National Academy of Sciences 97.2, pp. 698–703.

Lande, R (1979). “Quantitative genetic analysis of multivariate evolution, applied to brain: body size allometry”. In: Evolution 33.1, p. 402.

Leffler, Ellen M et al. (2012). “Revisiting an Old Riddle: What Determines Genetic Diversity Levels within Species?” In: PLoS Biol. 10.9, e1001388–9.

Lewontin, Richard C (1974). The genetic basis of evolutionary change. Vol. 560. Columbia University Press New York.

Lynch, Michael, Bruce Walsh, et al. (1998). Genetics and analysis of quantitative traits. Vol. 1. Sinauer Sunderland, MA.

Malaspinas, A S, O Malaspinas, and S N Evans (2012). “Estimating allele age and selection coefficient from time-serial data”. In: Genetics.

Mathieson, I and G McVean (2013). “Estimating selection coefficients in spatially structured populations from time series data of allele frequencies”. In: Genetics.

Mathieson, Iain et al. (2015). “Genome-wide patterns of selection in 230 ancient Eurasians”. In: Nature 528.7583, pp. 499–503.

Maynard Smith, John and John Haigh (1974). “The hitch-hiking effect of a favourable gene”. In: Genet. Res. 23.01, pp. 23–35.

McVicker, Graham, David Gordon, Colleen Davis, and Phil Green (2009). “Widespread Genomic Signatures of Natural Selection in Hominid Evolution”. In: PLoS Genet. 5.5, e1000471–16.

Messer, Philipp W, Stephen P Ellner, and Nelson G Hairston Jr (2016). “Can Population Genetics Adapt to Rapid Evolution?” In: Trends Genet. pp. 1–11.

Morley, F H W (1954). “Selection for economic characters in Australian Merino sheep”. In: Aust. J. Agric. Res. 5.2, pp. 305–316.

Mousseau, T A and D A Roff (1987). “Natural selection and the heritability of fitness components”. In: Heredity.

Mueller, L D, B A Wilcox, P R Ehrlich, and D G Heckel (1985a). “A direct assessment of the role of genetic drift in determining allele frequency variation in populations of Euphydryas editha”. In: Heredity.

Mueller, Laurence D, Lorraine G Barr, and Francisco J Ayala (1985b). “Natural selection vs. random drift: evidence from temporal variation in allele frequencies in nature”. In: Genetics 111.3, pp. 517–554.

Nachman, Michael W and Bret A Payseur (2012). “Recombination rate variation and speciation: theoretical predictions and empirical results from rabbits and mice”. en. In: Philos. Trans. R. Soc. Lond. B Biol. Sci. 367.1587, pp. 409–421.

Neher, R A and B I Shraiman (2011). “Genetic draft and quasi-neutrality in large facultatively sexual populations”. en. In: Genetics 188.4, pp. 975–996.

Neher, Richard A (2013). “Genetic Draft, Selective Interference, and Population Genetics of Rapid Adaptation”. In: Annu. Rev. Ecol. Evol. Syst. 44.1, pp. 195–215.

Nei, Masatoshi (1987). Molecular Evolutionary Genetics. en. Columbia University Press.

Nei, Masatoshi and Fumio Tajima (1981). “Genetic Drift And Estimation Of Effective Population Size”. In: Genetics 98.3, pp. 625–640.

Nordborg, Magnus et al. (2005). “The pattern of polymorphism in Arabidopsis thaliana”. en. In: PLoS Biol. 3.7, e196.

Ohta, T and M Kimura (1969). “Linkage disequilibrium at steady state determined by random genetic drift and recurrent mutation”. en. In: Genetics 63.1, pp. 229–238.

Orozco-terWengel, Pablo et al. (2012). “Adaptation of Drosophilato a novel laboratory environment reveals temporally heterogeneous trajectories of selected alleles”. In: Mol. Ecol. 21.20, pp. 4931–4941.

Pennings, Pleuni S and Joachim Hermisson (2006). “Soft Sweeps II—Molecular Population Genetics of Adaptation from Recurrent Mutation or Migration”. In: Mol. Biol. Evol. 23.5, pp. 1076–1084.

Pollak, Edward (1983). “A New Method For Estimating The Effective Population Size From Allele Frequency Changes”. In: Genetics 104.3, pp. 531–548.

Price, G R (1970). “Selection and covariance”. In: Nature.

Prout, T (1954). “Genetic Drift in Irradiated Experimental Populations of Drosophila Melanogaster”. en. In: Genetics 39.4, pp. 529–545.

Rajpurohit, Subhash et al. (2018). “Spatiotemporal dynamics and genome-wide association analysis of desiccation tolerance in Drosophila melanogaster”. In: Mol. Ecol.

Robertson, A (1970). “A Theory of Limits in Artificial Selection with Many Linked Loci”. In: Mathematical Topics in Population Genetics. Ed. by Ken-Ichi Kojima. Berlin, Heidelberg: Springer Berlin Heidelberg, pp. 246–288.

Robertson, Alan (1961). “Inbreeding in artificial selection programmes”. In: Genet. Res. 2.02, pp. 189–194.

Robertson, Alan (1966). “A mathematical model of the culling process in dairy cattle”. In: Anim. Sci. 8.1, pp. 95–108.

Robertson, Alan (1976). “Artificial selection with a large number of linked loci”. In: Proceedings of the International Conference on Quantitative Genetics. Ed. by Edward Pollak, Oscar Kempthorne, and Theodore B Bailey. Iowa State University Press, pp. 307–322.

Rockman, Matthew V, Sonja S Skrovanek, and Leonid Kruglyak (2010). “Selection at linked sites shapes heritable phenotypic variation in C. elegans”. en. In: Science 330.6002, pp. 372–376.

Santiago, Enrique and Armando Caballero (1995). “Effective size of populations under selection”. In: Genetics 139.2, pp. 1013–1030.

Santiago, Enrique and Armando Caballero (1998). “Effective Size and Polymorphism of Linked Neutral Loci in Populations Under Directional Selection”. In: Genetics 149.4, pp. 2105–2117.

Schmid, Karl J, Sebastian Ramos-Onsins, Henriette Ringys-Beckstein, Bernd Weisshaar, and Thomas Mitchell-Olds (2005). “A multilocus sequence survey in Arabidopsis thaliana reveals a genome-wide departure from a neutral model of DNA sequence polymorphism”. en. In: Genetics 169.3, pp. 1601–1615.

Sella, Guy, Dmitri A Petrov, Molly Przeworski, and Peter Andolfatto (2009). “Pervasive Natural Selection in the Drosophila Genome?” In: PLoS Genet. 5.6, e1000495.

Shaw, R G and F H Shaw (2013). “Quantitative genetic study of the adaptive process”. In: Heredity 112.1, pp. 13–20.

Stephan, W, The Wiehe, and M W Lenz (1992). “The effect of strongly selected substitutions on neutral polymorphism: analytical results based on diffusion theory”. In: Theor. Popul. Biol.

Stephan, Wolfgang, Yun S Song, and Charles H Langley (2006). “The hitchhiking effect on linkage disequilibrium between linked neutral loci”. en. In: Genetics 172.4, pp. 2647–2663.

Sved, J A (1971). Linkage Disequilibrium and Homozygosity of Chromosome Segments in Finite Populations. Vol. 2. Theoretical Population Biology. Elsevier.

Teotónio, Henrique, Ivo M Chelo, Martina Bradić, Michael R Rose, and Anthony D Long (2009). “Experimental evolution reveals natural selection on standing genetic variation”. In: Nat. Genet. 41.2, pp. 251–257.

Teplitsky, Céline, James A Mills, John W Yarrall, and Juha Merilä (2009). “Heritability of fitness components in a wild bird population”. en. In: Evolution 63.3, pp. 716–726.

Terhorst, Jonathan, Christian Schlötterer, and Yun S Song (2015). “Multi-locus Analysis of Genomic Time Series Data from Experimental Evolution”. In: PLoS Genet. 11.4, e1005069–29.

Thornton, Kevin R (2018). “Polygenic adaptation to an environmental shift: temporal dynamics of variation under Gaussian stabilizing selection and additive effects on a single trait”.

Turelli, M (1988). Population genetic models for polygenic variation and evolution. Proceedings of the Second International Conference On Quantitative Genetics.

Turelli, M and N H Barton (1990). “Dynamics of polygenic characters under selection”. In: Theor. Popul. Biol. 38.1, pp. 1–57.

Turelli, M and N H Barton (1994). “Genetic and statistical analyses of strong selection on polygenic traits: what, me normal?” In: Genetics 138.3, pp. 913–941.

Turner, T L and P M Miller (2012). “Investigating Natural Variation in Drosophila Courtship Song by the Evolve and Resequence Approach”. In: Genetics 191.2, pp. 633–642.

Turner, Thomas L, Andrew D Stewart, Andrew T Fields, William R Rice, and Aaron M Tarone (2011). “Population-Based Resequencing of Experimentally Evolved Populations Reveals the Genetic Basis of Body Size Variation in Drosophila melanogaster”. In: PLoS Genet. 7.3, e1001336–10.

Wallace, Bruce (1956). “Studies on irradiated populations ofDrosophila melanogaster”. en. In: J. Genet. 54.2, pp. 280–293.

Walsh, Bruce and Michael Lynch (2018). Evolution and Selection of Quantitative Traits. en. Oxford University Press.

Wang, Jinliang and Michael C Whitlock (2003). “Estimating effective population size and migration rates from genetic samples over space and time”. en. In: Genetics 163.1, pp. 429–446.

Waples, R S (1989). “A generalized approach for estimating effective population size from temporal changes in allele frequency”. In: Genetics 121.2, pp. 379–391.

Watterson, G A (1975). “On the number of segregating sites in genetical models without recombination”. In: Theor. Popul. Biol. 7.2, pp. 256–276.

Weir, Bruce S (1996). Genetic Data Analysis II: Methods for discrete population genetic data. Sinauer Associates.

Wiehe, T H and W Stephan (1993). “Analysis of a genetic hitchhiking model, and its application to DNA polymorphism data from Drosophila melanogaster”. en. In: Mol. Biol. Evol. 10.4, pp. 842–854.

Williamson, Robert J et al. (2014). “Evidence for widespread positive and negative selection in coding and conserved noncoding regions of Capsella grandiflora”. en. In: PLoS Genet. 10.9, e1004622.

Wray, Naomi R and Robin Thompson (1990). “Prediction of rates of inbreeding in selected populations”. In: Genet. Res. 55.01, pp. 41–54.

Wright, Sewall (1931). “Evolution in Mendelian populations”. In: Genetics 16.2, p. 97.

