## Supplementary material for "The Linked Selection Signature of Rapid Adaptation in Temporal Genomic Data": highlighted revisions from last bioRxiv version

$$\Delta p_t = \underbrace{\Delta_{N} p_t + \Delta_{M} p_t}_{\text{drift}} + \underbrace{\Delta_{H} p_t}_{\text{selection}} \tag{1}$$

*where, $\Delta_{N} p_t$, $\Delta_{M} p_t$, and $\Delta_{H} p_t$ are the neutral allele's frequency changes due to non-heritable variation in fitness between diploid individuals, Mendelian segregation of heterozygotes into offspring, and heritable variation in fitness (we refer to this as the* heritable change *in neutral allele frequency), respectively. Note while throughout the paper we consider the allele frequency change between adjacent generations, the same approach can be extended to situations where the study system cannot be observed every generation. Like the stochastic components of drift, the allele frequency change due to heritable fitness differences is directionless ($\mathbb{E}(\Delta_{H} p_t) = 0$). Additionally, since each component is uncorrelated with the others, the variance in allele frequency change is*

$$\mathrm{Var}(\Delta p_t) = \mathrm{Var}(\Delta_{N} p_t) + \mathrm{Var}(\Delta_{M} p_t) + \mathrm{Var}(\Delta_{H} p_t). \tag{2}$$

*The terms $\mathrm{Var}(\Delta_{N} p_t)$ and $\mathrm{Var}(\Delta_{M} p_t)$ capture the variance due to the random reproduction process, and the former can accommodate extra non-heritable variance in offspring number (as long as individuals are exchangeable with respect to their genotype; Cannings 1974), while the term $\mathrm{Var}(\Delta_{H} p_t)$ captures heritable fitness variation due to systematic differences in the fitness of individuals caused by their genotypes.*

*In addition to inflating the within-generation variance in allele frequency change, heritable fitness variation has another profound affect on neutral alleles: while the stochastic components of drift have independent effects on frequency change each generation, heritable variation in fitness creates* temporal autocovariance *in neutral allele frequency changes across generations. The contribution of temporal autocovariance is evident by writing the total cumulative allele frequency change as the sum of allele frequency changes each generation,*

$$\begin{aligned}
\mathrm{Var}(p_t - p_0) &= \mathrm{Var}(\Delta p_{t-1} + \Delta p_{t-2} + \ldots + \Delta p_0) \\
&= \sum_{i=0}^{t-1} \mathrm{Var}(\Delta p_i) + \sum_{i \neq j} \mathrm{Cov}(\Delta p_i, \Delta p_j) \\
\mathrm{Var}(p_t - p_0) &= \underbrace{\sum_{i=0}^{t-1} \left(\mathrm{Var}(\Delta_{N} p_i) + \mathrm{Var}(\Delta_{M} p_i)\right) +}_{\text{drift}} \\
&\quad \underbrace{\sum_{i=0}^{t-1} \mathrm{Var}(\Delta_{H} p_i)}_{\text{genetic variance in offspring number}} + \underbrace{\sum_{i \neq j} \mathrm{Cov}(\Delta_{H} p_i, \Delta_{H} p_j)}_{\text{temporal autocovariance}}.
\end{aligned} \tag{3}$$

*These covariance terms are expected to be non-zero only when there is heritable variation in fitness (assuming there is neither non-Mendelian segregation nor covariance between parental and offspring environment, e.g. as in Heyer et al. 2005).*

*Temporal autocovariance is caused by the persistence over generations of the statistical associations (linkage disequilibria) between a neutral allele and the fitnesses of the random genetic*

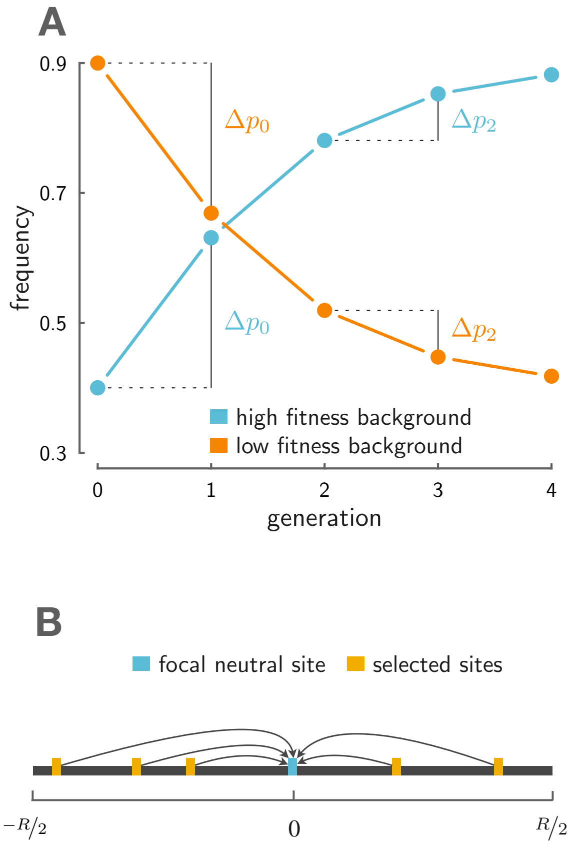


**Figure 1:** A: On an advantageous background (light blue), a neutral allele increases in frequency leading to a positive change in allele frequency early on, $\Delta p_0 = p_1 - p_0$. As long as some fraction of neutral alleles remain associated with this advantageous background, the neutral allele is expected to increase in frequency in later generations, here, $\Delta p_2 = p_3 - p_2$. This creates temporal autocovariance, $\text{Cov}(\Delta p_2, \Delta p_0) > 0$. Similarly, had the neutral allele found itself on a low-fitness background (orange), this would also create temporal autocovariance. B: This depicts the setup for our multilocus model. Multiple alleles (yellow) determine the fitness in a region $R$ Morgans in length, and these perturb the allele frequency trajectory of a focal neutral site (light blue).

#### 2 A model for multilocus temporal autocovariance

Here, we develop theory for the temporal autocovariance in a neutral allele's frequency changes through time, generated by the presence of heritable fitness in the population. We measure the temporal autocovariance $\text{Cov}(\Delta p_t, \Delta p_s)$ at a single diallelic neutral locus. Since only allele frequency changes due to heritable variation in *fitness* contribute to temporal autocovariance, we can focus exclusively on the behavior of $\Delta_H p_t$ in deriving our expressions for the autocovariance across timepoints. We imagine that an individual $i$ has fitness $f_i$, i.e. that their expected number of children is $f_i$. We assume a constant population size, and so the population average fitness $\mathbb{E}_i(f_i) = 1$. Additionally, we assume that all fitness variation has an additive, polygenic architecture. Then, with $L$ loci contributing to fitness, we can write individual $i$'s fitness as $f_i = 1 + \sum_{l=1}^{L} \alpha_{t,l} g_{i,l}$, where $\alpha_{t,l}$ is the effect size in generation $t$ and $g_{i,l} \in \{0, 1, 2\}$ is individual $i$'s gene content at locus $l$. *Here, each $\alpha_{t,l}$ is analogous to a selection coefficient acting at locus $l$, since the fitnesses for genotypes $A_1A_1$, $A_1A_2$, and $A_2A_2$ are 1, $1 + \alpha_{t,l}$, and $1 + 2\alpha_{t,l}$ respectively*. This formulation is approximately equivalent to exponential directional selection on some additively determined trait, *implying that selection does not create linkage disequilibria between unlinked loci*; see Appendix A.2 for more detail.

When fitness variation exists in the population (that is, $\text{Var}_i(f_i) > 0$), the frequency of the neutral allele changes stochastically, as more fit individuals leave more descendents that inherit the neutral allele they carry and less fit individuals leave fewer. Across the population, stochastic associations can form between the genetic components of an individual's fitness and the neutral allele they carry, leading the neutral allele frequency to change due to fitness differences across individuals. The total heritable change in neutral allele frequency $\Delta_H p_t$ can then be partitioned into each individual's contribution to this change based on their fitness $f_i$ and the number of the tracked neutral alleles they carry, $x_i \in \{0, 1, 2\}$, giving us

$$\Delta_H p = \frac{1}{2N} \sum_{i=1}^{N} x_i (f_i - 1) \tag{4}$$

(Santiago and Caballero 1995). Substituting each fitness $f_i$ with its genetic basis and simplifying (see Appendix Section A.2 for details) gives,

$$\Delta_{_H} p_t = \sum_{l=1}^{L} \alpha_{t,l} D'_{t,l} + \sum_{l=1}^{L} \alpha_{t,l} D''_{t,l} \tag{5}$$

where $D'_{t,l}$ is the *gametic* linkage disequilibrium between the neutral allele and the allele at the selected site $l$ on the same gamete, whereas $D''_{t,l}$ is the *non-gametic* linkage disequilibria, or the covariance *across* the neutral and selected allele on the two different gametes forming an individual (see Weir 1996, p. 121 for details and Appendix Figure 8A for an illustration). Intuitively, this expression tells us that the heritable change in neutral allele frequency is determined by the gametic and non-gametic linkage disequilibrium between the neutral site and all sites that affect an individual's fitness, scaled by the magnitude of each selected locus's effect. Alternatively, we can see this as the multivariate breeder's equation, where the neutral allele is a correlated trait responding to selection on other traits/loci (Lande 1979). This expression is the multilocus analog of the change in a neutral site's frequency due to hitchhiking at a single linked site (see e.g. equations (2) and (3) in Stephan et al. 2006).

Because the effects of the non-gametic LD are relatively weak compared to the gametic LD for tightly linked loci (see Appendix Section A.5 for an expression of their strength), we ignore these, and hereafter omit the primes in our notation so that $D_{t,l}$ refers to $D'_{t,l}$. Since $\mathbb{E}(\Delta_{_H} p_t) = 0$, we can write the covariance $\mathrm{Cov}(\Delta_{_H} p_t, \Delta_{_H} p_s)$ as $\mathbb{E}(\Delta_{_H} p_t \Delta_{_H} p_s)$. Hereafter, we also omit the subscript $H$ since $\mathrm{Cov}(\Delta p_t, \Delta p_s) = \mathrm{Cov}(\Delta_{_H} p_t, \Delta_{_H} p_s)$. Expanding these terms, the covariance between the allele frequency changes at generations $t$ and $s$ can be written as

$$\begin{aligned}\mathrm{Cov}(\Delta p_t, \Delta p_s) &= \mathbb{E}\left[\left(\sum_{l=1}^{L} \alpha_{t,l} D_{t,l}\right)\left(\sum_{l=1}^{L} \alpha_{t,l} D_{s,l}\right)\right] \\ &= \underbrace{\sum_{l=1}^{L} \alpha_{t,l}\alpha_{s,l} \mathbb{E}(D_{t,l} D_{s,l})}_{\text{persistence of associations selected site } l} + \underbrace{\sum_{l \neq k} \alpha_{t,k}\alpha_{s,l} \mathbb{E}(D_{t,k} D_{s,l})}_{\text{cross-associations between two selected sites}} \quad . \end{aligned} \tag{6}$$

$$\begin{aligned}
\mathrm{Cov}(\Delta p_t, \Delta p_s) &= \sum_{l=1}^{L} \alpha_l^2 \mathbb{E}(D_{t,l}D_{s,l}) \\
&= \sum_{l=1}^{L} \alpha_l^2 \mathbb{E}(\mathcal{R}_{t,l}^2) p_t(1-p_t) p_{s,l}(1-p_{s,l})(1-r_l)^{s-t} && \text{(7a)} \\
\frac{\mathrm{Cov}(\Delta p_t, \Delta p_s)}{p_t(1-p_t)} &= \sum_{l=1}^{L} \alpha_l^2 p_{s,l}(1-p_{s,l}) \mathbb{E}(\mathcal{R}_{t,l}^2)(1-r_l)^{s-t}, && \text{(7b)}
\end{aligned}$$

where $\mathbb{E}(\mathcal{R}_{t,l}^2)$ is the square of the correlation between the neutral site and selected site $l$ at time $t$ (a common measure of LD; Hill and Robertson 1968), and $r_l$ is the recombination fraction between the neutral site and selected site $l$.

We can further simplify this expression by assuming that there is no covariation between the additive genic variation at a selected site and the linkage disequilibrium between that selected site and the neutral site (see Appendix Section A.2 for more detail). This allows us to factor out the average additive genic variation for fitness at time $s$ and write the covariance as

$$\frac{\mathrm{Cov}(\Delta p_t, \Delta p_s)}{p_t(1-p_t)} = \frac{V_a(s)}{2L} \sum_{l=1}^{L} \mathbb{E}(\mathcal{R}_{t,l}^2)(1-r_l)^{s-t} \tag{8}$$

where $V_a(s)$ is the additive genic variance for fitness, which is the additive genetic variance for fitness ($V_A$) without the contribution of linkage disequilibria between selected sites, $V_a(s) = 2\sum_l \alpha_l^2 p_l(s)(1-p_l(s))$. *In part, our expression not relying on the linkage disequilibria between selected sites is a result of ignoring the second term in Equation (6); we revisit the consequences of this assumption further on in Section 2.5.*

$$\Sigma_{t,s} := \frac{\mathbb{E}_n(\mathrm{Cov}(\Delta p_t, \Delta p_s))}{\mathbb{E}_n(p_t(1-p_t))} = \frac{V_a(s)}{2} \underbrace{\int_0^R \mathbb{E}(\mathcal{R}_t^2(r(c)))\,(1-r(c))^{(s-t)} \frac{2(R-c)}{R^2}\,dc}_{\mathcal{A}(R,t,s)} \tag{10}$$

where $\mathbb{E}_n(\cdot)$ indicates an expectation taken over the position of the randomly placed neutral sites, and we define $\mathcal{A}(R,t,s)$ as the average linkage disequilibrium between selected and neutral sites that persists from generations $t$ to $s$ ($t \leq s$). As is common with estimating the expected values of other ratios like $F_{ST}$ (Bhatia et al. 2013), we use a ratio of expectations rather than the expectation of the ratio.

We can also use this expression to calculate the variance of allele frequency change. The standardized variance ${}^{\mathrm{Var}(\Delta p_t)}/_{p_t(1-p_t)}$ has two components: the drift term and the heritable variance in offspring number. Adding these independent contributions, the standardized variance is

$$\frac{\mathbb{E}_n(\mathrm{Var}(\Delta p_t))}{\mathbb{E}_n(p_t(1-p_t))} = \frac{V_N+2}{8N} + \frac{V_a}{2}\mathcal{A}(R,t,s) \tag{11}$$

where $V_N$ is the non-heritable variance in offspring number. Under a Wright–Fisher model of reproduction, $V_N \approx 2$, this simplifies to

$$\Sigma_{t,t} := \frac{\mathbb{E}_n(\mathrm{Var}(\Delta p_t))}{\mathbb{E}_n(p_t(1-p_t))} = \frac{1}{2N} + \frac{V_a}{2}\mathcal{A}(R,t,s). \tag{12}$$

When combined, this expression for the variance in allele frequency change and our expression for temporal autocovariance are in agreement with Robertson (1961) and Santiago and Caballero (1995; 1998) when predicting the total variance in allele frequency change; see Appendix A.6. *With the above expressions for the variances and covariances, we have a complete set of theoretic expressions for the variance-covariance matrix of allele frequency change, which we call* $\boldsymbol{\Sigma}$*, with the diagonal variance elements* $\Sigma_{t,t}$ *given by Equation (12), and the upper- and lower-triangle covariance elements* $\Sigma_{t,s}$ $(t \neq s)$ *given by Equation (10).*

We record 50 generations of *simulated evolution*, after which we compute the standardized *sample* temporal variance-covariance matrix $\mathbf{Q}$ *(this is the sample analog of our theoretic variance-covariance matrix* $\boldsymbol{\Sigma}$*)*, for each replicate *as follows*. First, we mark frequencies reaching fixation or loss as missing values. *This allows the frequency changes before fixation/loss to contribute to the measured covariance, rather than removing the entire locus's trajectory, which would act to condition the covariance on more intermediate frequencies. Note that one cannot ignore fixations or losses, as these have* $\Delta p_t = 0$ *and thus an autocovariance of zero, which would act to underestimate the true level of autocovariance at segregating sites. Having marked fixations/losses as missing, we take the frequency matrix* and calculate a vector of allele frequency changes $\Delta \vec{p}_n = [\Delta p_{n,1}, \Delta p_{n,2}, \ldots, \Delta p_{n,\tau}]$ using each neutral locus $n$'s $\tau + 1$ observed generations. *Finally*, we calculate the $\tau \times \tau$ sample standardized variance-covariance matrix $\mathbf{Q}$, averaging over $M$ neutral loci such that element $Q_{t,s}$ is calculated as

$$Q_{t,s} = \frac{\frac{1}{M-1} \sum_{n=1}^{M} \left( \Delta p_{n,t} \Delta p_{n,s} - \left( \frac{1}{M} \sum_{n=1}^{M} \Delta p_{n,t} \right) \left( \frac{1}{M} \sum_{n=1}^{M} \Delta p_{n,s} \right) \right)}{\frac{1}{M} \sum_{n=1}^{M} p_{\min(t,s)} \left( 1 - p_{\min(t,s)} \right)}, \tag{16}$$

*though see Appendix Section A.8 for a bias-corrected version when sample, rather than population allele frequencies are used.* Sums over missing values only use pairwise-complete observations, implemented by R's `cov()` function's `use='pairwise.complete'` argument.

We have extensively validated our simulation procedure in a neutrally evolving population, ensuring that the decay of linkage disequilibrium and the allele frequency change match expectations (see Supplementary Figures 3, 4, and 5).

#### 2.5 Comparing theory to simulation results

To validate our expressions for temporal autocovariance, we compare the levels of autocovariance and variance predicted by Equation (10) and (12) to the average levels observed across simulation replicates. To calculate the theoretic values of temporal autocovariance and variance, our expression requires the additive genic variation at $s$, $V_a(s)$; however, as described in Section 2.3, we lack an analytic expression for the dynamics of genic variation to plug into $V_a(s)$. Following the approach of others in evolutionary quantitative genetics (Turelli and Barton 1994, p. 930), we substitute the numerical values calculated directly from the simulation data for $V_a(s)$. Additionally we consider two other numerical values related to the additive genic variance: the observed additive genetic variance from our simulations ($V_A(s) = \text{Var}_i(z_i)$ at time $s$, which includes the contribution of LD between selected sites), and the additive genic variation at time $s$ as approximated by the observed decay in the sum of site heterozygosity at neutral sites ($V_{a,ssh_n}(1)$, as described in Section 2.3).

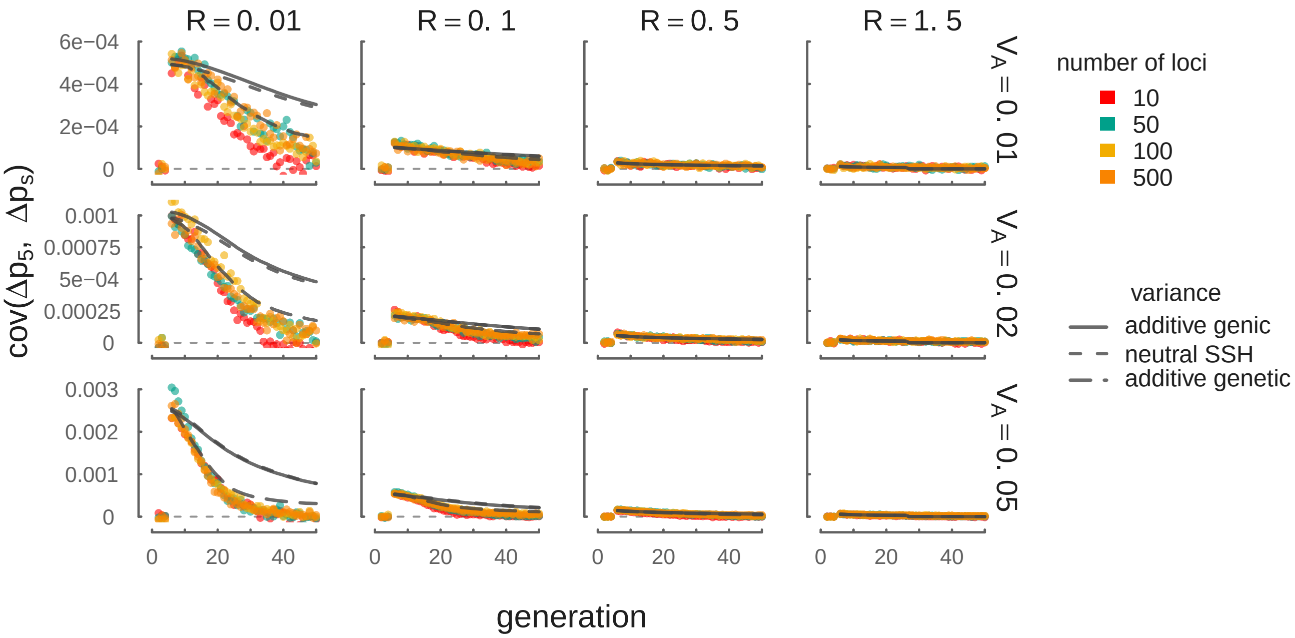


**Figure 2:** In each panel the temporal autocovariance $\text{Cov}(\Delta p_5, \Delta p_s)$ is shown on the y-axis while generation $s$ varies along the x-axis. Selection is initiated on the $5^{\text{th}}$ generation, so $\Delta p_5$ is the neutral allele's frequency change across the first generation of selection. Each point is the temporal autocovariance between $\Delta p_5$ and the $\Delta p_s$ in a region, averaged over 100 simulation replicates, with the colors indicating the number of selected loci. The gray curves indicate the theoretic predictions (for $L = 500$ loci only) using Equation (10), with the equation's variance provided by the empirically observed additive genic (solid), the additive genetic (long dashes), and the neutral sum of site heterozygosity approximation (short dashes). A thin horizontal dashed line indicates $y = 0$. Across the columns, the level of recombination (in Morgans) is varied; across rows, the initial level of additive genetic variation is varied. Note that while our results here are between the frequency change at the onset of selection $\Delta p_5$ and some later change $\Delta p_s$, our covariance theory matches simulation results between any two arbitrary frequency changes $\Delta p_t$, and $\Delta p_s$; see Supplementary Figure 7.

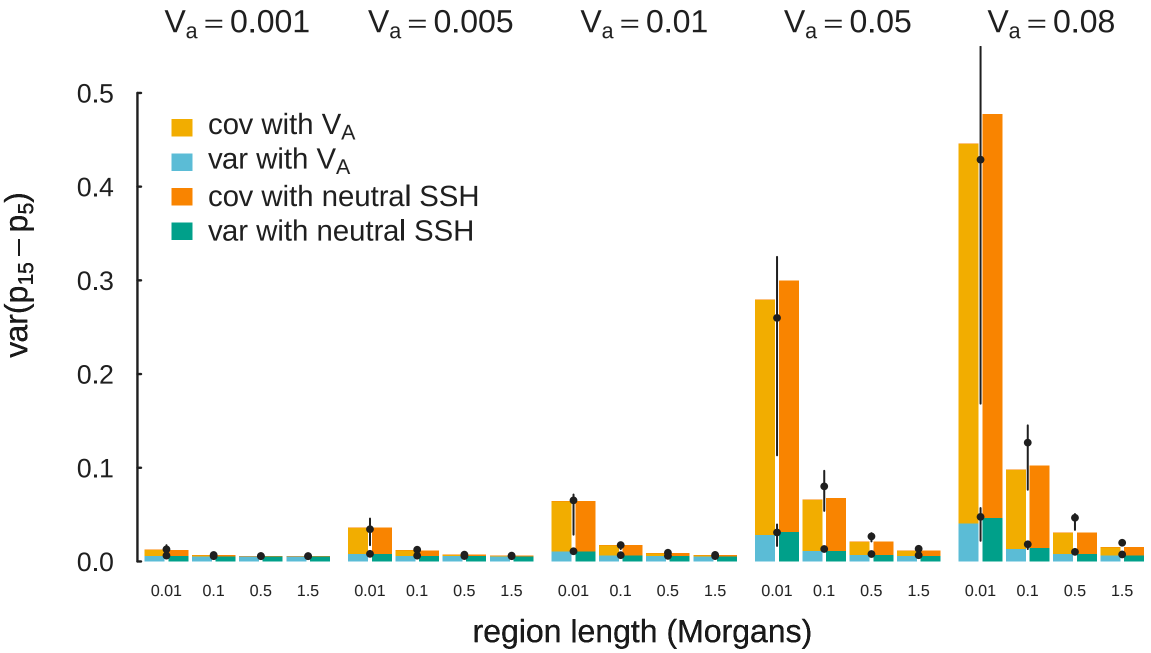


**Figure 3:** Summing over generations, Equations (6) and (12) accurately predicts the total variation in allele frequency change due to variance and covariance components. The predicted cumulative variance in allele frequency change across the ten generations after selection ($\mathrm{Var}(p_{15} - p_5)$) is shown as bars, using both the empirical additive genetic variation $V_A$ (bars to the left of the pointrange), and the empirical neutral sum of site heterozygosity (bars to the right of the pointrange). The variance and covariance components are represented by blue/green and orange/yellow tones respectively. Finally, we show the averaged results of our simulations as pointranges, with the point depicting the average and the bars representing the lower and upper quartiles.

*covariances matrices non-positive definite, which makes maximum likelihood estimation perhaps much more difficult. Throughout, we use population allele frequencies (i.e. there is no sampling noise) which simplifies the description of the method; in Appendix Section A.8 we describe how the method is changed by finite sampling of chromosomes from a population.*

From our multilocus theory, we have analytic expressions for each element of the $\tau \times \tau$ covariance matrix of allele frequency changes in a region. To model the additive genic variance through time, we use the empirical neutral sum of site heterozygosity approximation as described in Section 2.3. This approximates the rate that additive genic variation decreases through time from some initial level, $V_A(1)$, which we wish to estimate. In total, we have $\tau + {}^{\tau(\tau-1)}/_2$ unique moment equations, which for the variance and covariance are defined as

$$\frac{\mathrm{Var}(\Delta p_t)}{\mathbb{E}(p_t(1-p_t))} = \frac{\widehat{V_A(1)}}{2} \frac{SSH_n(t)}{SSH_n(1)} \mathcal{A}(R,t,t) + \widehat{F} \quad := \quad \Sigma_{t,t} \tag{18}$$

$$\frac{\mathrm{Cov}(\Delta p_t, \Delta p_s)}{\mathbb{E}(p_t(1-p_t))} = \frac{\widehat{V_A(1)}}{2} \frac{SSH_n(s)}{SSH_n(1)} \mathcal{A}(R,t,s) \quad := \quad \Sigma_{t,s} \ \ (\text{for } s > t). \tag{19}$$

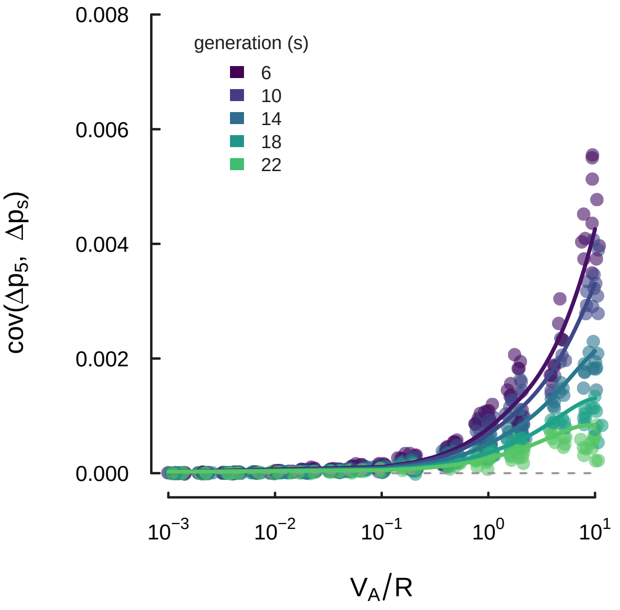


**Figure 4:** The compound parameter ${}^{V_A}\!/_{R}$ and the number of generations between the temporal autocovariance $s - t$ largely determines the magnitude of the temporal autocovariance across a wide spectrum of $V_A$ and $R$ parameters. Each point is a simulation replicate with its x-axis position given by ${}^{V_A}\!/_{R}$, the y-axis position equal to the temporal autocovariance, and the number of elapsed generations $(s - t)$. Each line is a loess curve fit through each set of points for a particular generation (with smoothing parameter $\alpha = 0.9$).

Here the first line gives the form of $\tau$ equations for the variance of allele frequency changes between subsequent generations, which includes the effect of genetic drift, $\widehat{F} = {}^{1}\!/_{2N}$. The second line gives the form of the covariances of allele frequency changes among different generations. The term $\mathcal{A}(R, t, s)$ is the average level of linkage disequilibrium after the $s - t$ generations that have elapsed, given there are $R$ Morgans of recombination. In our multilocus theory section and simulations, this is equal to the integral in Equation (10). However, we can also directly calculate a sample $\overline{\mathcal{A}(R, t, s)}$ from observed linkage disequilibrium in a region (for details, see Appendix Equation (53)).

Following the method-of-moments, we equate each of these independent $\tau + {}^{\tau(\tau-1)}\!/_{2}$ equations for $\Sigma_{t,s}$ to the observed sampling moments, the elements $Q_{t,s}$ of the upper triangle of the observed heterozygosity-normalized covariance matrix $\widehat{\mathbf{Q}}$ described in Equation (16). This yields $\tau + {}^{\tau(\tau-1)}\!/_{2}$ equations with 2 unknown parameters: $\widehat{V_A(1)}$ and $\widehat{N}$. We solve this overdetermined system of equations using least squares, an approach similar to the generalized method-of-moments in econometrics (Hansen 1982). This approach finds parameter estimates that minimize the squared error between the moment-based parameter estimate and the true parameter value, with respect to the true parameter value. We write the elements $Q_{t,s}$ of the upper triangle of the observed covariance matrix in the vector $\vec{q}$, and write the method-of-moments equations as,

$$\vec{q} = \widehat{V_A(1)}\vec{a} + \widehat{F}\vec{b} + \vec{\varepsilon} \tag{20}$$

where the elements of $\vec{a}$ and $\vec{b}$, in the same order as $\vec{q}$, are given by

$$a_{t,s} = \frac{1}{2}\frac{SSH_n(s)}{SSH_n(t)}\mathcal{A}(R,t,s), \;\; b_{t,s} = \delta_{t,s} \tag{21}$$

where $\delta_{t,s}$ is an indicator variable that is 1 when $s = t$ and zero otherwise.

Then, we can readily estimate the parameters $\widehat{V_A(1)}$ and $\widehat{F}$ using least squares. We then obtain an estimate of $\widehat{N}$ by taking $1/2\widehat{F}$. Since these equations are not statistically independent, we cannot assume $\mathrm{Cov}(\vec{\varepsilon}) = \sigma^2\mathbf{I}$. However, this does not affect our estimates $\widehat{V_A(1)}$ and $\widehat{F}$, as the least squares procedure is unbiased regardless of the covariance structure between the error terms (Christensen 2011, p. 26).

$$\Sigma_{t,s,i} = \frac{\widehat{V_A(1)}}{2}\frac{1}{B}\frac{SSH_n(s)}{SSH_n(1)}\mathcal{A}(R_i,t,s) + \widehat{F}\delta_{t,s} + \varepsilon, \;\; \text{for } s \geq t. \tag{22}$$

However, we expect *a priori* that windows containing more coding bases might disproportionately contribute to the total additive genetic variance. This suggests an alternative model to fit where partitions of the additive genetic variance across windows are proportional to the number of coding bases, similar to background selection and other linked selection models (Corbett-Detig et al. 2015; McVicker et al. 2009; Rockman et al. 2010). Thus, we could write total

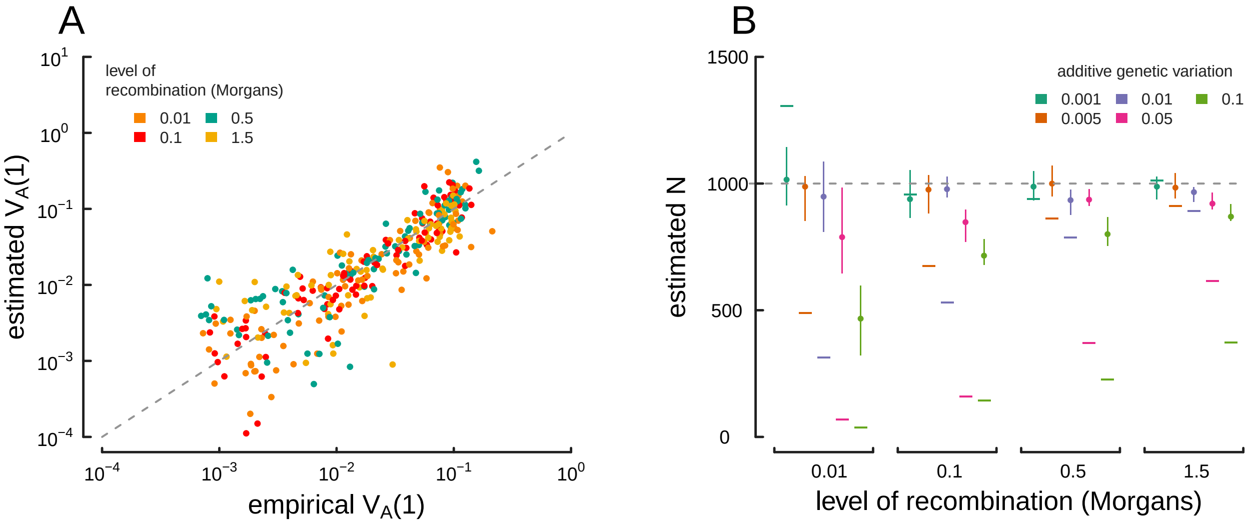


**Figure 5:** True parameter values and estimates using the method-of-moments approach on multilocus simulation data. (A) The true $V_A(1)$ (x-axis) and $\widehat{V_A(1)}$ estimated from the variance/covariance matrix (y-axis) for each simulation replicate across different levels of recombination (indicated by each point's color). The dashed gray line shows the $y = x$ line where an estimate is exactly true to its real value. Note that the plot is on a log-log scale, as $V_A$ varies across orders of magnitude in our simulations. (B) *Estimated drift-effective population size ($\widehat{N}$) across a range of simulations with different levels of additive genetic variance and recombination. Each point denotes the median, with lines denoting the interquartile range. A simple temporal estimate of the effective population size, estimated with accounting for the effects of selection, is averaged for each replicate and plotted as a dash.* The true value ($N = 1,000$) is shown with the dashed gray line. *Population frequencies (without sampling noise) are used in this figure; see Appendix Figure 1 for an analogous figure calculated with sample frequencies.*

$V_A(1) = v_A(1)\sum_{i=1}^{B} w_{\text{CBP},i}$ where $w_{\text{CBP},i}$ is the number of coding or exonic basepairs in window $i$ (this could be any quantifiable annotation feature in the window), and $W_{\text{CBP}} = \sum_{i=1}^{B} w_{\text{BP}}, i$ is the total number of coding bases in the genome. With window $i$ contributing $v_A(1)w_{\text{CBP},i}$ to the additive genetic variance and having map length $R_i$, we now define $\vec{q}$, $\vec{a}$, and $\vec{b}$ as having elements given by the equations

$$\Sigma_{t,s,i} = \frac{\widehat{V_A(1)}}{2}\frac{w_{\text{CBP},i}\; SSH_n(s)}{W_{\text{CBP}}\; SSH_n(1)}\mathcal{A}(R_i, t, s) + \widehat{F}\delta_{t,s} + \varepsilon, \quad \text{for } s \geq t. \tag{23}$$

Again, the parameters of this model, $\widehat{V_A(1)}$ and $\widehat{N}$, can be estimated with least squares. When analyzing genome-wide data, these various models could potentially be compared to an out-of-sample procedure, using inferred parameters to estimate the mean-squared predictive error between the two models for the remaining windows (Elyashiv et al. 2016). The confidence intervals for our method-of-moments estimates could be obtained through bootstrapping genomic windows since the errors are not identically and independently distributed.

#### 3.1 Estimating the proportion of allele frequency change due to linked selection

We can also estimate what fraction of allele frequency change over $t$ generations $(\text{Var}(p_t - p_0)/t)$ is due to linked selection acting to perturb the frequency trajectories of neutral alleles. We have developed two approaches: first, a more conservative approach that considers only the contribution of selection to the temporal autocovariance, and second, a more exact approach that uses the estimated effective population size to include the contribution of selection to both variances and covariances of allele frequency change.

First, a simple estimate of the total fraction of the variance in allele frequency change ($G$) caused by linked selection is

$$G = \frac{\sum_{t \neq s} \text{Cov}(\Delta p_t, \Delta p_s)}{\text{Var}(p_t - p_0)}. \tag{24}$$

However, this estimator is conservative because it ignores the contribution that linked selection has on the variance in allele frequency change across a single generation (the $\text{Var}(\Delta_{{}_H} p_t)$ term in Equation 2). If we include these variance terms, we have a less-conservative estimator we call $G'$,

$$\begin{aligned} G' &= \frac{\sum_{t \neq s} \text{Cov}(\Delta p_t, \Delta p_s) + \sum_{i=1}^{t} \text{Var}(\Delta_{{}_H} p_i)}{\text{Var}(p_t - p_0)} \\ &= 1 - \frac{\sum_{i=1}^{t} \text{Var}(\Delta_{{}_M} p_i) + \sum_{i=1}^{t} \text{Var}(\Delta_{{}_N} p_i)}{\text{Var}(p_t - p_0)}. \end{aligned} \tag{25}$$

We can think of the numerator of the second term in Equation (25) as the variance in allele frequency change in a Wright–Fisher population without selection. Recall that under a Wright–Fisher model, the standardized variance across $t$ generations is approximately $(1 - \exp(-t/2N)) \approx p_0(1 - p_0) \times t/2N$, where this second approximation works for short time spans ($t/2N \ll 1$). This suggests that we can use our method-of-moments estimate of the effective population size without

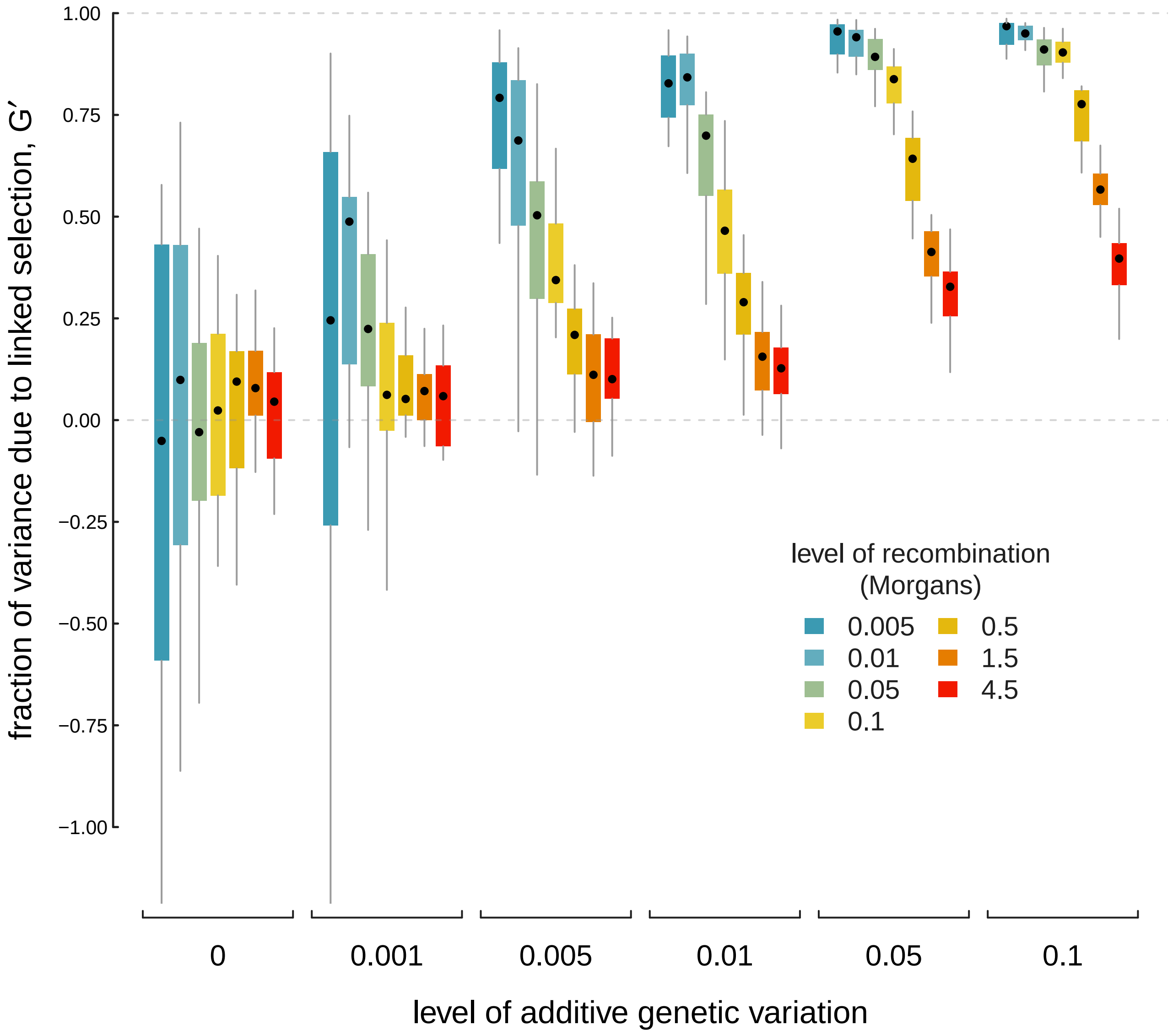


**Figure 6:** The proportion of total variance in allele frequency changes caused by linked selection, $G'$, across a variety of different levels of additive genetic variance (each group of boxplots), and different levels of recombination (each colored boxplot within a group). Each boxplot shows the spread of values across 20 replicates, with $\widehat{N}$ being calculated across each replicate.

the effects of selection, $\widehat{N}$, and compare the fraction of standardized variance we expect under this rate of drift to the empirical standardized variance,

$$G' = 1 - \frac{t\mathbb{E}(p_0(1-p_0))}{2\widehat{N}\,\mathrm{Var}(p_t - p_0)}. \tag{26}$$

Figure 6 shows the estimated $G'$ from the method-of-moment $\widehat{N}$ estimates using 20 replicates of simulated data. We learn three important points about our $G'$ estimator. First, for low $V_A$, or $V_A = 0$, the estimator is quite noisy. Second, although the signal can be noisy for low $V_A$, the relationship between $G'$ and level of recombination is consistent with selection affecting the total variance in allele frequency changes across the genome. Finally, this suggests that a negative relationship between $G'$ and recombination rate, calculated in windows across the genome, is a robust signal of linked selection impacting the total variance in allele frequency change. Furthermore, we would expect a positive relationship between the number of coding basepairs per window (when such information is available) and $G'$, which could serve as another robust signal of linked selection impacting the total variance in allele frequency change.

#### 3.2 Fluctuating Selection

Thus far we have assumed that fitness effect sizes are constant through time, that is $\alpha_{t,l} = \alpha_{s,l}$, for all $t, s$. In natural populations, changes in the environment or composition of the population may cause these effect sizes to change through time, due to changing selection pressures and changes in the epistatic environment experienced by alleles. If these changes occur within the timeframe of recorded allele frequency changes, the levels of temporal autocovariance will differ from the levels predicted from our directional selection theory. However, from Equation (6) we can see that the magnitude of temporal autocovariance is determined in large part by $\mathbb{E}(\alpha_t \alpha_s)$.

Here, we discuss how temporal autocovariance behaves under an example of strong fluctuating selection: when selection on a trait changes direction at some point. Specifically, we change the fitness function $w(z_i) = e^{z_i}$ to $w(z_i) = e^{-z_i}$ after some timepoint $t^*$; this is equivalent to changing $\alpha_{s,l} = -\alpha_{t,l}$ iff $s \geq t^*$, and $\alpha_{s,l} = \alpha_{t,l}$ otherwise for all other $s < t^*$.

When such a strong change in the direction of selection occurs, the temporal autocovariance between timepoints before and after the change becomes negative, since temporal autocovariance is determined by the product $\alpha_t \alpha_s$ for $t \neq s$ (here we are holding effects constant across loci). We have validated this using the same simulation procedure as described in Section 2.4, except on generation 15 we reverse the direction of selection on the trait by changing the fitness function $w(z) = e^z$ to $w(z) = e^{-z}$. In Figure 7A, we show the temporal autocovariance $\mathrm{Cov}(\Delta p_5, \Delta p_s)$ for varying $s$ along the x-axis (in this case, $V_a = 0.05$, $R = 0.1$, and $L = 500$). During the first five generations, the temporal autocovariance behaves as it does under directional selection, decaying due to the decrease in additive genetic variance and the breakdown of linkage disequilibrium. Then, on the 15$^{\text{th}}$ generation, the direction of selection on the trait with breeding value $z_i$ reverses and temporal autocovariance becomes negative since $\alpha_{s,l}\alpha_{t,l} < 0$ for all $s \geq t^*$ and $t < t^*$. Under this simple flip in the direction of selection pressure, the genic variance, $V_a(s) = 2\alpha^2 \sum_l p_{s,l}(1-p_{s,l})$, in the expressions for the temporal autocovariance can be replaced with $V_a(s) = 2\alpha_s \alpha_t \sum_l p_{s,l}(1-p_{s,l})$, akin to a genetic/genic covariance. In Figure 7A, the gray line is our predicted level of temporal autocovariance proportional to $V_A \mathcal{A}(R,t,s)$ given by Equation (8) before generation 15, and after generation it is proportional to $-V_A \mathcal{A}(R,t,s)$ (using the empirical additive genetic variance).

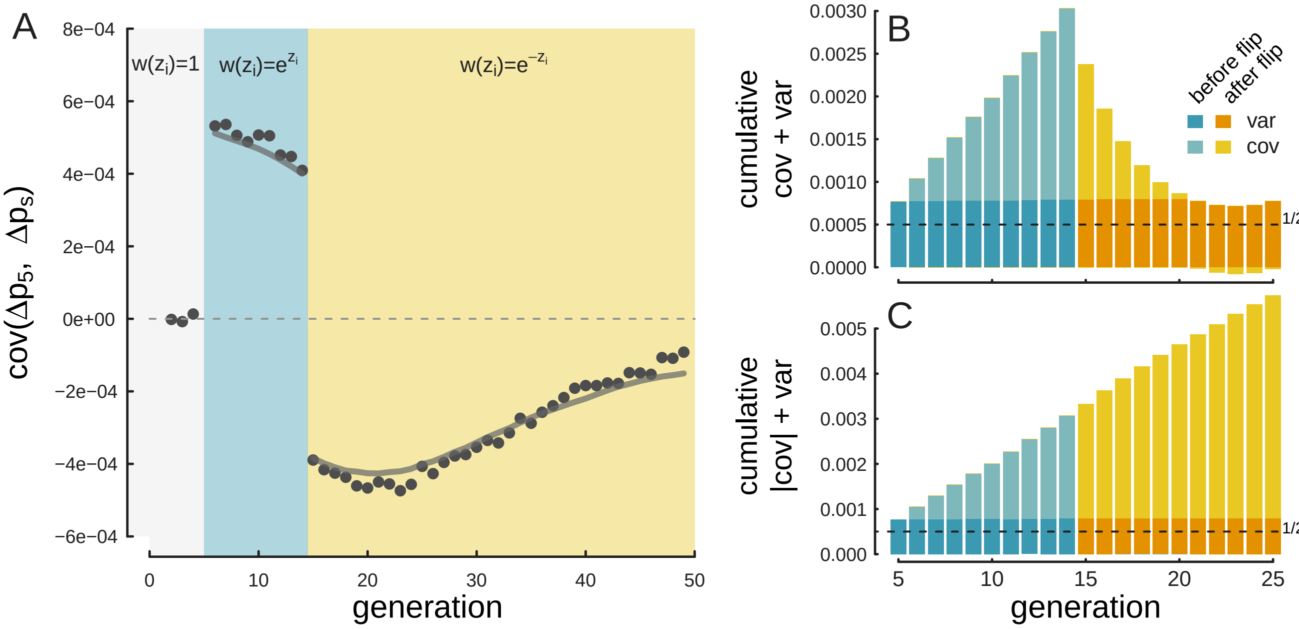


**Figure 7:** A: The covariance $\text{Cov}(\Delta p_5, \Delta p_s)$, where $s$ is varied on the x-axis, for $V_a = 0.05$, $R = 0.1$, and $L = 500$ averaged over 100 replicates. Selection begins in generation 5 (with the fitness function $w(z_i) = e^{z_i}$) and on generation 15 the direction of selection flips (and the fitness function becomes $w(z_i) = e^{-z_i}$). The gray line shows our directional selection temporal autocovariance prediction modified so that after the flip in the direction the trait is selected, we plot the *negative* theoretic level of temporal autocovariance. B: The average cumulative variance and covariances through the generations for the same simulation parameters, with the height of the bar representing the total cumulative variation $\text{Var}(p_t - p_0)/t$. Since the direction of selection flips, the covariances terms after generation 15 become negative, leading the total variance to decrease (the negative covariances are plotted below the x-axis line). After generation 21, the total covariance is negative, leading the total variance to dip below the level of variance alone (determined by drift and heritable fitness variation). The dark gray dashed line shows the level of variance expected by drift alone ($^{\text{Var}(p_t - p_0)}/_{t} = {}^{1}/_{2N}$). C: The effect of using the absolute value of covariance, which prevents the negative autocovariances from canceling out the effects of other covariances before the direction of selection changed.

To more fully capture the contribution of selection allele frequency change, we modify $G$, using the absolute value of the covariances,

$$G_{abs} = \frac{\sum_{t \neq s} |\,\mathrm{Cov}(\Delta p_t, \Delta p_s)|}{\mathrm{Var}(p_t - p_0)} \tag{27}$$

Wray, Naomi R and Robin Thompson (1990). "Prediction of rates of inbreeding in selected populations". In: *Genet. Res.* 55.01, pp. 41–54.

Wright, Sewall (1931). "Evolution in Mendelian populations". In: *Genetics* 16.2, p. 97.

Wright, Sewall (1938). “Size of population and breeding structure in relation to evolution”. In: *Science* 87.2263, pp. 430–431.

### A Appendix

**Table 1:** Notation

| Symbol | Usage |
|---:|---|
| $p_t$ | Allele frequency in generation/timepoint $t$ |
| $\Delta p_t$ | Allele frequency change between generations $t+1$ and $t$, $\Delta p_t = p_{t+1} - p_t$ |
| $\Delta_{N} p_t$ | Frequency change due to non-heritable variation in fitness, (1), (2) |
| $\Delta_{M} p_t$ | Frequency change due to Mendelian segregation, (1), (2) |
| $\Delta_{H} p_t$ | Frequency change due to heritable differences, (1), (2) |
| $N$ | Census population size of breeding individuals |
| $N_e$ | Effect population size |
| $f_i$ | Fitness (expected number of offspring) of individual $i$, (28) |
| $\alpha_{t,l}$ | Effect size in generation $t$ and locus $l$, (5), (36) |
| $L$ | Total number of loci impacting fitness, (5) |
| $g_{i,l} \in \{0, 1, 2\}$ | Individual $i$'s gene count at locus $l$, (37) |
| $x_i \in \{0, 1, 2\}$ | Individual $i$'s neutral gene count at the tracked neutral site, (5), (37), (38) |
| $D_{t,l}$ or $D'_{t,l}$ | Gametic linkage disequilibrium between the tracked neutral site and selected locus $l$ at time $t$, Supplementary Figure 8, (5), (37), (38) |
| $D''_{t,l}$ | Non-gametic disequilibrium between the tracked neutral site and selected locus $l$ at time $t$, Supplementary Figure 8, (5), (37), (38) |
| $\mathbb{E}(\mathcal{R}^2_{t,l})$ | The squared correlation coefficient of linkage disequilibrium between the tracked neutral site and selected site $l$ at time $t$, (7), (44) |
| $r_l$ | The recombination fraction between the tracked neutral site and selected site $l$ |
| $V_a(s)$ | The additive genic variance, (8) |
| $V_A(s)$ | The additive genetic variance, (17) |
| $R$ | The total level of recombination in the region, in Morgans, (9) and Figure 1 |
| $r(g)$ | A mapping function (i.e. Haldane's), which maps a position $g$ to a recombination fraction. |
| $\rho$ | The population recombination rate, $\rho = 4Nr$, Section 2.2 |
| $\mathcal{A}(R, t, s)$ | The average linkage disequilibrium in a region of $R$ Morgans, that persisted from generation $t$ to generation $s$, (10) |
| $V_N$ | The non-heritable variance in offspring number, (11) |
| $SSH(t)$ | The sum of site heterozygosity at selected sites time $t$, (13) |
| $SSH_n(t)$ | The sum of site heterozygosity at neutral sites at time $t$, (13) |
| $ssh_n(t)$ | The proportion of sum of site heterozygosity at neutral sites at time $t$ relative to $SSH(1)$, $SSH_n(t)/SSH_n(1)$ (15) |
| $z_i$ | The breeding value of the trait that determines fitness, $z_i = \sum_{i=1}^{L} \alpha g_{i,l}$, see appendix section A.2 |
| $w(z_i)$ | The fitness of individual $i$ with fitness function $w(\cdot)$, see appendix section A.2 |

**Table 1:** Notation, continued

| Symbol | Usage |
|---:|---|
| $\mathbf{Q}$ | The sample standardized variance-covariance matrix, (16) |
| $Q_{t,s}$ | The elements of the observed sample matrix $\mathbf{Q}$, (16) |
| $\mathbf{\Sigma}$ | The standardized variance-covariance matrix, based on our theoretic expressions |
| $\Sigma_{t,s}$ | The elements of the standardized variance-covariance matrix, (10), (11) |
| $\Delta p_{n,t}$ | The allele frequency change at site $n$ between times time $t+1$ and $t$, (16) |
| $\tau$ | The number of allele frequency changes observed, e.g. after sampling for $\tau+1$ timepoints |
| $V_{a,ssh_n}(t)$ | The additive genic variation at time $s$ as approximated by the observed decay in the sum of site heterozygosity at neutral sites, (15) |
| $\text{Var}_i(z_i)$ | The variance in trait values taken over individuals, (17) |
| $\widehat{V_A(1)}$ | The method-of-moments estimate of the additive genetic variance in the first generation, (19), (18) |
| $\widehat{F}$ | The method-of-moments estimate of Wright's standardized variance, $F = 1/2N$, (19), (18) |
| $\widehat{N}$ | The method-of-moments estimate of drift-effective population size, $N = 1/2F$ |
| $\sigma^2$ | The sampling noise around each element of the sample variance-covariance matrix. |
| $B$ | The total number of windows after partitioning the genome, (22) |
| $w_{textBP}$ | Width of windows (in basepairs), (22) |
| $v_A(1)$ | The average additive genetic variance per basepair |
| $w_{\text{CBP}}$ | Number of coding basepairs in a window, (22) |
| $W_{\text{CBP}}$ | Total number of coding basepairs in the genome, (23) |
| $G$ | A conservative measure of the total variance in allele frequency change due to linked selection, (24) |
| $G'$ | An alterate, less conservative measure of the total variance in allele frequnecy change due to linked selection, (25) |
| $G_{abs}$ | A variant of $G$ using the absolute value of covariances, (27) |

#### A.1 Decomposition of Allele Frequency Change

This decomposition of *neutral* allele frequency change between two consecutive generations is based on that of Santiago and Caballero (1995). We imagine a closed Wright–Fisher population of $N$ diploids, where each diploid $i$ contributes $k_i \sim \text{Multinom}(f_i/N, 2N)$ gametes to the next generation. We assume the population size is constant, such that one diploid begets one diploid and thus $\mathbb{E}_i(f_i) = 1$. The neutral allele frequency in the next generation can be thought of as each of the $N$ parents passing their average genotype $x_i/2$ (where $x_i \in \{0, 1, 2\}$ is the number of tracked neutral alleles individual $i$ carries) to their $k_i$ gametes, plus a random Mendelian deviation $b_{ij} \in \{0, -1/2, 1/2\}$ to each offspring $j$. Then, the frequency in the next generation can be written as

$$p_1 = \frac{1}{N} \sum_{i=1}^{N} \left( k_i \frac{x_i}{2} + \sum_{j=1}^{k_i} b_{ij} \right) \tag{28}$$

where $b_{ij} = \delta_{x_i,1}(1/2 - B_j)$, $B_j \sim \text{Bernoulli}(1/2)$ and $\delta_{x_i,1}$ is an indicator function that is one when the individual $i$ is a heterozygote (i.e. $x_i = 1$), and zero otherwise.

If we further decompose the number of offspring of individual $i$ into the genetic and non-genetic contributions, $k_i = f_i + d_i$, then

$$\begin{aligned} p_1 &= \frac{1}{N} \sum_{i=1}^{N} \left( (f_i + d_i) \frac{x_i}{2} + \sum_{j=1}^{k_i} b_{ij} \right) \\ &= \frac{1}{2N} \sum_{i=1}^{N} f_i x_i + \frac{1}{2N} \sum_{i=1}^{N} d_i x_i + \frac{1}{N} \sum_{i=1}^{N} \sum_{j=1}^{k_i} b_{ij} \end{aligned} \tag{29}$$

and the change of the neutral allele's frequency is the difference $\Delta p = p_1 - p_0$ where $p_0 = 1/2N \sum_{i=1}^{N} x_i$. Then,

$$\begin{aligned} \Delta p = p_1 - p_0 &= \frac{1}{2N} \sum_{i=1}^{N} f_i x_i + \frac{1}{2N} \sum_{i=1}^{N} d_i x_i + \frac{1}{N} \sum_{i=1}^{N} \sum_{j=1}^{k_i} b_{ij} - \frac{1}{2N} \sum_{i=1}^{N} x_i \\ &= \underbrace{\frac{1}{2N} \sum_{i=1}^{N} x_i (f_i - 1)}_{\Delta_H p_1} + \underbrace{\frac{1}{2N} \sum_{i=1}^{N} d_i x_i}_{\Delta_N p_1} + \underbrace{\frac{1}{N} \sum_{i=1}^{N} \sum_{j=1}^{k_i} b_{ij}}_{\Delta_M p_1} . \end{aligned} \tag{30}$$

These $d$'s broadly capture non-heritable variation in an individual's offspring number, with $\mathbb{E}[d_i] = 0$. In a quantitative genetics framework $\text{Var}_i(d_i)$ can include non-genetic variation in the lifetime reproductive success of individuals ($V_E$), in population genetic models $\text{Var}_i(d_i)$ can accommodate the sampling of parents to form the next generation, e.g. multinomial sampling of individuals from fitnesses (Santiago and Caballero 1995). From now forwards, we assume that variation in these $d$'s is non-heritable. We note that in non-panmictic populations, chance covariances could be created between a neutral polymorphism and environmental component of their phenotype, especially as a population expands its range into new environments that affect the phenotype, and variants "surf" to higher frequencies (Edmonds et al. 2004; Excoffier and Ray 2008; Hallatschek and Nelson 2008).

Note that by construction, the allele frequency change components $\Delta_N p_t$ $\Delta_M p_t$, and $\Delta_H p_t$ are orthogonal within each individual, given the neutral allele frequency $x$. Partitioning the allele frequency change for an individual $i$ into its components,

$$\Delta p_{t,i} = \underbrace{\frac{1}{2}x_i(f_i - 1)}_{\Delta_H p_{t,i}} + \underbrace{\frac{1}{2}x_i d_i}_{\Delta_N p_{t,i}} + \underbrace{\delta_{x_i,1}({}^{k_i}\!/\!{}_2 - M(k_i))}_{\Delta_M p_{t,i}} \tag{31}$$

$$= \frac{1}{2}x_i(f_i - 1) + \frac{1}{2}x_i d_i + \delta_{x_i,1}({}^{f_i}\!/\!{}_2 - M(f_i)) + \delta_{x_i,1}({}^{d_i}\!/\!{}_2 - M(d_i)) \tag{32}$$

where $M(n) = \sum_{j=1}^{n} B_j \sim \text{Binom}(n, 1/2)$.

Two random variables $X, Y$ are uncorrelated if $\text{Cov}(X, Y) = \mathbb{E}(XY) - \mathbb{E}(X)\mathbb{E}(Y) = 0$, and are orthogonal if either has an expected value of zero such that $\mathbb{E}(XY) = 0$. We show briefly that taking expectations *over conceptual evolutionary replicates*, the terms are orthogonal. First, the terms $x(f - 1)$ and $xd$ are orthogonal (dropping $i$ subscripts),

$$\begin{aligned} 1/4\,\text{Cov}(x(f-1), xd) &= 1/4(\mathbb{E}(x^2 d(f-1)) - \mathbb{E}(x(f-1))\mathbb{E}(xd)) \\ &= 1/4(\mathbb{E}(x^2)\mathbb{E}(d)\mathbb{E}(f-1) - \mathbb{E}(x)\mathbb{E}(f-1)\mathbb{E}(xd)) \\ &= 0 \end{aligned} \tag{33}$$

since $\mathbb{E}(f) = 1$ across evolutionary replicates due to the assumption that population size is constant, and $x \perp f$, as across all evolutionary replicates, there is no dependence between a particular neutral allele an individual carries and their fitness (though in particular replicates, such associations occur). Similarly, for the case $x = 1$ (other cases are all zero, and can be ignored), it can be shown using the law of total expectation that $\text{Cov}(xd, {}^{d}\!/\!{}_2 - M(d)) = 0$ (and likewise with $x(f - 1)$ and ${}^{f}\!/\!{}_2 - M(f)$). Note that across individuals within a population, there are weak covariances in their number of offspring as the total number of offspring must sum to $N$; under a Multinomial offspring distribution, these are of order $1/N$.

#### A.2 Temporal variance and autocovariance under multilocus selection

We assume the phenotype of an individual $i$ has an additive polygenic basis, such that their breeding value is $z_i = \sum_{l=1}^{L} \alpha_{t,l} g_{i,l}$ which deviates around a mean of zero, and $\alpha_{t,l}$ is the additive effect size at locus $l$ in generation $t$, and $g_{i,l} \in \{0, 1, 2\}$ is individual $i$'s allele count at this locus (note that effect of non-heritable environmental noise affecting the trait is accounted for in the $d_i$ terms above). We impose directional selection on this trait using an exponential fitness function, such that individual $i$'s fitness is $f_i = {}^{w(z_i)}\!/\!{}_{\bar{w}} \approx e^{z_i}$ (assuming $\bar{w} \approx 1$). If we assume individuals' phenotypic values do not deviate too far from their mean value of zero, we can approximate $f_i$ as: $f_i \approx 1 + \sum_{l=1}^{L} \alpha_{t,l} g_{i,l}$. Then, we can write the change in neutral allele frequency due to only heritable variation in fitness ($\Delta_H p_t$) as a covariance between fitness and the neutral allele frequency across individuals in generation $t$,

$$\begin{aligned}
\Delta_H p_t &= \frac{1}{2N}\sum_{i=1}^{N} x_i\,(f_i - 1) &(34)\\
&= \frac{1}{2}\operatorname{Cov}_i(x_i, f_i) &(35)\\
&= \frac{1}{2}\operatorname{Cov}_i(x_i, \sum_{l=1}^{L}\alpha_{t,l} g_{i,l}) &(36)
\end{aligned}$$

which is the is the Robertson-Price covariance (Lynch, Walsh, et al. 1998; Price 1970; Robertson 1966; Walsh and Lynch 2018).

Now, we break up the genotypic value $x_i$ into the contributions of each of the two gametes that formed individual $i$, $x_i = x_i' + x_i''$, and likewise with the trait locus $g_{i,l} = g_{i,l}' + g_{i,l}''$, where $x_i', x_i'', g_i'$, and $g_i''$ are all indicator variables. Expanding out the covariances, we have

$$\begin{aligned}
\Delta_H p_t &= \frac{1}{2}\operatorname{Cov}(x_i' + x_i'', \sum_{l=1}^{L}\alpha_{t,l}(g_{i,l}' + g_{i,l}''))\\
&= \frac{1}{2}\left(\operatorname{Cov}(x_i', \sum_{l=1}^{L}\alpha_{t,l}(g_{i,l}' + g_{i,l}'')) + \operatorname{Cov}(x_i'', \sum_{l=1}^{L}\alpha_{t,l}(g_{i,l}' + g_{i,l}''))\right)\\
&= \frac{1}{2}\left(\operatorname{Cov}(x_i', \sum_{l=1}^{L}\alpha_{t,l} g_{i,l}') + \operatorname{Cov}(x_i', \sum_{l=1}^{L}\alpha_{t,l} g_{i,l}'') + \operatorname{Cov}(x_i'', \sum_{l=1}^{L}\alpha_{t,l} g_{i,l}') + \operatorname{Cov}(x_i'', \sum_{l=1}^{L}\alpha_{t,l} g_{i,l}'')\right).
\end{aligned} \tag{37}$$

Each of these covariances is between the neutral allele and a selected allele, either on the same gamete (either maternal or paternal) or across gametes. These covariances can be written as linkage disequilibrium terms,

$$\begin{aligned}
\Delta_H p_t &= \frac{1}{2}\left(\operatorname{Cov}(x_i', \sum_{l=1}^{L}\alpha_{t,l} g_{i,l}') + \operatorname{Cov}(x_i', \sum_{l=1}^{L}\alpha_{t,l} g_{i,l}'') + \operatorname{Cov}(x_i'', \sum_{l=1}^{L}\alpha_{t,l} g_{i,l}') + \operatorname{Cov}(x_i'', \sum_{l=1}^{L}\alpha_{t,l} g_{i,l}'')\right)\\
&= \frac{1}{2}\left(\sum_{l=1}^{L}\alpha_{t,l}\operatorname{Cov}(x_i', g_{i,l}') + \sum_{l=1}^{L}\alpha_{t,l}\operatorname{Cov}(x_i', g_{i,l}'') + \sum_{l=1}^{L}\alpha_{t,l}\operatorname{Cov}(x_i'', g_{i,l}') + \sum_{l=1}^{L}\alpha_{t,l}\operatorname{Cov}(x_i'', g_{i,l}'')\right)\\
&= \frac{1}{2}\left(\sum_{l=1}^{L}\alpha_{t,l} D_l' + \sum_{l=1}^{L}\alpha_{t,l} D_l'' + \sum_{l=1}^{L}\alpha_{t,l} D_l'' + \sum_{l=1}^{L}\alpha_{t,l} D_l'\right)\\
&= \sum_{l=1}^{L}\alpha_{t,l} D_l' + \sum_{l=1}^{L}\alpha_{t,l} D_l''
\end{aligned} \tag{38}$$

where $D_L'$ is the linkage disequilibrium between alleles on the same gamete (the gametic LD), and the $D_l''$ is the across-gamete LD (non-gametic LD); see Weir (1996), p. 121.

We ignore non-gametic linkage disequilibria $D_l''$ as these are weak under random mating, and write the multilocus temporal covariance between the allele frequency changes $\Delta p_t$ and $\Delta p_s$ as

$$\begin{aligned}\text{Cov}(\Delta p_t, \Delta p_s) &= \mathbb{E}(\Delta p_t \Delta p_s) - \mathbb{E}(\Delta p_t)\mathbb{E}(\Delta p_s) \\ &= \mathbb{E}\left(\sum_{l=1}^{L} \alpha_{t,l} D'_{t,l} \sum_{l=1}^{L} \alpha_{s,l} D'_{s,l}\right) \\ &= \underbrace{\sum_{l=1}^{L} \alpha_{t,l}\alpha_{s,l}\mathbb{E}(D'_{t,l}D'_{s,l})}_{\text{persistence of association to selected site } l} + \underbrace{\sum_{l\neq k} \alpha_{t,k}\alpha_{s,l}\mathbb{E}(D'_{t,k}D'_{s,l})}_{\text{cross-associations between two selected sites } k \text{ and } l}\end{aligned} \tag{39}$$

$$(p_s^{(1)} - p_s^{(2)}) = (p_t^{(1)} - p_t^{(2)})(1 - r_l)^{s-t}. \tag{41}$$

Then, we can use this to describe the dynamics of $D_{s,l}$ to generation $t$,

$$\begin{aligned}
\frac{p_{s,l}(1-p_{s,l})}{p_{s,l}(1-p_{s,l})}(p_s^{(1)} - p_s^{(2)}) &= \frac{p_{t,l}(1-p_{t,l})}{p_{t,l}(1-p_{t,l})}(p_t^{(1)} - p_t^{(2)})(1-r)^{s-t} \\
\frac{D_{s,l}}{p_{s,l}(1-p_{s,l})} &= \frac{D_{t,l}}{p_{t,l}(1-p_{t,l})}(1-r)^{s-t} \\
D_{s,l} &= D_{t,l}\frac{p_{s,l}(1-p_{s,l})}{p_{t,l}(1-p_{t,l})}(1-r_l)^{s-t} \qquad (42)
\end{aligned}$$

(c.f. Stephan et al. 2006, equations (30) and (31)). Now, we can find the expected product $\mathbb{E}(D_{t,l}D_{s,l})$ by multiplying both sides by $D_{t,l}$ and taking expectations. We treat the allele frequency trajectory as deterministic, giving us,

$$\begin{aligned}
D_{t,l}D_{s,l} &= D_{t,l}^2\frac{p_{s,l}(1-p_{s,l})}{p_{t,l}(1-p_{t,l})}(1-r_l)^{s-t} \\
\mathbb{E}(D_{t,l}D_{s,l}) &= \mathbb{E}(D_{t,l}^2)\frac{p_{s,l}(1-p_{s,l})}{p_{t,l}(1-p_{t,l})}(1-r_l)^{s-t}. \qquad (43)
\end{aligned}$$

Then, we simplify this by replacing $\mathbb{E}(D_{t,l}^2)$ with $\mathbb{E}(\mathcal{R}_{t,l}^2)p_{t,l}(1-p_{t,l})p_t(1-p_t)$ (where $\mathbb{E}(\mathcal{R}_{t,l})$ is the square of the correlation between the neutral site and selected site $l$ at time $t$; Hill and Robertson 1968),

$$\begin{aligned}
\mathbb{E}(D_{t,l}D_{s,l}) &= \mathbb{E}(D_{t,l}^2)\frac{p_{s,l}(1-p_{s,l})}{p_{t,l}(1-p_{t,l})}(1-r_l)^{s-t} \\
&= \mathbb{E}(\mathcal{R}_{t,l}^2)p_t(1-p_t)p_{t,l}(1-p_{t,l})\frac{p_{s,l}(1-p_{s,l})}{p_{t,l}(1-p_{t,l})}(1-r_l)^{s-t} \\
&= \mathbb{E}(\mathcal{R}_{t,l}^2)p_t(1-p_t)p_{s,l}(1-p_{s,l})(1-r_l)^{s-t}. \qquad (44)
\end{aligned}$$

Returning to Equation (39) and replacing the $\mathbb{E}(D_{t,l}D_{s,l})$ terms with our expression above,

$$\mathrm{Cov}(\Delta p_t, \Delta p_s) = \sum_{l=1}^{L} \alpha_{t,l}\alpha_{s,l}\mathbb{E}(\mathcal{R}_{t,l}^2)p_t(1-p_t)p_{s,l}(1-p_{s,l})(1-r_l)^{s-t} \qquad (45)$$

again ignoring the cross-associations between selected sites. Dividing our temporal covariance by the neutral site's $p_t(1-p_t)$, we can write the multilocus temporal covariance in a standardized form (analogous to Wright's F),

$$\frac{\mathrm{Cov}(\Delta p_t, \Delta p_s)}{p_t(1-p_t)} = \sum_{l=1}^{L} \alpha_{t,l}\alpha_{s,l}p_{s,l}(1-p_{s,l})\mathbb{E}(\mathcal{R}_{t,l}^2)(1-r_l)^{s-t}. \qquad (46)$$

**Using average additive genetic variation** We can approximate Equation (46) by noticing that the terms $\alpha_{t,l}\alpha_{s,l}p_{s,l}(1-p_{s,l})$ are similar to an additive genic variation if effect sizes remain constant through time. We make that assumption here, writing $\alpha_l := \alpha_{t,l} = \alpha_{s,l}$, leading to

$$\frac{\mathrm{Cov}(\Delta p_t, \Delta p_s)}{p_t(1-p_t)} = \sum_{l=1}^{L} \alpha_l^2 p_{s,l}(1-p_{s,l})\mathbb{E}(\mathcal{R}_{t,l}^2)(1-r_l)^{s-t}. \tag{47}$$

We can further simplify this by assuming that there is no covariance between the additive genic variation at a selected site, and the LD between that selected site and the neutral site. We write the additive genic variation at site $l$ at time $s$ as $v_{a,l}(s) = 2\alpha_l^2 p_{s,l}(1-p_{s,l})$, and the average additive genic variation across loci as $\overline{v_a(s)} := {}^{V_a(s)}/_{L} = \frac{1}{L}\sum_l 2\alpha_l^2 p_{s,l}(1-p_{s,l})$. Then, each locus's additive genic variation can be expressed as: $v_{a,l}(s) = \overline{v_a(s)} + \varepsilon_l$. Substituting this, the autocovariance is

$$\begin{aligned}\frac{\mathrm{Cov}(\Delta p_t, \Delta p_s)}{p_t(1-p_t)} &= \frac{1}{2}\sum_{l=1}^{L} v_{a,l(s)}\mathbb{E}(\mathcal{R}_{t,l}^2)(1-r_l)^{s-t} \\ &= \frac{1}{2}\sum_{l=1}^{L}(\overline{v_a(s)} + \varepsilon_l)\mathbb{E}(\mathcal{R}_{t,l}^2)(1-r_l)^{s-t} \\ &= \frac{1}{2} \underbrace{\overline{v_{a(s)}}}_{\substack{\text{average genic} \\ \text{variation per locus}}} \times \underbrace{\left(\sum_{l=1}^{L}\mathbb{E}(\mathcal{R}_{t,l}^2)(1-r_l)^{s-t}\right)}_{\text{sum of persistence associations}} + \underbrace{\frac{1}{2}\left(\sum_{l=1}^{L}\varepsilon_l\mathbb{E}(\mathcal{R}_{t,l}^2)(1-r_l)^{s-t}\right)}_{\text{effect-association covariation}}.\end{aligned} \tag{48}$$

We assume that this last term, which is non-zero in expectation only if there is covariance between the additive genic variation at a selected site and the expected LD between the selected and the neutral sites is zero. Rewriting the total genic variation as $V_a(s)$,

$$\frac{\mathrm{Cov}(\Delta p_t, \Delta p_s)}{p_t(1-p_t)} = \frac{V_a(s)}{2L}\sum_{l=1}^{L}\mathbb{E}(\mathcal{R}_{t,l}^2)(1-r_l)^{s-t}. \tag{49}$$

**Continuous approximation to chromosomes** Currently, we have treated the positions of the selected sites are fixed, conditioning on knowing that a selected site $l$ has a recombination fraction $r_l$ away from the focal neutral site site. Here, we make a few further assumptions. First, we assume the $L$ selected loci are each independently and identically uniformly distributed along a *continuous* region of $R$ Morgans in length ($g \sim U(-{}^{R}/_{2}, {}^{R}/_{2})$), and we now calculate the covariance at a focal neutral site at the origin by taking expectations over the random positions of these sites. Second, we assume that the LD between the neutral and selected site $l$ only depends on the recombination fraction between the sites. Since $g$ is the genetic distance, the recombination fraction is now provided by a mapping function $r(g)$ that maps genetic distances to recombination fractions. Throughout the paper and in the simulations, we use Haldane's mapping function, $r(g) = \frac{1}{2}(1-e^{-2|g|})$ (note the absolute value translates positions on $[-{}^{R}/_{2}, {}^{R}/_{2}]$ to distances to the focal neutral site). Next, we assume the LD between two sites can be completely determined by the distance between the focal neutral site and a random selected site, allowing us to rewrite $\mathbb{E}(\mathcal{R}_{t,l}^2)$ as the function $\mathbb{E}(\mathcal{R}_t^2(r(g)))$. Now, letting $\mathbb{E}_{r_l}(\cdot)$ represent the expectation taken over the random positions of selected sites on the genetic map,

$$\frac{\text{Cov}(\Delta p_t, \Delta p_s)}{p_t(1-p_t)} = \frac{V_a(s)}{2L} \sum_{l=1}^{L} \mathbb{E}_{r_l} \left( \mathbb{E}(\mathcal{R}_t^2(r_l))(1-r_l)^{s-t} \right) \tag{50a}$$

$$= \frac{V_a(s)}{2L} \sum_{l=1}^{L} \int_{-R/2}^{R/2} \mathbb{E}(\mathcal{R}_t^2(r(g)))(1-r(g))^{s-t} \frac{1}{R} dg \tag{50b}$$

$$= \frac{V_a(s)}{2R} \int_{-R/2}^{R/2} \mathbb{E}(\mathcal{R}_t^2(r(g)))(1-r(g))^{s-t} dg \tag{50c}$$

since the $1/L$ cancels with the $L$ from the sum of expectations.

As our trait is neutrally evolving before directional selection starts we use the expected neutral LD, $\mathbb{E}(\mathcal{R}^2) = (10+\rho)/(22+13\rho+\rho^2)$ where $\rho = 4Nr(g)$ (Hill and Robertson 1968; Ohta and Kimura 1969) when $t$ is the first generation selection begins.

#### A.3 Averaging covariance across multiple loci

Thus far, our covariance assumes that a single neutral site is positioned in the center of a region $R$ Morgans long, with selected sites uniformly distributed along this region. In our simulations, however, we simulate a region that contains many neutral sites which we average over in calculating the temporal autocovariance. In this case, we average over the random distance between a neutral site's position $n$ and a selected site's position $g$, which is $c = |n-g|$, where $n, g \sim U(0, R)$. This random variable $c$ is distributed according to the triangle distribution, $f(c) = 2(R-c)/R^2$; we replace the uniform PDF in Equation (50b) with the triangle density PDF and average over the distance between sites,

$$\frac{\mathbb{E}_n(\text{Cov}(\Delta p_t, \Delta p_s))}{\mathbb{E}_n(p_t(1-p_t))} = \frac{V_a(s)}{2} \int_0^R \mathbb{E}(\mathcal{R}_t^2(r(c)))\ (1-r(c))^{(s-t)} \frac{2(R-c)}{R^2}\, dc \tag{51}$$

$$= \frac{V_a(s)}{2} \mathcal{A}(R,t,s) \tag{52}$$

where $\mathbb{E}_n(\cdot)$ indicates we take the expectation also over neutral sites, and we use $\mathcal{A}(R,t,s)$ to denote the average linkage disequilibrium between selected and neutral sites that persists from generations $t$ to $s$ ($t \leq s$). Note that in calculating the standardized covariance above, we use a ratio of expectations rather than the expectation of the ratio (Bhatia et al. 2013).

#### A.4 Empirically Calculating the Average LD Persisting Across Generations

In the previous expressions for temporal autocovariance, we stepped through a conceptual model for the average levels of linkage disequilibria between neutral and selected sites that persists across $|s-t|$ generations ($\mathcal{A}(R,t,s)$), where the positions of selected and neutral sites are randomly distributed along a chromosomal region. In systems with a known recombination map and studies where linkage disequilibria can be calculated, we have the recombination fraction $r_{i,j}$ and the pairwise linkage disequilibria $\mathbf{R}^2_{i,j}$ between between two loci $i$ and $j$ (where $\mathbf{R}^2$ is the $M \times M$ matrix of pairwise LD calculated at time $t$). Since we do not *a priori* know whether a site is selected or not, we sum over all polymorphic $M$ loci, thus characterizing the average linkage disequilibria in a region as

$$\overline{\mathcal{A}(t,s)} = \frac{2}{M(M-1)} \sum_{i=1}^{M} \sum_{j>i} \mathbf{R}^2_{i,j} (1 - r_{i,j})^{|t-s|}. \tag{53}$$

This sum is the empirical analog to the integral in Equation (10).

#### A.5 The Strength of Unlinked and Non-gametic Associations

Here, we characterize the contribution of completely genetically unlinked loci segregating for fitness variation to the change in frequency of our neutral allele. Across evolutionary replicates, there is no expected covariance between the neutral allele an individual carries and their fitness ($\mathbb{E}(\Delta_H p_t) = \mathbb{E}(\mathrm{Cov}(x_i, f_i)) = 0$); rather, for unlinked loci, chance associations are created from the variance around this sampling process of neutral alleles into individuals with varying fitness ($\mathrm{Var}(\Delta_H p_t) = \mathrm{Var}(\mathrm{Cov}(x_i, f_i))$). As the neutral allele and fitness variation independently assort themselves into individuals, the chance associations that form have a variance given by $\mathrm{Var}(\mathrm{Cov}_i(x_i, f_i))$. This has the form of the sampling variance of a covariance, which for random variables $X$ and $Y$ is given by Kendall et al. (1994, p. 472),

$$\mathrm{Var}(\mathrm{Cov}(X,Y)) = \frac{(n-1)^2}{n^3}(\mu_{22} - \mu_{11}^2) + \frac{n-1}{n^3}(\mu_{20}\mu_{02} - \mu_{11}^2) \tag{54}$$

$$\text{where } \mu_{rs} = \mathbb{E}(X - \mu_X)^r (Y - \mu_Y)^s) \tag{55}$$

where $\mu_X$ and $\mu_Y$ are the means of $X$ and $Y$ respectively, *and the variance is taken over conceptual replicate populations and the covariance is calculated over the individuals in a population*. Then, applying this to our covariance $\Delta_H p_1 = \mathrm{Cov}_i(x_i, f_i)$,

$$\begin{aligned}
\mathrm{Var}(\Delta_H p_1) &= {}^1\!/\!_4 \,\mathrm{Var}(\mathrm{Cov}_i(x_i, f_i)) \\
&= \frac{(N-1)^2}{4N^3} \left( \mathbb{E}[(x_i - p_0)^2 (f_i - 1)^2] - \mathbb{E}[(x_i - p_0)(f_i - 1)]^2 \right) + \\
&\quad \frac{N-1}{4N^3} \left( \mathbb{E}[(x_i - p_0)^2] \mathbb{E}[(f_i - 1)^2] - \mathbb{E}[(x_i - p_0)(f_i - 1)]^2 \right) \\
&= \frac{(N-1)}{4N^3} \mathrm{Var}(x_i) \,\mathrm{Var}(f_i) \\
&\approx \frac{p_0(1-p_0)}{2N} \mathrm{Var}(f_i).
\end{aligned} \tag{56}$$

Thus the chance covariances that form between the neutral alleles individuals carry and their fitness have a variance proportional to ${}^{\mathrm{Var}(f_i)}\!/\!_{2N}$.

**Non-gametic linkage disequilibrium's contribution to temporal autocovariance** Throughout the paper, we ignore the effects of non-gametic linkage disequilibria, $D''_{t,l}$, the disequilibria that

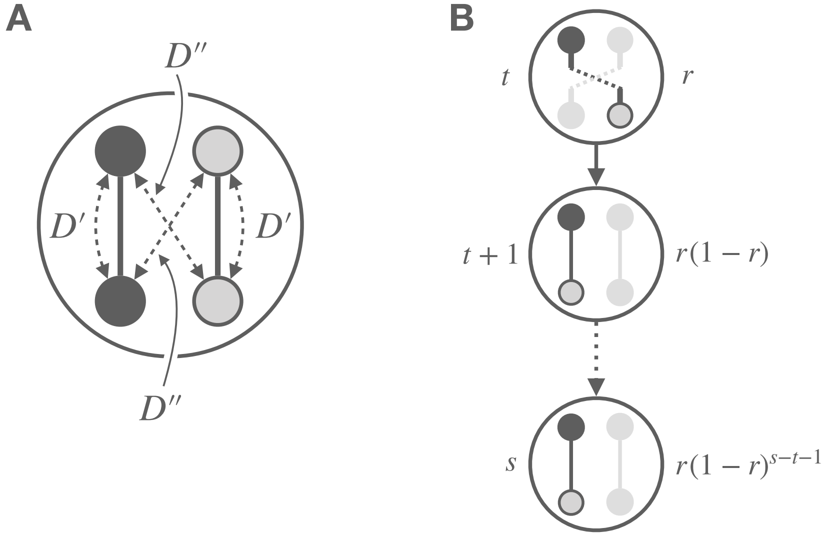


**Figure 8:** A: An illustration of gametic ($D'$) and non-gametic ($D''$) linkage disequilibria between two loci in a diploid. B: An illustration of how non-gametic linkage disequilibria in generation $t$ is converted to gametic linkage disequilibria through recombination (which happens with probability $r$, the recombination fraction between the two loci), and is then maintained until generation $s$ with probability $(1-r)^{s-t-1}$. The gray loci on the gray gamete indicate the homologous, but not tracked focal association. Overall, the covariance created by the conversion of non-gametic LD to gametic LD is $\mathbb{E}(D'_s D''_t) = r(1-r)^{s-t-1}\mathbb{E}\left((D''_t)^2\right)$.

occurs between the two gametes (maternal and paternal) at two loci (see Figure 8 (A) for an illustration of gametic LD $D'$ and non-gametic LD $D''$). Following Weir's equation for the sampling variance of non-gametic LD $D_{A/B}$ (p. 124, 1996), the chance non-gametic disequilibrium that builds up sampling $2N$ gametes into $N$ individuals is,

$$\mathbb{E}\left((D''_{t,l})^2\right) = \mathrm{Var}(D_{A/B}) = \frac{1}{2N} p_A(1-p_A)p_B(1-p_B). \tag{57}$$

Note the similar form to Equation (56) as both the unlinked and non-gametic LD arise from the random sampling of alleles at different loci into individuals.

There is no expected covariance between our gametic and non-gametic LD within a generation $\mathbb{E}(D''_{t,l}, D'_{t,l}) = 0$, assuming random mating. However, $\mathbb{E}(D''_{t,l}, D'_{s,l}) > 0$ for $s > t$, as a fraction of the non-gametic LD may be converted into gametic LD in the next generation. Specifically, following Santiago and Caballero (1995), we can write the product of the non-gametic LD in generation $t$ with the gametic LD in generation $s$ as

$$\mathbb{E}(D''_{t,l}, D'_{s,l}) = r(1-r)^{(s-t-1)}\mathbb{E}\left((D''_{t,l})^2\right) \tag{58}$$

where a proportion $r$ the non-gametic LD in generation $t$ is converted into gametic LD, and a proportion $(1-r)^{(s-t-1)}$ of this is carried forward unbroken by recombination over the remaining $s-t-1$ generations (see Figure 8B for an illustration of this process).

$$\begin{aligned}\text{Var}(p_3 - p_0) = \mathbb{E}\Bigg(\Big(&\underbrace{S_1 + D_1 + H_1}_{\Delta p_1} + \\ &\underbrace{(1-r)GS_1 + S_2 + D_2 + H_2}_{\Delta p_2} + \\ &\underbrace{(1-r)^2G^2S_1 + (1-r)GS_2 + S_3 + D_3 + H_3}_{\Delta p_3}\Big)^2\Bigg). \end{aligned} \tag{61}$$

Grouping terms by the generation that the initial association was formed, we see how Santiago and Caballero (1995) define the $Q_i$ terms in their notation,

$$\begin{aligned}\text{Var}(p_3 - p_0) = \mathbb{E}\Bigg(\Big(&\underbrace{S_1(1 + (1-r)G + (1-r)^2G^2)}_{\text{creation and persistence of generation 1 associations } := S_1Q_3} + D_1 + H_1 + \\ &\underbrace{S_2(1 + (1-r)G)}_{\text{creation and persistence of generation 2 associations } := S_2Q_2} + D_2 + H_2 + \\ &\underbrace{S_3}_{\text{creation of generation 3 associations } := S_3Q_1} + D_3 + H_3\Big)^2\Bigg). \end{aligned} \tag{62}$$

since the associations created in generation $i$ ($0 < i \leq t$) persist with probability $(1-r)^{t-i}$, with proportion $G^{t-i}$ of its original fitness variation in generation $t$. In general, the cumulative impact of the associations formed $i$ generations ago has coefficient $Q_i = \sum_{j=0}^{i-1}(1-r)^jG^j$. Using these $Q_i$ terms simplifies this equation to

$$\begin{aligned}\text{Var}(p_4 - p_0) = \mathbb{E}\Big(\Big(&S_1Q_4 + D_1 + H_1 + S_2Q_3 + D_2 + H_2 + \\ &S_3Q_2 + D_3 + H_3 + S_4Q_1 + D_4 + H_4\Big)^2\Big) \end{aligned} \tag{63}$$

or, in general,

$$\text{Var}(p_t - p_0) = \sum_{i=1}^{t} \mathbb{E}(D_i^2) + \mathbb{E}(H_i^2) + Q_{t-i+1}^2 E(S_i^2). \tag{64}$$

Then, Santiago and Caballero (1995) note that assuming $V_n$, $C^2$, and population size $N$ are constant across generations, the magnitude of all of the effects $\mathbb{E}(D_i^2)$, $\mathbb{E}(H_i^2)$, and $\mathbb{E}(S_i^2)$ are constant across all generations (for all $i$, so we omit the $i$ subscript for these terms), except for a geometric decay due to drift at a rate $(1 - 1/2N_e)$ per generation that effects all terms. Such that, when we include the decay in the variance due to drift,

$$\operatorname{Var}(p_t - p_0) = \sum_{i=1}^{t} \left(\mathbb{E}(D^2) + \mathbb{E}(H^2) + Q_i^2 E(S^2)\right) \left(1 - \frac{1}{2N_e}\right)^{t-i} \tag{65}$$

(cf. p. 1018 of Santiago and Caballero 1995). In the long run, the variance in the neutral allele's frequency change hits a balance. Many copies of the neutral allele segregating in the population are on fitness backgrounds it has recently become stochastically associated with, as segregation and recombination have not broke these associations apart. A few copies of the neutral allele are on fitness backgrounds they became associated with many generations ago, that have by chance survived to remain associated. In all cases, the effect these associations have on present-day allele frequency change is weakened by the fact that natural selection has reduced the genetic variance of these fitness backgrounds. Since the long-run variance in allele frequency under drift in a Wright–Fisher population is

$$\operatorname{Var}(p_t) = p_0(1 - p_0)\left[1 - \left(1 - \frac{1}{2N}\right)^t\right]. \tag{66}$$

One can estimate the effective population size $N_e$ using the observed difference in variances $\operatorname{Var}(p_t)$ and $\operatorname{Var}(p_{t-1})$. (Note that this is a different long-run effective population size used by others, $N_e = p_0(1 - p_0)t/(2\operatorname{Var}(p_t - p_0))$, Crow and Kimura 1970) Santiago and Caballero use Equation (66), taking the difference $\operatorname{Var}(p_t) - \operatorname{Var}(p_{t-1})$ and rearranging to end up with the large $t$ estimator of $N_e$,

$$N_e = \frac{p_0(1 - p_0) - \operatorname{Var}(p_{t-1})}{2(\operatorname{Var}(p_t) - \operatorname{Var}(p_{t-1}))} \tag{67}$$

(cf. p. 1018, Santiago and Caballero 1995). Rearranging,

$$\begin{aligned} 2N_e(\operatorname{Var}(p_t) - \operatorname{Var}(p_{t-1})) &= p_0(1 - p_0) - \operatorname{Var}(p_{t-1}) \\ 2N_e \operatorname{Var}(p_t) - 2N_e \operatorname{Var}(p_{t-1}) + \operatorname{Var}(p_{t-1}) &= p_0(1 - p_0) \\ 2N_e\left(\operatorname{Var}(p_t) - \operatorname{Var}(p_{t-1})\left(1 - \frac{1}{2N_e}\right)\right) &= p_0(1 - p_0). \end{aligned} \tag{68}$$

This very conveniently simplifies the sum in Equation (65), as we can show with the case of $t = 3$,

$$
\begin{aligned}
\mathrm{Var}(p_3) &= \left(\mathbb{E}(D^2) + \mathbb{E}(H^2) + Q_1^2\mathbb{E}(S^2)\right)\left(1 - \frac{1}{2N_e}\right)^2 + \\
&\quad \left(\mathbb{E}(D^2) + \mathbb{E}(H^2) + Q_2^2\mathbb{E}(S^2)\right)\left(1 - \frac{1}{2N_e}\right) + \\
&\quad \mathbb{E}(D^2) + \mathbb{E}(H^2) + Q_3^2\mathbb{E}(S^2) \\
\mathrm{Var}(p_2)\left(1 - \frac{1}{2N_e}\right) &= \left(\mathbb{E}(D^2) + \mathbb{E}(H^2) + Q_1^2\mathbb{E}(S^2)\right)\left(1 - \frac{1}{2N_e}\right)^2 + \\
&\quad \left(\mathbb{E}(D^2) + \mathbb{E}(H^2) + Q_2^2\mathbb{E}(S^2)\right)\left(1 - \frac{1}{2N_e}\right) \\
\mathrm{Var}(p_3) - \mathrm{Var}(p_2)\left(1 - \frac{1}{2N_e}\right) &= \mathbb{E}(D^2) + \mathbb{E}(H^2) + Q_3^2\mathbb{E}(S^2)
\end{aligned}
$$

or, generally,

$$
\mathrm{Var}(p_t) - \mathrm{Var}(p_{t-1})\left(1 - \frac{1}{2N_e}\right) = \mathbb{E}(D^2) + \mathbb{E}(H^2) + Q_t^2\mathbb{E}(S^2) \tag{69}
$$

Inserting this into Equation (68), the long-run effective population size can be written as

$$
\begin{aligned}
N_e &= \frac{p_0(1-p_0)}{2\left(\mathrm{Var}(p_t) - \mathrm{Var}(p_{t-1})\left(1 - \frac{1}{2N_e}\right)\right)} \\
&= \frac{p_0(1-p_0)}{2\left(\mathbb{E}(D^2) + \mathbb{E}(H^2) + Q_\infty^2\mathbb{E}(S^2)\right)}
\end{aligned} \tag{70}
$$

where $Q_\infty = 1 + 1/2 + 1/4 + \ldots = 2$ in Robertson's (1961) model, and $Q_\infty = 1/(1 - G(1-r))$ in Santiago and Caballero's (1995) model. (Note: $r$ here represents the recombination fraction between fitness variation and neutral sites, which differs from Santiago and Caballero (1995) equation 17, where $r$ represents the correlation between parental fitness).

Then, Santiago and Caballero (1995) show,

$$
\mathbb{E}(S^2) = \frac{p_0(1-p_0)}{2N}C^2 \tag{71}
$$

$$
\mathbb{E}(D^2) = \frac{p_0(1-p_0)}{2N}\frac{V_n}{4} \tag{72}
$$

$$
\mathbb{E}(H^2) = \frac{p_0(1-p_0)}{2N}\frac{1}{2} \tag{73}
$$

(cf. Santiago and Caballero 1995, equation 11). Inserting these into Equation (70), we have

$$
N_e = \frac{4N}{2 + V_n + 4Q_\infty^2 C^2} \tag{74}
$$

which is a simplified version of Santiago and Caballero's equation 16, and which further simplifies to $N_e = N$ when $V_n = 2$ and $C^2 = 0$, the effective population size of a Wright–Fisher population of $N$ hermaphroditic individuals.

**The covariances caused by fitness associations** With an understanding now of the basics of Santiago and Caballero's (1995, 1998) models, how they connect to our notation, and how they reach their expression for the long-run effective population size, we turn now to finding the temporal autocovariances implicit in their model. We start by looking at the variance in allele frequency between generations 0 and 4 (Equation (61)), including an additional generation so the pattern is clearer later,

$$\begin{aligned}\mathrm{Var}(p_4 - p_0) = \mathbb{E}\Bigg(\Bigg(&\underbrace{S_1 + D_1 + H_1}_{\Delta p_1} + \\ &\underbrace{(1-r)GS_1 + S_2 + D_2 + H_2}_{\Delta p_2} + \\ &\underbrace{(1-r)^2G^2S_1 + (1-r)GS_2 + S_3 + D_3 + H_3}_{\Delta p_3} + \\ &\underbrace{(1-r)^3G^3S_1 + (1-r)^2G^2S_2 + (1-r)GS_3 + S_4 + D_4 + H_4}_{\Delta p_4}\Bigg)^2\Bigg).\end{aligned}$$

The cross-terms like $\mathbb{E}(D_1D_2)$, $\mathbb{E}(H_1D_1)$ and $\mathbb{E}(S_1S_2)$ are all expected products of independent random variables, where the expectation of each random variable is zero, and consequently are all zero. The only non-zero cross terms are products of $\mathbb{E}(S_i^2)$. When we look at the covariances with the allele frequency change in the initial generation and a later generation $s$, $\mathrm{Cov}(\Delta_H p_1, \Delta_H p_s)$,

$$\mathrm{Cov}(\Delta_H p_1, \Delta_H p_1) = \mathrm{Var}(\Delta_H p_1) = \mathbb{E}(S_1^2) \tag{75}$$

$$\mathrm{Cov}(\Delta_H p_1, \Delta_H p_2) = \frac{\mathbb{E}(S_1^2)G}{2}(1-r) \tag{76}$$

$$\mathrm{Cov}(\Delta_H p_1, \Delta_H p_3) = \frac{\mathbb{E}(S_1^2)G^2}{2}(1-r)^2 \tag{77}$$

$$\mathrm{Cov}(\Delta_H p_1, \Delta_H p_4) = \frac{\mathbb{E}(S_1^2)G^3}{2}(1-r)^3 \tag{78}$$

where the $1/2$ coefficient comes from the fact that in Santiago and Caballero's work, the $\mathbb{E}(S_1^2)$ products represent *both* the $\mathrm{Cov}(\Delta_H p_t, \Delta_H p_s)$ and $\mathrm{Cov}(\Delta_H p_s, \Delta_H p_t)$ terms, so a single temporal autocovariance in our notation is half their joint covariance term.

However, if our reference generation is different, say 2, the associations from earlier generations that have persisted to that generation can also lead to covariances to later generations. Looking at the covariances

$$\mathrm{Cov}(\Delta_H p_2, \Delta_H p_2) = \frac{\mathbb{E}(S_2^2) + \mathbb{E}(S_1^2)(1-r)G}{2} \tag{79}$$

$$\mathrm{Cov}(\Delta_H p_2, \Delta_H p_3) = \frac{\mathbb{E}(S_2^2)G + \mathbb{E}(S_1^2)(1-r)G^2}{2}(1-r) \tag{80}$$

$$\mathrm{Cov}(\Delta_H p_2, \Delta_H p_4) = \frac{\mathbb{E}(S_2^2)(1-r)G^2 + \mathbb{E}(S_1^2)(1-r)^2G^3}{2}(1-r)2. \tag{81}$$

Likewise, the covariances $\text{Cov}(\Delta_H p_3, \Delta_H p_s)$ include the associations that persist from earlier generations. In general,

$$\text{Cov}(\Delta_H p_t, \Delta_H p_s) = \frac{1}{2} \sum_{i=1}^{t} \mathbb{E}(S_i^2) \left(G(1-r)\right)^{t+s-2i}, \quad \text{for } \; t \leq s. \tag{82}$$

This expression is more complex than our expression for temporal autocovariance because its modeling the LD in generation $t$ as it builds up from generations 1 to $t$. In contrast, our expressions for covariance incorporate all of this build up of LD as the initial LD term $\mathbb{E}(\mathcal{R}_t)$ (for a single locus case). This expression for autocovariance implied by Santiago and Caballero's (1995) work matches our expression for autocovariance (for a single locus) when the arbitrary first generation is $t$ rather than 1

$$\text{Cov}(\Delta_H p_t, \Delta_H p_s) = \frac{\mathbb{E}(S_t^2) G^{s-t}}{2} (1-r)^{s-t}, \quad \text{for } \; t \leq s. \tag{83}$$

Using the expression for $\mathbb{E}(S_t^2)$ (equivalent to $\text{Var}(\Delta_H p_t)$ in our notation) derived in Appendix Section A.5,

$$\frac{\text{Cov}(\Delta_H p_t, \Delta_H p_s)}{p_t(1-p_t)} = \frac{C^2 G^{s-t}}{4N} (1-r)^{s-t}, \quad \text{for } \; t \leq s. \tag{84}$$

This is analogous to Equation (8) for a single locus, where $C^2 G^{s-t}$ is the additive variation in generation $s$ (equivalent to our $V_a(s)$), and the factor $1/2N$ represents the chance build up of LD between the neutral site and an unlinked fitness background. In our expression, we condition on existing linkage disequilibrium $\mathbb{E}(\mathcal{R}_t^2)$ between the neutral site and its fitness background, whereas they assume a buildup of linkage disequilibria to a drift-recombination equilibrium. We can see this by returning to the $\Delta_H p_4$ term of Equation (61),

$$\begin{aligned} \text{Var}(\Delta_H p_4) &= \mathbb{E}\left((1-r)^3 G^3 S_1 + (1-r)^2 G^2 S_2 + (1-r) G S_3 + S_4 + D_4 + H_4\right)^2 && (85) \\ &= (1-r)^5 G^5 \mathbb{E}(S_1^2) + (1-r)^4 G^4 \mathbb{E}(S_2^2) + (1-r)^2 G^2 \mathbb{E}(S_3)^2 + \mathbb{E}(S_4^2) && (86) \end{aligned}$$

where following Santiago and Caballero's (1995, p. 1018) approach, we can replace each $\mathbb{E}(S_i)$ with $\mathbb{E}(S_i^2) = \mathbb{E}(S^2)(1 - 1/2N)^{i-1}$ and let $G = 1$ as we focus on the buildup of LD. This gives us the general equation,

$$\text{Var}(\Delta_H p_t) = \mathbb{E}(S^2) \sum_{i=1}^{t} (1-r)^{2i} \left(1 - \frac{1}{2N}\right)^{i-1} \tag{87}$$

and taking this geometric series to infinity converges (since $(1-r)^2(1 - 1/2N) < 1$) and replacing $\mathbb{E}(S^2)$ with the chance associations that build up gametes sampled into individuals (Equation (56)),

$$\begin{aligned}
\mathbb{E}((\Delta_H p_\infty)^2) &= \mathbb{E}(S^2)\sum_{i=1}^{\infty}(1-r)^{2i}\left(1-\frac{1}{2N}\right)^{i-1} \\
&= \frac{\mathbb{E}(S^2)}{1-1/2N}\sum_{i=1}^{\infty}\left((1-r)^2\left(1-\frac{1}{2N}\right)\right)^{i} \\
&= \frac{C^2 p_0(1-p_0)}{1-1/2N}\frac{1}{1-(1-r)^2(1-1/2N)}.
\end{aligned} \tag{88}$$

When we assume $r \to 0$ and $1/N \to 0$ and $Nr$ is a constant, this gives us

$$\mathbb{E}(\mathcal{R}^2) = \frac{\mathbb{E}((\Delta_H p_\infty)^2)}{C^2 p_0(1-p_0)} \approx \frac{1}{1+4Nr} \tag{89}$$

which is analogous to the $\mathbb{E}(\mathcal{R}^2)$ measure of linkage disequilibrium, standardized to rescale the fitness variation. The right-hand side is identical to Sved's identity by descent equilibrium $\mathbb{E}(\mathcal{R}^2)$ under drift-recombination balance (1971). Note that our expression can be recovered from Equation (82) when the reference generation $t \to \infty$ such that the LD hits its equilibrium level,

$$\frac{\mathrm{Cov}(\Delta_H p_t, \Delta_H p_s)}{p_0(1-p_0)} = \frac{1}{2}\underbrace{C^2 G^{s-t}}_{V_A(s)}\underbrace{\frac{1}{1+4Nr}}_{\mathbb{E}(\mathcal{R}_t^2)}(1-r)^{s-t} \tag{90}$$

which is identical to our expression for temporal autocovariance when initial LD is due to a neutral drift-recombination balance, and the change in $V_A$ during selection is modeled as a geometric decay at rate $G$. Note that the terms in underbraces indicate the corresponding terms in Equation (8) for a single locus.

#### A.7 Multilocus simulation details

**Targeting an initial level of Additive Genetic Variation** We choose $\theta$ for the coalescent simulations to target a total number of segregating sites $L+M$, where $L$ is the number of selected sites (a parameter we vary in our multilocus simulations), and at least $M = 200$ randomly placed neutral sites over which we can calculate the temporal autocovariance. Then, the total number of target sites $M+L$ is then inflated by a factor of 1.5 to ensure a sufficient number of sites given the random mutation process. The $L$ selected sites are randomly chosen from the segregating sites, and all remaining mutations are neutral. Thus, using Watterson's expression for the expected number of segregating sites under the coalescent (1975), we have $\theta = 1.5(M+L)/(\gamma + \log(2N))$ where $\gamma \approx 0.577$ is Euler's Gamma. Each of the $L$ selected sites is given a random effect size of $\pm\alpha$ with equal probability, where we choose by targeting a specific level additive genic variation $V_a = 2\alpha^2\sum_l^L p_l(1-p_l) = \alpha^2 SSH_L$ where $SSH_L = 2\sum_l^L p_l(1-p_l)$ is the sum of site heterozygosity of the $L$ neutrally evolving sites that will be selected once selection begins. Under neutrality, $\mathbb{E}(SSH_L) = \theta_L = L/(\gamma+\log(2N))$. Then, we set $\alpha = \sqrt{V_a/\theta_L}$. We empirically validate that our target genic variation is close to the empirically observed level.

#### A.8 Accounting for allele frequency sampling noise

*In practice, one will calculate the temporal variance-covariance matrix on allele frequency trajectories calculated from sampled chromosomes from the population. We assume a binomial sampling process, where $n$ chromosomes are sampled from the population such that $\widetilde{p} = X/n$, and $X \sim Binom(p, n)$. We can then write $\widetilde{p}_t = p_t + \varepsilon_t$, and our covariances can be written as*

$$\begin{aligned}\mathrm{Cov}(\Delta\widetilde{p}_t, \Delta\widetilde{p}_s) &= \mathrm{Cov}(\widetilde{p}_{t+1} - \widetilde{p}_t, \widetilde{p}_{s+1} - \widetilde{p}_s) \\ &= \mathrm{Cov}(p_{t+1} + \varepsilon_{t+1} - p_t - \varepsilon_t, p_{s+1} + \varepsilon_{s+1} - p_s - \varepsilon_s).\end{aligned} \tag{91}$$

*Note this simplifies to*

$$\mathrm{Cov}(\Delta\widetilde{p}_t, \Delta\widetilde{p}_s) = \mathrm{Cov}(p_{t+1} - p_t, p_{s+1} - p_s) \;\text{ if }\; |t - s| > 1, \tag{92}$$

*since in these cases, the sampling noise at a timepoint is not shared between the estimated allele frequency changes. However, if $|t - s| = 1$ the sampling noise from timepoint $t + 1$ is shared, biasing the sample estimate of covariance:*

$$\begin{aligned}\mathrm{Cov}(\Delta\widetilde{p}_t, \Delta\widetilde{p}_{t+1}) &= \mathrm{Cov}(p_{t+1} + \varepsilon_{t+1} - p_t - \varepsilon_t, p_{t+2} + \varepsilon_{t+2} - p_{t+1} - \varepsilon_{t+1}) \\ &= \mathrm{Cov}(\Delta p_t + \varepsilon_{t+1} - \varepsilon_t, \Delta p_{t+1} + \varepsilon_{t+2} - \varepsilon_{t+1}) \\ &= \mathrm{Cov}(\Delta p_t, \Delta p_{t+1}) - \mathrm{Var}(\varepsilon_{t+1}). \end{aligned} \tag{93}$$

*Similarly, the variance ($t = s$) is biased, as it is impacted by the binomial sampling noise too,*

$$\begin{aligned}\mathrm{Var}(\Delta\widetilde{p}_t) &= \mathrm{Cov}(\Delta p_t + \varepsilon_{t+1} - \varepsilon_t, \Delta p_t + \varepsilon_{t+1} - \varepsilon_t) \\ &= \mathrm{Var}(\Delta p_t) + \mathrm{Var}(\varepsilon_{t+1}) + \mathrm{Var}(\varepsilon_t) \end{aligned} \tag{94}$$

*In practice, these covariances are calculated over loci in a region or across the entire genome. We assume that the tracked allele has been randomly swapped (e.g. the tracked allele frequency is not systematically the minor, major, or reference allele), such that $\mathbb{E}(\Delta p_{t,l}) = 0$ for all $t$ and $l$. Then, the unbiased covariance estimate as calculated over loci is*

$$\frac{1}{L}\sum_{l=1}^{L}(\Delta p_{t,l}\Delta p_{t+1,l}) = \frac{1}{L}\sum_{l=1}^{L}\Delta\widetilde{p}_{t,l}\Delta\widetilde{p}_{t+1,l} + \frac{1}{L}\sum_{l=1}^{L}\varepsilon^2_{t+1,l}. \tag{95}$$

*Then, we can use an unbiased plugin estimate of the frequency sampling variance $\mathbb{E}(\varepsilon^2_{t+1,l}) = V(\varepsilon_{t+1,l}) = {}^{p_{t+1,l}(1-p_{t+1,l})}/_{(n_{t,l}-1)}$ (Nei 1987, p. 191) to estimate these bias terms, and add or subtract them from the estimator accordingly. Accounting for finite sampling, the unbiased sample variance-covariance matrix now has elements:*

$$Q_{t,t+1} = \frac{1}{L}\sum_{l=1}^{L}\Delta\widetilde{p}_{t,l}\Delta\widetilde{p}_{t+1,l} + \frac{1}{L}\sum_{l=1}^{L}\frac{p_{t+1,l}(1-p_{t+1,l})}{n_{t+1,l}-1}, \tag{96}$$

*and variance*

$$Q_{t,t} = \frac{1}{L}\sum_{l=1}^{L}(\Delta\widetilde{p}_{t,l})^2 - \frac{1}{L}\sum_{l=1}^{L}\frac{p_{t,l}(1-p_{t,l})}{n_{t,l}-1} - \frac{1}{L}\sum_{l=1}^{L}\frac{p_{t+1,l}(1-p_{t+1,l})}{n_{t+1,l}-1}. \tag{97}$$

*In Section 2.5, we used population frequencies in introducing the method of moments estimators of $\widehat{V_A(1)}$ and $\widehat{N}$. Here, we discuss the performance of these estimators with sample allele frequencies. Our simulations are identical to those described in Section 2.4, except we have increased the target number of neutral sites in each region so it is around $10,000$. We mimic sampling of $n = \{50, 100, 200, 500\}$ chromosomes from the population, and use the bias-corrected sample variance-covariance matrix in the method-of-moments approach described in Section 3 to estimate $\widehat{V_A(1)}$ and $\widehat{N}$.*

*In Figure 1, we show the performance of our estimators in the case where $n = 100$ chromosomes have been sampled from the population. Overall, there are two important differences compared with Figure 5 of the main text. First, while the estimator $\widehat{V_A(1)}$ performs well for high levels around*

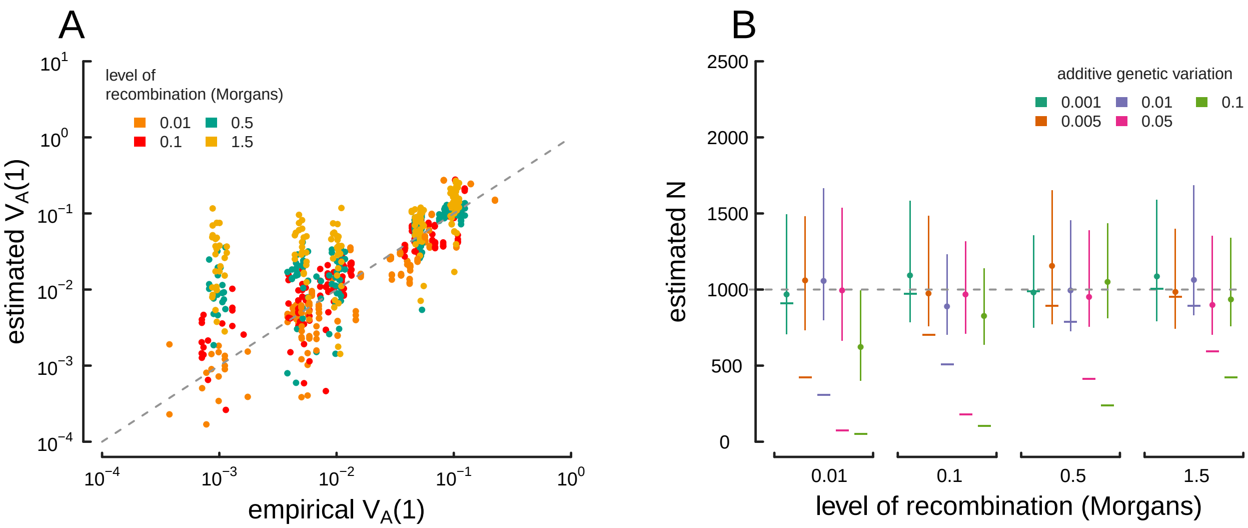


**Figure 1:** *True parameter values and estimates using the method-of-moments approach on sample ($n = 100$ chromosomes) multilocus simulation data; these figures are analogous to Figure 5 in the main text, except the estimators have been calculated on sample, rather than population frequency data. (A) The true $V_A(1)$ (x-axis) and $\widehat{V_A(1)}$ estimated from the sample variance/covariance matrix (y-axis) for each simulation replicate across different levels of recombination (indicated by each point's color). (B) Estimated drift-effective population size ($\widehat{N}$) across a range of simulations with different levels of additive genetic variance and recombination. Each point denotes the median, with lines denoting the interquartile range. A simple temporal estimate of the effective population size, estimated with accounting for the effects of selection, is averaged for each replicate and plotted as a dash. The true value ($N = 1,000$) is shown with the dashed gray line.*

*$V_A \sim 0.1$, the variance around the estimator increases significantly as $V_A$ grows weaker. As the covariances are proportional to $V_A/R$, sampling noise grows larger than the theoretic temporal autocovariance as $V_A$ become weaker. Then, one cannot discriminate against the chance covariances formed by the sampling process from the temporal autocovariance created by linked selection without either a large sample size or more timepoints. Second, the approach underestimates $V_A(1)$ for very loose linkage ($R > 1/2$). This is another consequence of the first problem; as sampling noise grows equal, or larger than the magnitude of temporal autocovariance, the estimation procedure performs poorly. As the linkage becomes more loose, the magnitude of temporal autocovariance grows weak relative to the sampling noise, and this noise can be partially absorbed by $\widehat{N}$. This effect can be somewhat ameliorated by calculating the sample variance-covariance matrix over a shorter region of the genome such that $R$ is smaller (as long as SNP density is sufficient), or by increasing the sample size. Finally, the estimation of effective population size shown in Figure 1 are also affected by $V_A$, as discussed in Section 3 (though the effect is obscured by the sampling noise): high levels of $V_A$ in regions with low recombination generally lead to underestimates of $\widehat{N}$.*

*To understand how sample size affects the method-of-moments estimators, Figure 2 depicts median relative error of $\widehat{V_A(1)}$ and $\widehat{N}$ for various sample sizes.*

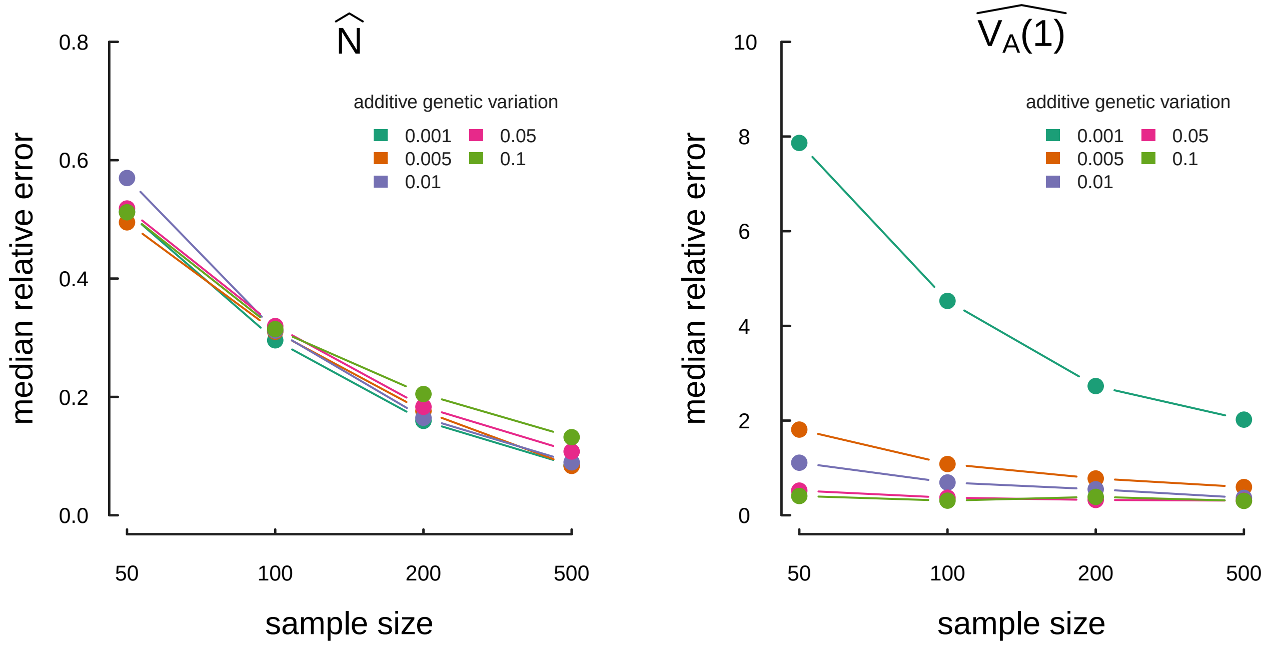


**Figure 2:** The median relative estimation error, over 30 replicate simulations, of the method of moments estimator for drift effective population size ($\widehat{N}$) and the initial additive genetic variance ($\widehat{V_A(1)}$).

### 1563 B Appendix Figures

#### 1564 B.1 Validation of simulation routine

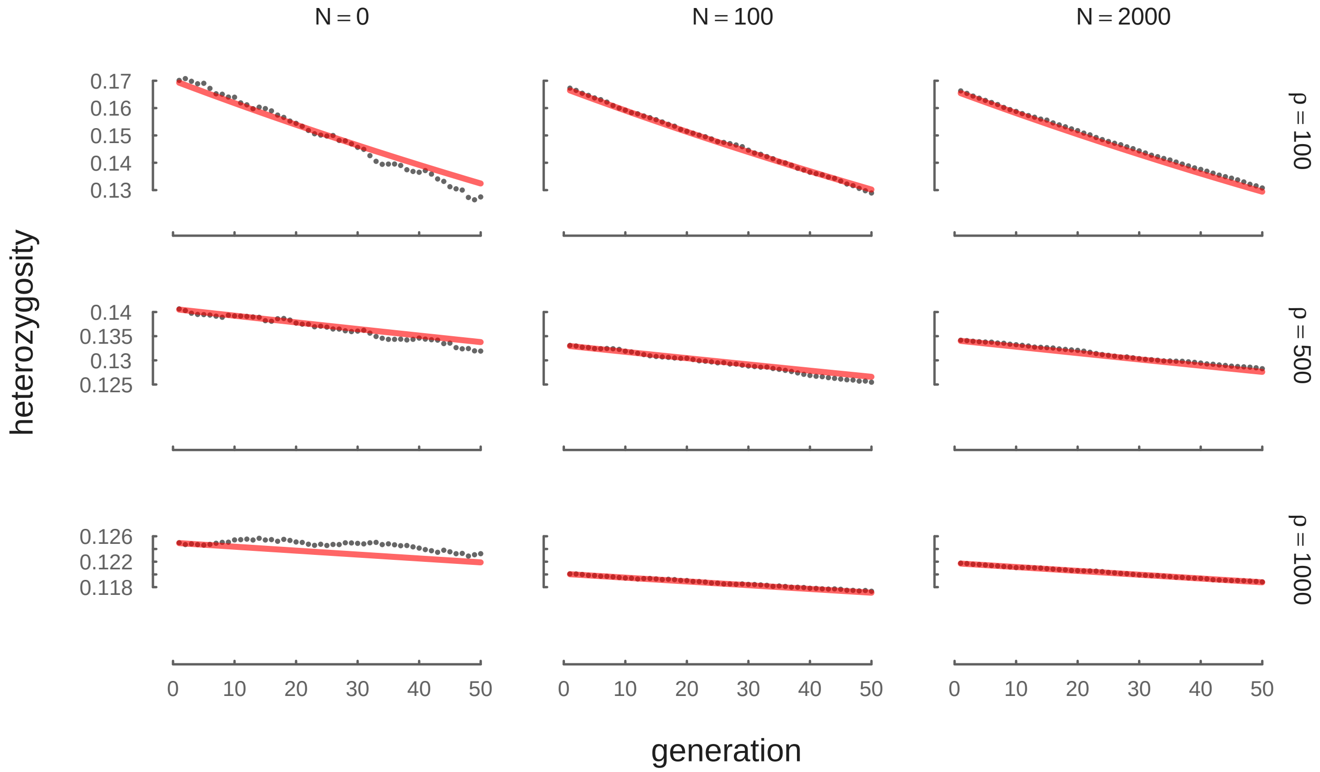


**Figure 3:** The neutral decay of heterozygosity due to drift, averaged across 100 replicates and a variety of $N$ and $\rho$ levels. The red line is the theoretic expectation $H_t = H_1(1 - 1/(2N))^{t-1}$.

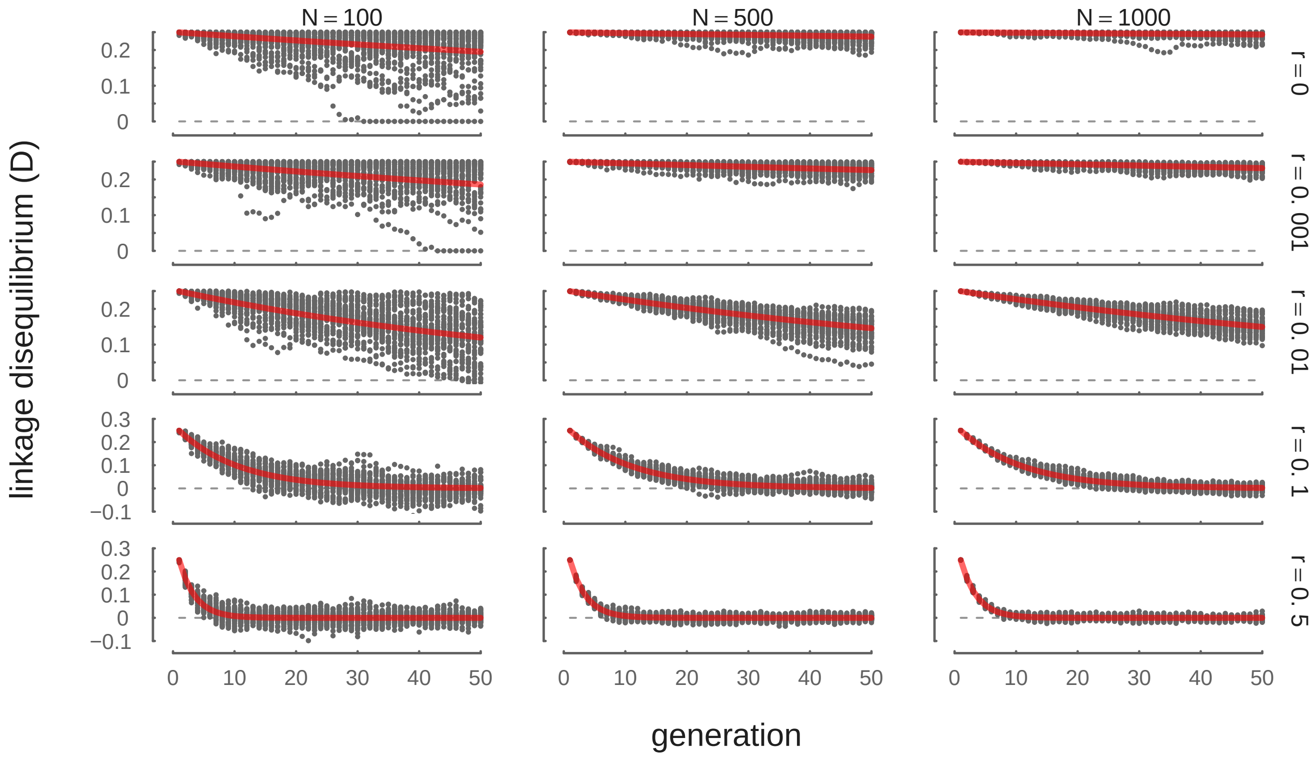


**Figure 4:** The decay of LD between neutral sites due to recombination, across 100 replicates and a variety of $V_A$ and $R$ levels. The initial population is created to have an artificial level of initial LD, with half the gametes carrying all derived alleles, and the other half carrying all ancestral alleles. The red line is the theoretic expectation $D_t = D_1(1 - 1/(2N))^{t-1}(1 - r)^{t-1}$.

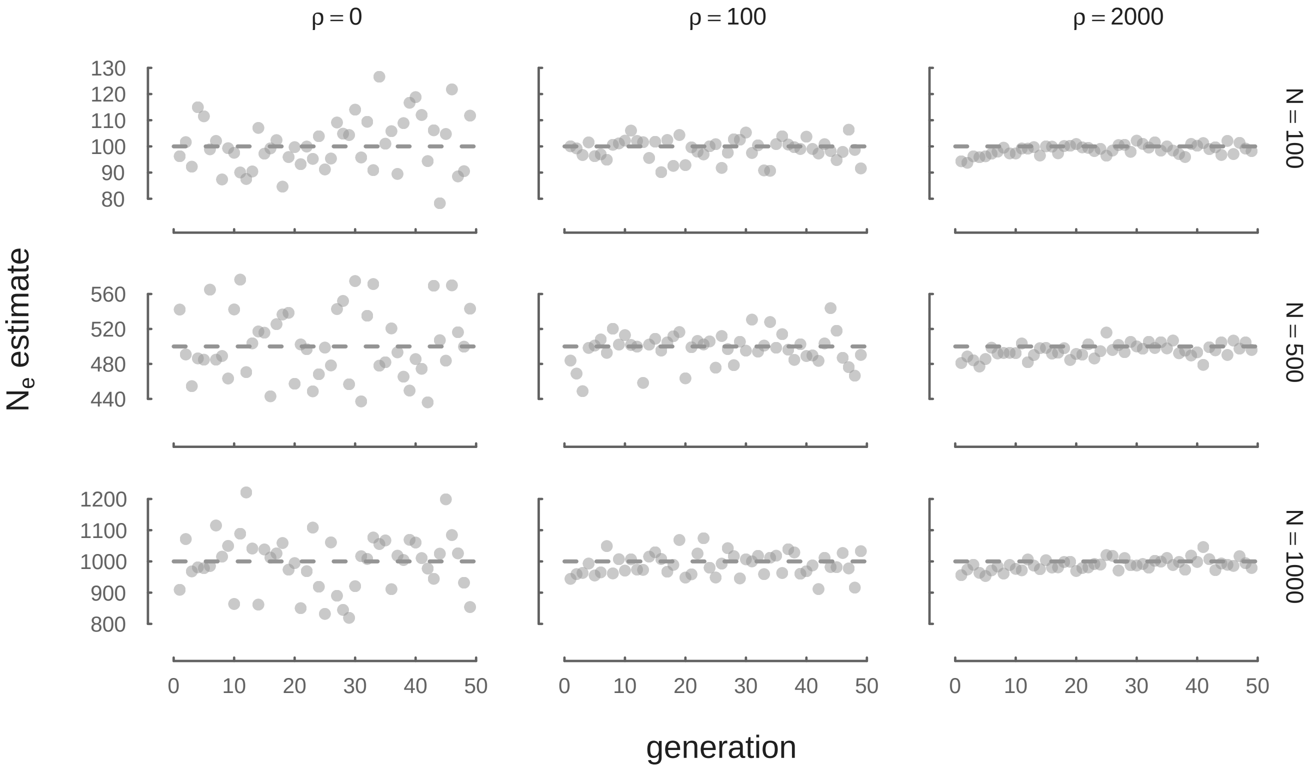


**Figure 5:** $N_e$ estimated through time from neutral forward simulations is consistent with true $N$ across a variety of recombination $\rho$ and $N$ parameters. Here, we estimate $N_e$ with $\widehat{N_e} = {p(1-p)}/{2\,\mathrm{Var}(\Delta p)}$ from neutral allele frequency changes.

#### B.2 Dynamics of Variances

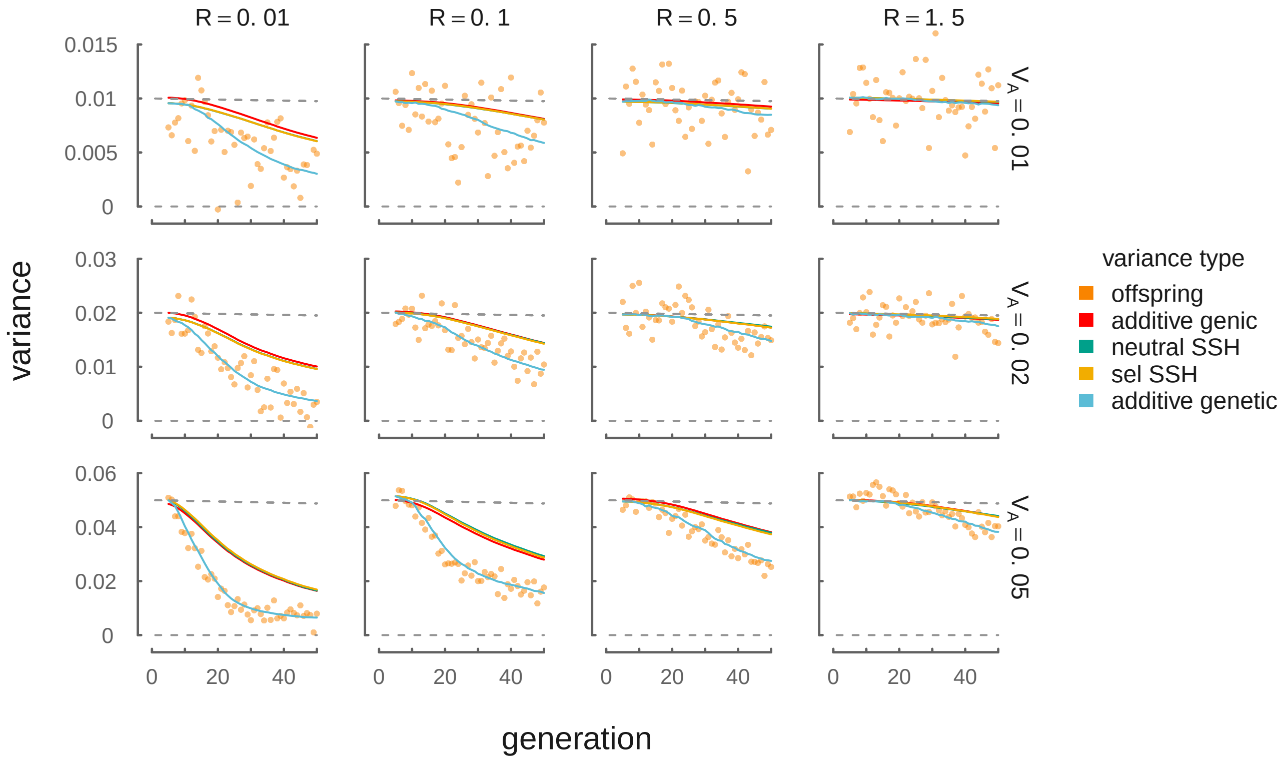


**Figure 6:** The dynamics of different variances in our simulations, across a variety of initial target $V_A$ and $R$ levels. Orange points represent the empirical heritable variance in offspring (e.g. with the noise of the Wright–Fisher reproduction process removed). These closely track the additive genetic variance for the trait undergoing directional selection, $V_A = \mathrm{Var}(z)$. The red line shows the dynamics of additive genic variance, $V_a = 2\sum_l \alpha_l^2 p_l(1-p_l)$, which is closely tracked by both the selected (yellow line) and neutral (green line) sum of site heterozygosity proxies described in Section 2.3.

1566 ## B.3 Supplementary Temporal Autocovariance Figures

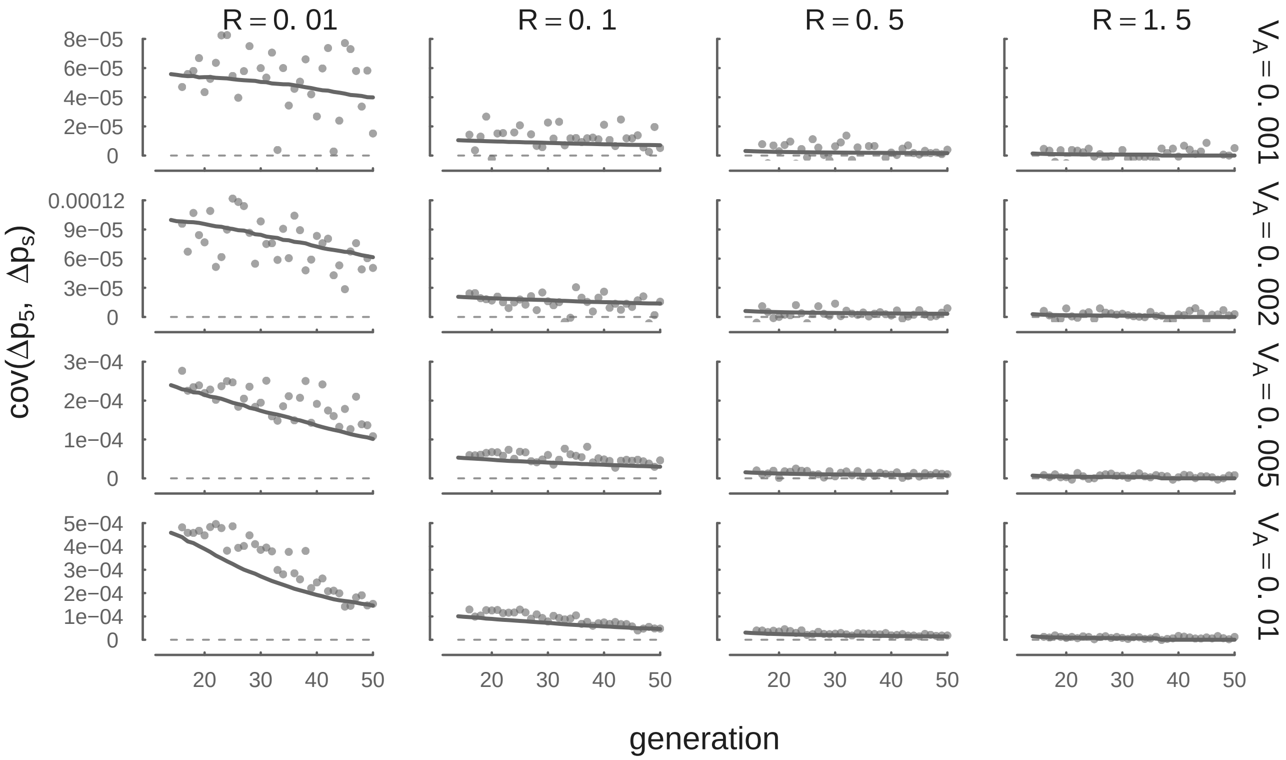


**Figure 7:** Each panel shows the temporal autocovariance $\mathrm{Cov}(\Delta p_{13}, \Delta p_s)$ on the y-axis, where $s$ varies along the x-axis. This is analogous to Figure 2 with a different reference generation ($t = 13$), and compares the averaged simulation results (points) with the temporal autocovariance predicted by Equation (10) using the empirical additive genetic variance (curve). *The covariances in these panels are weaker compared to those in Figure 2 because by generation 13, additive genetic variance for fitness and the linkage disequilibria between neutral and selected sites has decayed.*

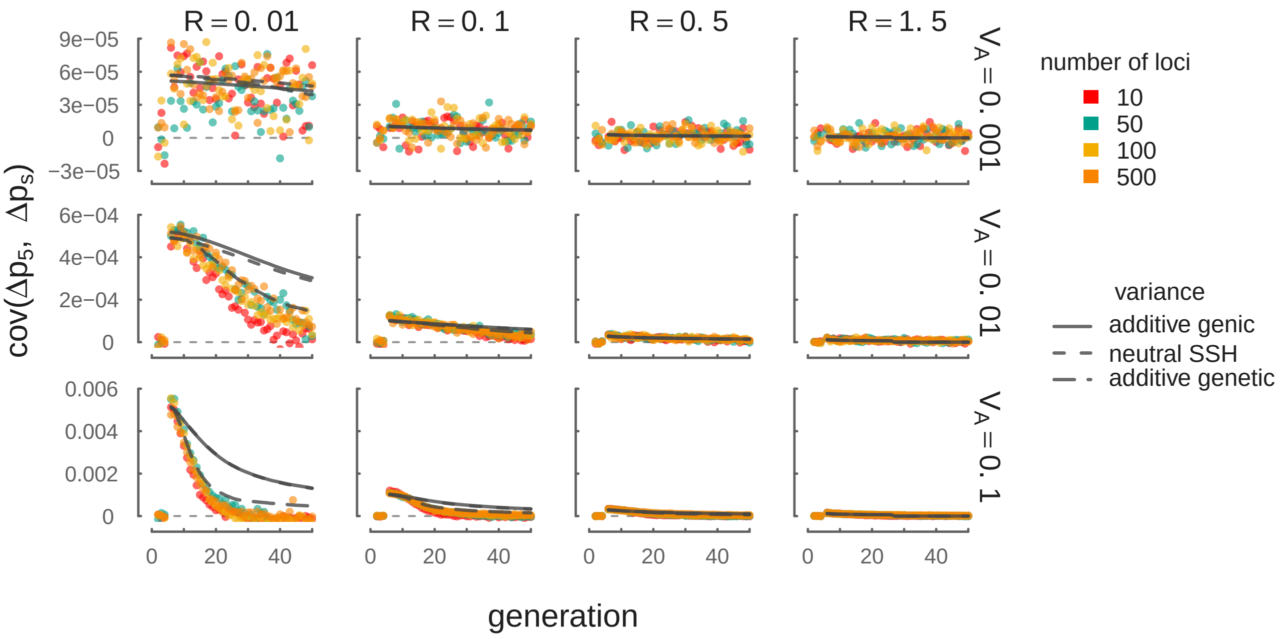


**Figure 8:** A version of Figure 2 with a subset of $V_A$ parameters used in our simulations that vary over orders of magnitude. This demonstrates that our theory using the empirical additive genic variance (solid gray line), additive genetic variance (long dashed gray line), and the neutral SSH proxy (short dashed gray lines) performs as described in Section 2.5 even when $V_A$ varies over orders of magnitude in a region. Higher variance in the empirical covariances with weak selection ($V_A = 0.001$) are due to chance covariances due to drift. The light gray dashed line depicts $\text{Cov}(\Delta p_5, \Delta p_s) = 0$.

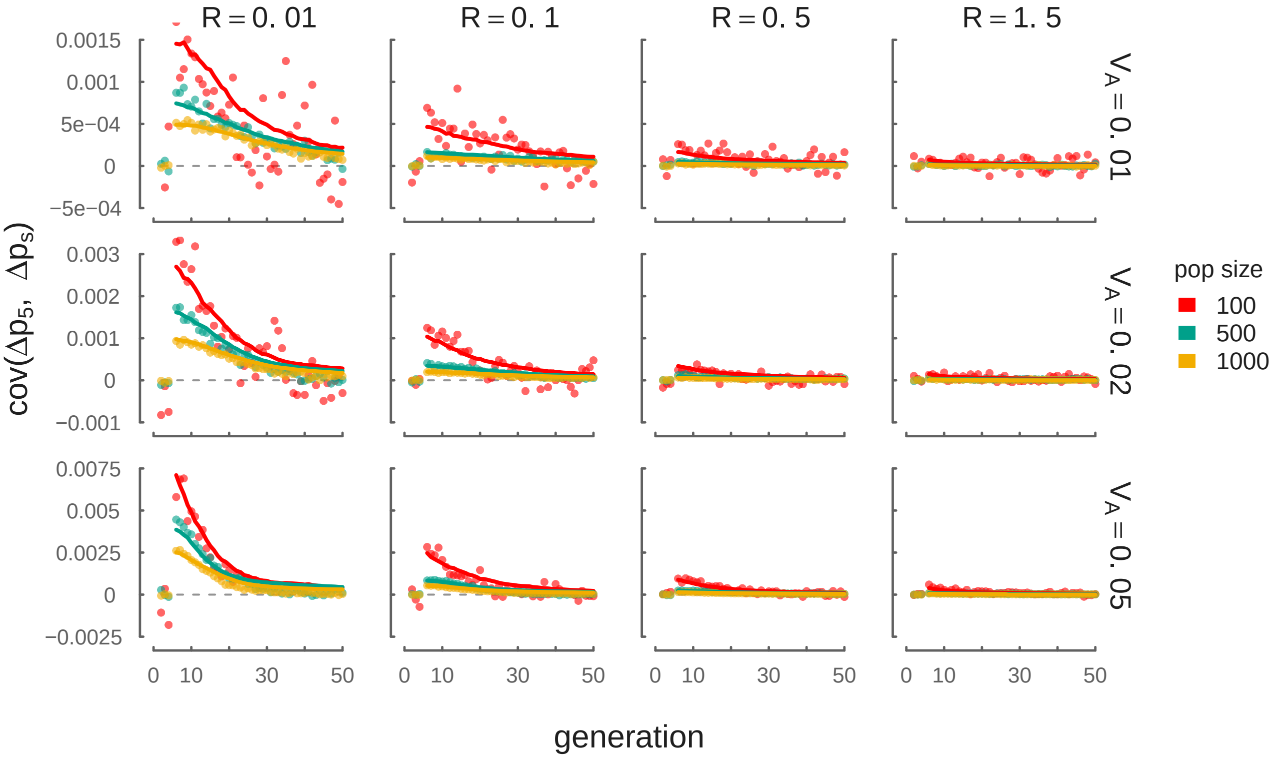


**Figure 9:** A version of Figure 2 demonstrating temporal autocovariance simulation results and theoretic predictions with varying $N$. This demonstrates that our theory using the empirical additive genetic variance (lines) fits simulations across a variety of $N$ parameters. The light gray dashed line depicts $\mathrm{Cov}(\Delta p_5, \Delta p_s) = 0$. Note that the initial LD varies due to differing equilibrium levels of LD from our burnin across varying $N$.
